# Aβ Amyloid Scaffolds the Accumulation of Matrisome and Additional Proteins in Alzheimer’s Disease

**DOI:** 10.1101/2023.11.29.568318

**Authors:** Yona Levites, Eric B. Dammer, Yong Ran, Wangchen Tsering, Duc Duong, Measho Abreha, Joshna Gadhavi, Kiara Lolo, Jorge Trejo-Lopez, Jennifer Phillips, Andrea Iturbe, Aya Erquizi, Brenda D. Moore, Danny Ryu, Aditya Natu, Kristy Dillon, Jose Torrellas, Corey Moran, Thomas Ladd, Farhana Afroz, Tariful Islam, Jaishree Jagirdar, Cory C. Funk, Max Robinson, David R. Borchelt, Nilüfer Ertekin-Taner, Jeffrey W. Kelly, Frank L. Heppner, Erik C. B. Johnson, Karen McFarland, Allan I. Levey, Stefan Prokop, Nicholas T. Seyfried, Todd E. Golde

**Affiliations:** Department of Pharmacology and Chemical Biology, School of Medicine, Emory University, Atlanta, GA, USA; Department of Biochemistry, School of Medicine, Emory University, Atlanta, GA, USA; Department of Neurology, School of Medicine, Emory University, Atlanta, GA, USA; Goizueta Brain Health Institute and Alzheimer’s Disease Research Center, Emory University School of Medicine, Atlanta, GA, USA; Center for Neurodegenerative Disease Center, Emory University School of Medicine, Atlanta, GA, USA; Department of Pathology, College of Medicine, University of Florida, Gainesville, USA; Evelyn F. and William L. McKnight Brain Institute, University of Florida, Gainesville, Fl, USA; Center for Translational Research in Neurodegenerative Disease, University of Florida, Gainesville, Fl, USA; Department of Neuroscience, College of Medicine, University of Florida, Gainesville, USA; Department of Neurology, College of Medicine, University of Florida, Gainesville, USA; Mayo Clinic, Department of Neuroscience, Jacksonville, FL, USA; Mayo Clinic, Department of Neurology, Jacksonville, FL, USA; Department of Pathology and Laboratory Medicine, Emory University Hospital. Atlanta, GA, USA; Institute for Systems Biology, Seattle, WA, USA; Department of Chemistry and The Skaggs Institute for Chemical Biology, The Scripps Research Institute, La Jolla, California, USA; Department of Neuropathology, Charité – Universitätsmedizin Berlin, corporate member of Freie Universität Berlin and Humboldt-Universität zu Berlin; German Center for Neurodegenerative Diseases (DZNE) Berlin; Cluster of Excellence, NeuroCure, Charitéplatz 110117 Berlin, Germany

**Author notes:** These authors contributed equally (Yona Levites, Eric B. Dammer).

**Keywords:** proteomics, Alzheimer’s disease, amyloid, models, plaques, aggregation

## Abstract

We report a highly significant correlation between human Alzheimer’s disease (AD) brain proteome changes and those in CRND8 APP695NL/F transgenic mice. Comparing protein changes observed in the CRND8 mice with co-expression networks derived from human Alzheimer’s disease (AD), reveals both conserved and divergent module changes. Many proteins in the most highly conserved module (M42, matrisome) accumulate in plaques, cerebrovascular amyloid (CAA), dystrophic neuronal processes, or a combination thereof. Overexpression of two M42 proteins, midkine (Mdk) and pleiotrophin (PTN), in CRND8 mice brains leads to increased accumulation of Aβ in plaques and in blood vessels; further, recombinant MDK and PTN enhance Aβ aggregation into amyloid structures. Multiple M42 proteins bind to fibrillar Aβ42 and a non-human amyloid fibril *in vitro*. Supporting this binding data, MDK and PTN co-accumulate with transthyretin (TTR) amyloid in the heart. Notably, our findings establish that proteomic changes in modules observed in human AD brains define an Aβ amyloid “responsome” that is well conserved from mouse models to humans. Further, distinct amyloid structures appear to serve as scaffolds, facilitating the co-accumulation of proteins with signaling functions, and this co-accumulation may contribute to downstream pathological sequalae. Overall, this contextualized understanding of proteomic changes and their interplay with amyloid deposition provides valuable insights into the complexity of AD pathogenesis and potential biomarkers and therapeutic targets.

## INTRODUCTION

The amyloid cascade hypothesis (ACH) serves as a foundational framework for comprehending the pathogenesis of Alzheimer’s disease (AD) ^1,2^. Recent evidence from anti-amyloid immunotherapies, developed based on this hypothesis, offers the initial therapeutic validation, affirming the central tenet that Aβ deposition plays a role in the AD pathogenic cascade ^3-5^. These antibodies target the early symptomatic phase of AD demonstrating robust reductions in brain Aβ amyloid, slowing AD decline, but not halting or reversing it. Collectively, these findings reinforce the previously proposed notion within the field that, by the time symptoms manifest, the influence of amyloid itself is limited, while additional pathologies may become self-reinforcing or irreversible ^6^.

Despite extensive investigation, significant gaps in our understanding of AD pathogenesis remain ^2,7,8^. The initial notion of a purely linear amyloid cascade, whereby Aβ amyloid deposition in sequence triggers neurofibrillary tau pathology, neuronal degeneration, immune responses, and clinical AD symptoms, is now recognized as overly simplistic. Studies utilizing proteomic, transcriptomic, metabolomic, pathological, and biomarker approaches have unveiled the intricate complexity of changes occurring over decades in the brains of individuals as various AD hallmark pathologies emerge ^7,9-24^. These investigations reveal potentially non-linear and multifaceted alterations that affect all major cell-types in the brain, ultimately resulting in regional and widespread organ failure.

At a systems-level, these pathological and cellular changes manifest as fluctuations in protein levels, transcript quantities, and metabolite concentrations, even in brain regions where significant pathology may not be apparent ^13^. This further underscores the complexity of AD pathogenesis and highlights the need to consider diverse factors contributing to disease progression beyond the core proteins implicated by the ACH or genetic studies. Comprehensive analyses incorporating multi-omic datasets such as those generated by the Accelerating Medicine Partnership for AD (AMP-AD) and other consortiums provide crucial insights into the systems pathobiology of AD, enabling a more nuanced comprehension of the disease’s dynamic processes occurring at the cellular and molecular ^7,9-24^.

Typically, systems-level omic studies alone cannot establish cause and effect relationships. However, by combining strong correlational inference with omics obtained from different AD stages, we can generate hypotheses regarding causality or the role of these changes in the disease. To determine whether these omic-level changes are a consequence of disease or play a more fundamental role in pathogenesis, it is essential to i) compare the data across related disorders and disease models, ii) integrate the data with genetic risk and biomarker information, and iii) directly manipulate the levels of the altered proteins or RNAs in various model systems to observe their impact on pathophysiology potentially relevant to human AD. By undertaking these approaches, we can gain deeper insights into the significance of omic-level changes in AD, going beyond their descriptive nature and unraveling their potential role in the disease process.

Here, we performed quantitative tandem mass tag mass spectrometry (TMT-MS) based proteomics analyses of the brains from CRND8 transgenic mice at various ages and compared these findings to human proteomic data previously generated from >500 subjects from the Religious Orders Study/Memory and Aging Project (ROSMAP) and Banner Sun Health cohorts ^9^. We find robust and highly significant correlation in global proteome changes in human AD brain compared to CRND8 transgenic mouse models. Through integration of protein changes observed in mouse models with co-expression networks derived from human AD, we resolve conserved and divergent network module changes between humans and mice. Multiple proteins that map to the M42 matrisome module, accumulate in Aβ plaques, cerebrovascular amyloid (CAA) and/or dystrophic neuronal processes. We also demonstrate that overexpressing two M42 proteins, midkine (Mdk) and pleiotrophin (PTN), in CRND8 mice leads to accelerated accumulation of Aβ, accompanied by dramatic Aβ amyloid increases in blood vessels linked to CAA. Moreover, *in vitro* experiments using recombinant MDK and PTN reveal their ability to enhance Aβ aggregation into amyloid structures. Surprisingly, we find that many M42 proteins exhibit binding to fibrillar Aβ and non-human amyloid fibrils *in vitro*. Supporting this association, both MDK and PTN co-accumulate with transthyretin (TTR) amyloid in the heart. These findings establish several critical insights. First, that specific proteomic or module changes observed in human AD brains result from amyloid deposition. Second, amyloid structures seem to serve as scaffolds, facilitating the co-accumulation of proteins such as MDK and PTN with well-established signaling functions. This comprehensive understanding of proteomic changes and their interplay with amyloid deposition provides valuable insights into the complexity of AD pathogenesis and potential biomarkers. More specifically, they suggest that antagonizing MDK and/or PTN binding to amyloid could be a therapeutic strategy.

## RESULTS

### Deep TMT MS proteomes from 6, 12and 18M CRND8 APP Mice

We have generated TMT-MS proteomic data from the 8M urea extracts of the brains of 6 months (M) 12M and 18M CRND8 mice and non-transgenic (non-Tg) controls (Figure 1, Table S1-S5). We identified 8561, 10066, and 10543 proteins in the 6M, 12M and 18M cohorts with <50% missing values. In the 6M cohort, CRND8 mice show 1097 significant differentially expressed proteins (DEPs) with 172 increased and 65 decreased by more than 20% (Figure 1A). In the 12M cohort, 2340 DEPs are detected with 454 increased and 327 decreased by more than 20% (Figure 1B). In the 18M cohort, 2244 DEPs are detected with 639 increased and 450 decreased by more than 20% (Figure 1C). Notably, proteomic changes in 12M and 18M cohorts are highly reproducible (R^2^=0.8129 p < 1e^-200^) (Figure 1D), despite being close, but not precise biological replicates. Indeed, out of the 1006 proteins that were significantly altered and detected in both cohorts only 8 were discordant and for these the log2 fold change (log2 Fc) was modest. Such data indicate that using small cohorts of CRND8 mice and matched non-Tg brains with varying levels of amyloid deposition, we can reliably and reproducibly detect proteomic alterations and that these changes increase with age in this transgenic mouse model of rapid and robust Aβ deposition.

**Figure 1.**
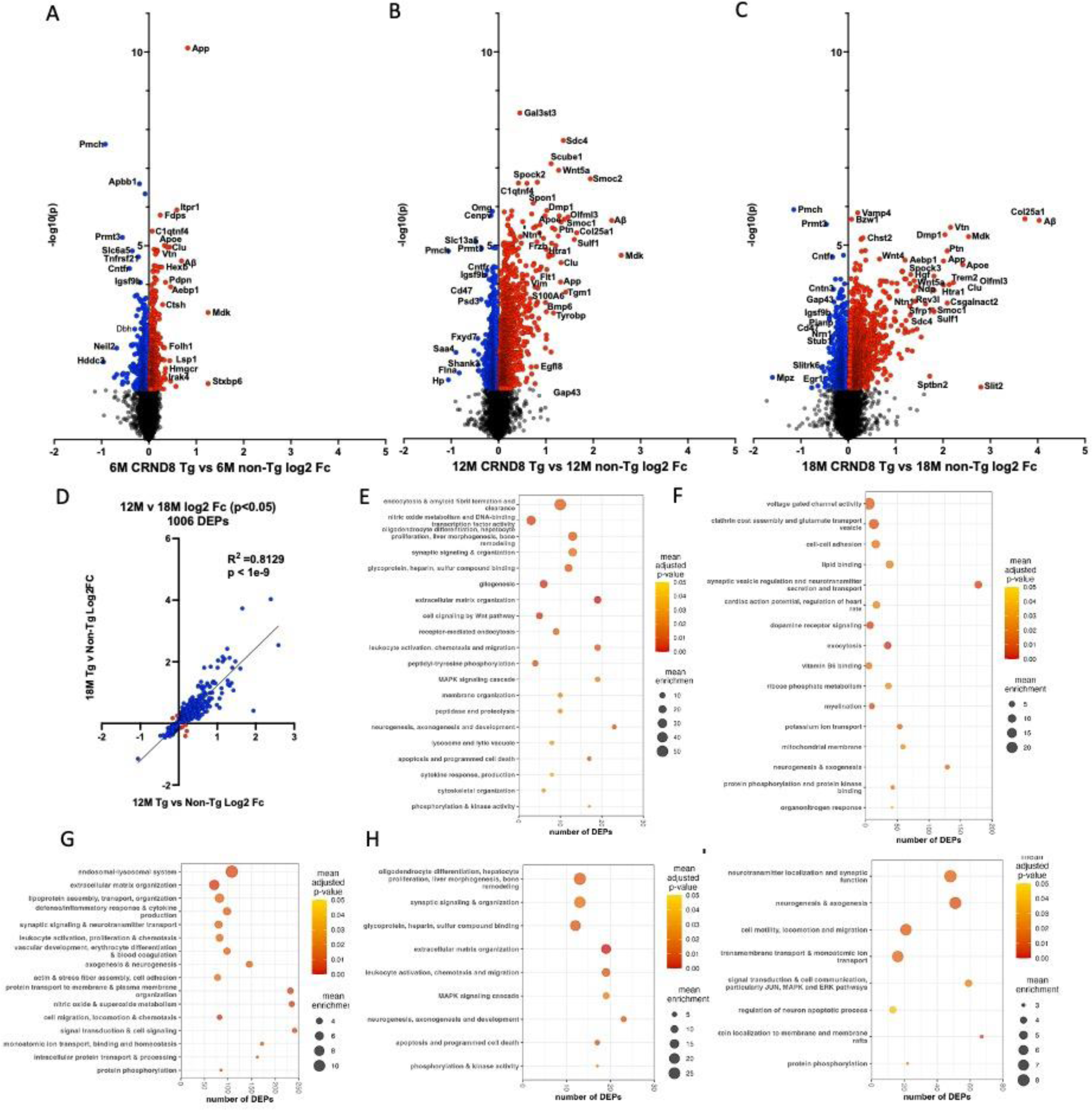
TMT-MS analysis and gene ontologies of CRND8 mice versus non-Tg brains. (A) Volcano plot of the 6M CRND8 cohort. (B) Volcano plot of the 12M CRND8 cohort. (C) Volcano plot of the 18M CRND8 cohort. (D) Correlation of CRND8 brain DEPs between the 12M (x) and 18M cohorts (y). (E) Thematic grouping of gene ontology categories of DEPs increased in all CRND8 cohorts. (F) Thematic grouping of gene ontology categories of DEPs decreased in all CRND8 cohorts. (G) Thematic grouping of gene ontology categories of DEPs (log 2 Fc > 0.2) and increased by >2 fold in the 18M CRND8 cohort relative to the 6M cohort. (H) Thematic grouping of gene ontology categories of DEPs (log 2 Fc > 1) and increased by >2 fold in the 18M CRND8 cohort relative to the 6M cohort. (I) Thematic grouping of gene ontology categories of DEPs (log 2 Fc < −0.1) and decreased by >2 fold in the 18M CRND8 cohort relative to the 6M cohort. Lists of all genes used for gene ontology analyses are provided in Table S7.

Both full-length APP and specific peptides mapping to the human Aβ peptide sequence also distinguishes Tg from non-Tg mice in all cohorts, consistent with human transgene expression and increasing deposition of Aβ (Figure 1A-C, Table S1). Other proteins previously shown to interact with APP such as (Apbb1 (Fe65), Vac14, Rufy2, Tnfrsf21) are also altered at various ages in the CRND8 proteome and several of these (Apbb1, Vac14, Rufy2) show relatively stable alterations across the age groups suggesting that these may be related to mutant APP overexpression and not linked to Aβ or the response to Aβ accumulation per se (Table S1) ^25-27^. Top gene ontology (GO) terms for the DEPs increased in all cohorts highlight heparin and glycosaminoglycan binding, amyloid fibril formation and regulation, vacuolar and lysosomal transport, and immune phagosome and vesicular and lysosomal transport, integrin binding and extracellular matrix (ECM) organization (Figure 1E). Top ontology terms for the decreased DEPs in all cohorts highlight impacts on lipid binding, synaptic vesicle regulation and neurotransmitter release, mitochondrial membrane, and neurogenesis and axogenesis. (Figure 1E).

In the 12 and 18M cohorts there is an appreciable rightward shift in the volcano plots reflecting many proteins that show an age dependent increase in protein level indicating such changes may reflect the evolving “responsome” to enhanced accumulation of Aβ. A subset of the proteins that show the largest fold change increase have been observed to be altered in other APP mouse models and in humans ^9,10,14,18,28,29^ (Table S1). However, in this study using the older CRND8 mice which have extensive parenchymal amyloid deposition in the forebrain and hippocampus and cerebrovascular amyloid deposition (CAA) in the cerebellum, we appear to detect a more comprehensive picture of the alterations in the proteome in response to Aβ deposition in the brain. Top gene ontology (GO) and KEGG terms for the DEPs that are significantly increased in the 18M CRND8 cohort by log2 Fc > 0.2 and increase by more than 2-fold relative to the change observed in the 6M cohort highlight endosomal and lysosomal processes, axonogenesis and neurogenesis, ECM organization, nitric oxide and superoxide metabolism, and glycosaminoglycan degradation, and many other processes (Figure 1G). Limiting the input for these analyses to the subset of proteins with significant log 2Fc > 1 in the 18M cohort, highlight oligodendrocyte differentiation, synaptic signaling and organization, glycoprotein and, heparin, sulfur compound binding, ECM, and neurogenesis and axogenesis (Figure 1H). Top gene ontology (GO) and KEGG terms for the DEPs that are significantly decreased in the 18M CRND8 cohort by log2 Fc < −0.1 and decrease by more than 2- fold relative to the change observed in the 6M cohort highlight terms around neurotransmitter localization and synaptic function, neurogenesis and axogenesis, signal transduction and protein localization to membrane rafts (Figure 1I).

### Integration of the AD human brain and CRND8 mouse brain proteome reveals both conserved and divergent changes

To better understand the similarities and differences in protein levels between human AD and control brains and in theCRND8 Tg and non-Tg mouse brain, we leveraged an extensive proteomic network analysis and the resulting modular framework from human AD and control brains that has been previously published. Using identical TMT-MS methodologies, we have built a deep consensus proteomic network of >8,600 proteins from the dorsal lateral prefrontal cortex of >500 subjects from the Religious Orders Study/Memory and Aging Project (ROSMAP) and Banner Sun Health cohorts ^9^. This network was constructed using the weighted gene co-expression network algorithm (WGCNA) and consists of 44 modules that reflect multiple biological and cell type processes, many of which are altered in AD. Globally, when the levels of 7588 overlapping proteins are compared between the 18M CRND8 Tg and non-Tg brains and AD and control brains there is a significant correlation (p<1e^-9^, R^2^=0.158) (Figure 2A); the strength of the correlation is increased when filtering on the significant DEPs (n =709) in both the CRND8 and human AD brain proteomes relative to respective controls (p<1e^-9^, R^2^=0.458) (Figure 2B). Comparison of the 6M CRND8 proteome changes with AD reveals a much weaker correlations for all overlapping proteins (n=6794, p<1e^-9^, R^2^=0.018) (Figure 2C) but still a strong correlation for shared DEPs (p<1e^-9^, R^2^=0.438) albeit with many fewer shared DEPs (n=161) (Figure 2D). Notably, the correlation for proteins is stronger than for the correlations between protein coding mRNAs in 20M old CRND8 mice vs. non-Tg mice brains and human AD and control brain superior temporal cortex from Mayo Clinic AD vs. control RNAseq data (orthologous protein coding RNAs, n=13066, R^2^=0.079; all shared DEGs, n=1399, R^2^=0.276, Figure S1) ^12,30^.

**Figure 2.**
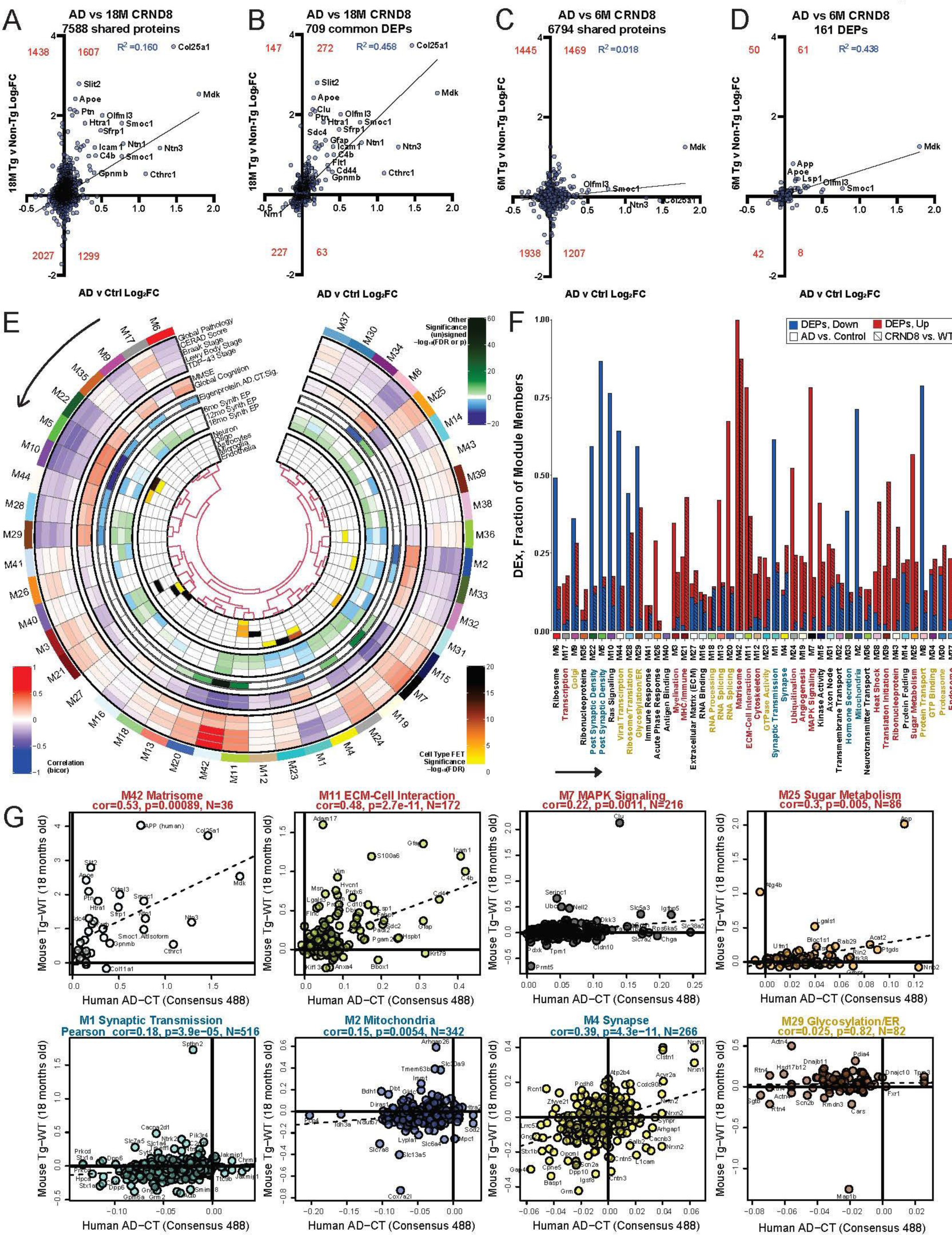
Integrated mouse and human brain proteome comparison reveals concordant and discordant changes between CRND8 mice brains and human AD. (A) 7588 mouse orthologs of human brain proteins quantified in both proteomes of the consensus human AD brain proteome (AD vs. Ctrl) and the 18M mice (CRND8 Tg vs non-Tg) are compared by their effect size (log 2 Fc) in human (x) vs. mouse (y). Red numbers indicate number of proteins in each quadrant. (B) Proteins in panel A were filtered to retain only orthologs nominally significantly changed in both human and the 18M mice. (C) 6794 mouse orthologs of human brain proteins quantified in both proteomes of the consensus human AD brain proteome (AD vs. Control) and the 6M mice (CRND8 Tg vs non-Tg) are compared by their effect size in human (x) vs. mouse (y). (D) Proteins in panel C were filtered to retain only orthologs nominally significantly changed in both human and the 6M mice. (E) Overview of human consensus brain proteome mapping to the CRND8 mouse brain shows trait correlations in human (outer two tracks; red, positive correlation to white, no correlation, to blue, negative correlation), effect size directionality and significance for AD (third track from the outside), the effect size directionalities and significance for CRND8 Tg vs. non-Tg at 6M, 12M, and 18M in cognate mouse synthetic modules (Synth eigenproteins, EP), and hypergeometric overlap significance with cell type marker lists, indicating cell type enrichment of modules in the innermost track. Modules are numbered M1 to M44, ordered by bicor correlation relatedness (dendrogram), from M6, top left, to M37, top right, in a counterclockwise direction. (F) The 44 consensus modules were assessed for fraction of module member proteins achieving differentially expressed protein (DEP) status, discerning decreased DEPs in disease or Tg as blue, and increased DEPs as red in a stacked bar chart. Human and mouse bars for each module are shown side-by-side, left and right, respectively, above each module description. Descriptions below the numbered identifiers match the order of modules presented in the previous panel (*arrow*). The color of description text indicates concordant increases (red), decreases (blue), discordant in direction of change (gold), or non-concordant (black). (G) Mouse orthologs CRND8 18M Tg vs. non-Tg effect sizes are compared to the AD vs. Control effect sizes, module by module, with selected modules shown here. Pearson correlation rho and Student’s significance of correlation are provided along with the number of orthologues mapping to each consensus module plotted. Text color of each module plot title matches the scheme described for the previous panel.

Inclusive correlations of all proteins shared within each of the human modules revealed select modules with high overall correlations in terms of directionality and magnitude of changes, and modules with negative correlations (Figure 2E). More granularly, when we assess the CRND8 mouse proteome changes by evaluating the DEPs and preservation of direction of change within the human AD brain module structure (Figure 2F), we find that there are number of modules in which there is a high degree of preservation between the CRND8 mouse to human proteome in terms of DEP representation. There are also modules for which many DEPs are discordant in terms of directionality, as well as modules where the preservation of directionality is mixed (referred to as non-concordant). Modules showing strong preservation and shared DEPs with increased levels are M42 (Matrisome), M11 (ECM), M7 (MAPK Signaling), M24 (Ubiquitination), M25 (Sugar Metabolism), M19 (Angiogenesis), M12 Cytoskeleton, M39 (Translation Initiation), M21 (MHC/Immune), M38 (Heat Shock), M43 (Ribonucleoprotein), M12 (Cytoskeleton), M3 (Myelination) and M17 (Translation). Modules showing strong preservation and shared DEPs with decreased levels are M22/M5 (Postsynaptic Density), M1 (Synaptic Transmission), M4 (Synapse), M2 (Mitochondria) and M33 (Hormone Secretion). Discordant changes are observed in M9 (Golgi), M44 (Viral Transcription), M29 (Glycosylation) M28 (Ribosome/Translation), M23 (GTPase Activity), M34 (GTP Binding), M8 (Protein Transport), M30 (Proteasome), M16 (RNA Processing), and M20/M13 (RNA Splicing). Examples of the cross-species correlations observed between all proteins within individual modules detected between the human AD proteome and the 18M CRND8 proteome are shown in Figure 2G and Figure S2. These correlations range from robust and significant (M42, M11, M4) to modest but significant (M7, M25, M2) to absent (M29).

### Human AD protein modules show increased magnitude of change in aged CRND8 brains

The human proteome and modular network included brains of individuals who had no reported cognitive decline yet neuropathologic criteria of AD. These were referred to as asymptomatic AD (Asym AD). Here we assessed the synthetic eigenprotein levels in the three CRND8 cohorts to evaluate how the overlapping mouse proteins within these modules change over age and how they compare to the modular eigenprotein changes in human Asym AD, and AD (Figure 3 and Figure S3). These data show that across many modules there is an increase in the magnitude of the eigenprotein change with preserved directionality as CRND8 mice age (e.g., M42, M11, M21, M7, M1, M9). Other modules show different patterns of change in the cohorts. For example, M15 (Kinase activity) and M2 (Mitochondria) show increased eigenproteins in the 6M cohort and then decreased eigenprotein in the older cohorts, whereas M3 (Myelination) shows a decreased eigenprotein at 6M and then increased at later timepoints. Such data demonstrate that the DEP changes observed often show progressive changes over the age range studied and reveal complex relationships between age, pathology, and proteome changes.

**Figure 3.**
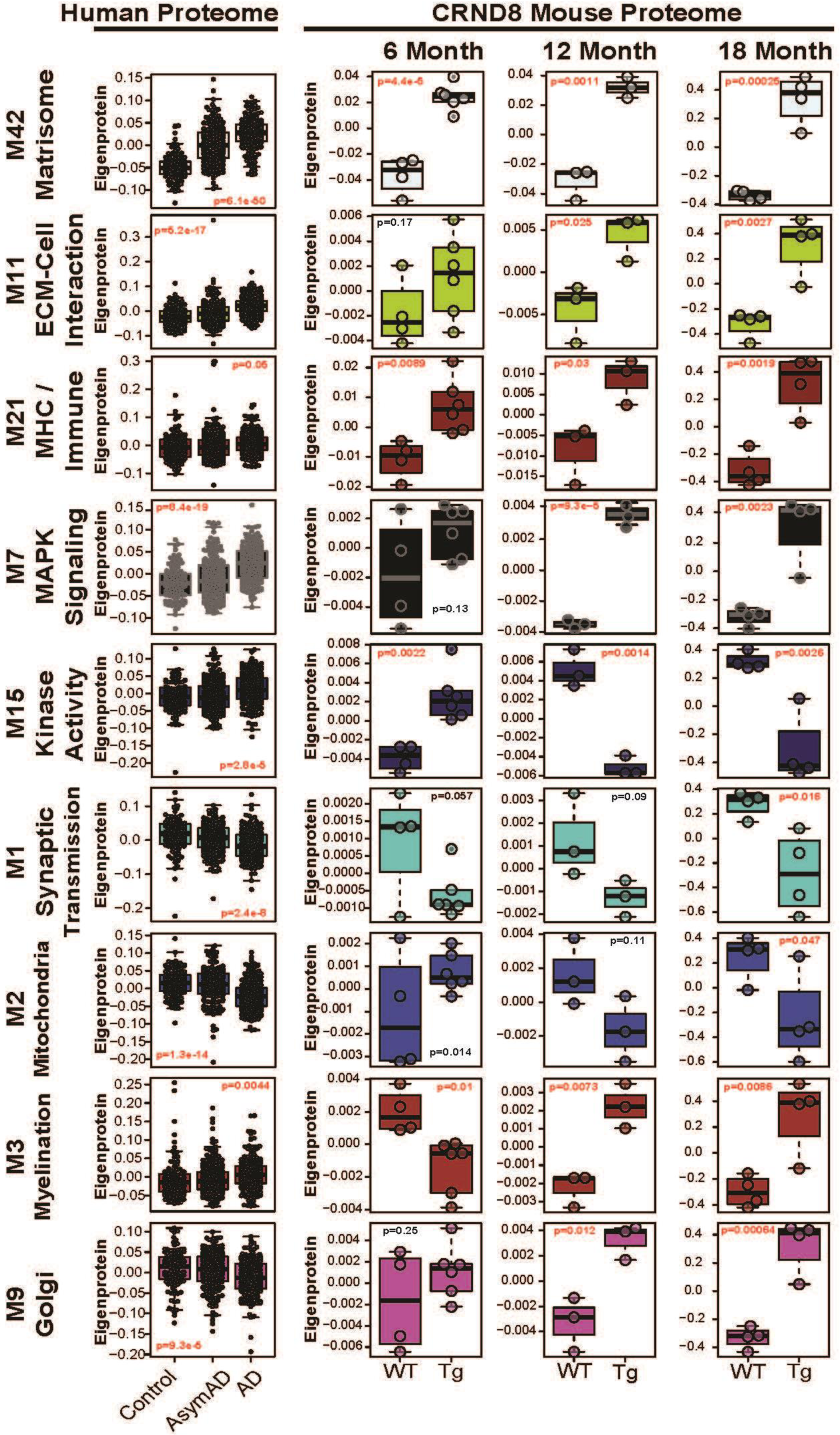
Consensus human AD brain synthetic module eigenproteins in mouse show transient and/or age-progressive CRND8 Tg effects. Nine consensus human modules are shown for the network of healthy control, pathology-bearing asymptomatic AD (AsymAD), and symptomatic AD (AD) individuals totaling N=488 (*left panels*). To the right, the mouse orthologs of hub proteins for these modules were used to calculate the first principal component of variance in the 6M, 12M, and 18M mouse proteomes, allowing for extrapolation of the modules into Tg and non-Tg mice, determination of effect size by module in each mouse cohort, and the significance of Tg effect within each module. Significant one-way ANOVA p values are colored red. Note the Y-axis are different across ages, and in many modules the eigenprotein change between Tg and non-Tg mice increases dramatically with age.

### Correlations between Protein and RNA changes in the 18M CRND8 brains

In human brain, the overall correlation between changes in mRNA and protein levels between AD and control brains is weak ^9^. Here we directly compared the protein and RNA levels in the 18M CRND8 cohort. Analyzing the correlation between the changes in 9319 common gene products detected in the 18M CRND8 vs non-Tg proteome at the protein and mRNA levels (Figure S4A) reveals a weak correlation, which is on par with the overall correlation observed in the human brain (protein vs RNA changes in AD vs Control DLPFC) ^9^. However, when these data are analyzed within the framework of the protein modular networks, a more complex relationship between changes in mRNA levels and protein levels is observed (Figure S4B-D, Figure S5). For example, within M42 there is no correlation between protein and mRNA levels in either human or mice. However, for modules M11 and M21 there is robust and highly significant correlation between protein and mRNA levels in the CRND8 mouse brain. A more modest, but still highly significant, correlation is observed in the same modules in human AD brain.

### Validation of the proteome changes-focus on M42

We have taken a multifaceted approach to validate the protein changes observed with a major focus on M42 proteins, the module with the most robust changes in CRND8 and strongest correlations between mice and human brains. We also included studies of additional DEPs identified only in the CRND8 proteomes that based on biological inference relating to the protein being secreted, a component of the ECM, and known or inferred (based on homology) binding to heparan sulfate (HS) or heparan sulfate proteoglycans (HSPGs) likely had biological roles relevant to M42. We thus refer to this group of proteins as M42+. Several additional DEPs, which showed a log2 Fc >1 in the 18M cohort (Ctss, S100a6, Rev3l), were also targeted in this validation even though they had no obvious links to M42 biology. There was a manifold rationale for the primary focus on M42+ proteins. First, M42 was strongly associated with both core pathological features of AD and cognitive decline and both APOE and APP/Aβ were present in the network. Second, M42, as shown above, was the most conserved module between human AD and CRND8 mice. Third, with a few notable exceptions, protein changes appeared to be dissociated from changes in RNA, implicating possible disruptions in proteostasis. Fourth, many M42 proteins show the largest log2 Fc between AD and control brain and in many cases even more striking log2 Fc changes in CRND8 mice. Finally, M42 proteins, represent a mix of DEPs, several with links to AD and amyloid pathology, but others with little or no previous links to AD and amyloid.

### MDK and PTN co-accumulate with Aβ in plaques and CAA

We focused initial validation studies at the brain tissue level, as successful detection of the DEP *in situ* would enable us to put the protein changes in pathological context. As exemplars of these studies, we highlight work on midkine (Mdk) and pleiotrophin (Ptn). Mdk and Ptn are homologous small secreted signaling molecules known to interact with HS/HSPGs and signal through a set of overlapping receptors (Sdc1-4, Ptprz1, Ptprb, Lrp1, Ncl, and Integrins α4β1, α6β1, α5β3) ^31,32^. In the TMT-MS data, Mdk is significantly increased in all three cohorts studied (log2 Fc) and has the largest log2 Fc of any DEP in the 6M cohort, whereas Ptn shows significant increases in the 12 and 18M cohorts. Neither mRNA is altered in bulk transcriptomic studies of human AD brain or APP CRND8 mice brains, though the mRNA levels (Fpkm) are significantly higher for Ptn than Mdk in both human and mouse brain. Select Mdk/Ptn receptors are also altered at the protein level in human AD, CRND8 or both. For example, Sdc2 and Sdc4 are significantly increased in AD and CRND8 brains; Ptprz1 is significantly downregulated in CRND8 brains; Lrp1 is significantly upregulated in AD and CRND8 brains; one isoform of Ncl is decreased in 18M CRND8 mice; Itga4 and Itgb4 are increased in both CRND8 and AD; Itga5 and b3 are increased in CRND8 mice (Table S1).

Immunostaining for Mdk or Ptn shows both exhibit a high degree of overlap with Aβ in CRND8 mice (Figure 4A) and in human brain (Figure 4B). Not only do Mdk and Ptn co-localize to plaques (Figure 4A), but they also co-localize to Aβ amyloid in blood vessels associated with CAA, which is most easily observed in CRND8 in the meningeal vessels of the cerebellum (Figure 4A). In humans, MDK and PTN are detected in both plaques and CAA deposits (Figure 4B). Though both Ptn and Mdk are detected in plaques, they show distinct staining within the cored plaques (Figure 4B-D). Mdk is more prominent in the center of the plaque core in mice and intensity of staining decreases towards the plaque periphery (Figure 4C). Ptn staining is almost the mirror image of Mdk, and it is localized more on the periphery of the fibrillar plaque. In many plaques Ptn does not stain the center where Mdk is most strongly localized (Figure 4C, D). In contrast, in humans, though MDK is more intensely detected in the plaque center, the entire plaque is typically stained, and PTN diffusely stains the fibrillar plaque (Figure 4B). In CAA in the CRND8 mice, both Ptn and Mdk non-homogenously label amyloid deposits detected by Thioflavin S (Thio S) (Figure 4D). In vessels there are some areas where Ptn staining does not overlap with Thio S staining, this might be attributable to expression in pericytes, as Ptn is highly expressed in these cells. Further, staining of Mdk and Ptn in CRND8 mice brains shows co-localization with plaques from 3-12M, as well as staining of plaques in numerous lines of APP transgenic mice (Figure S6A).

**Figure 4.**
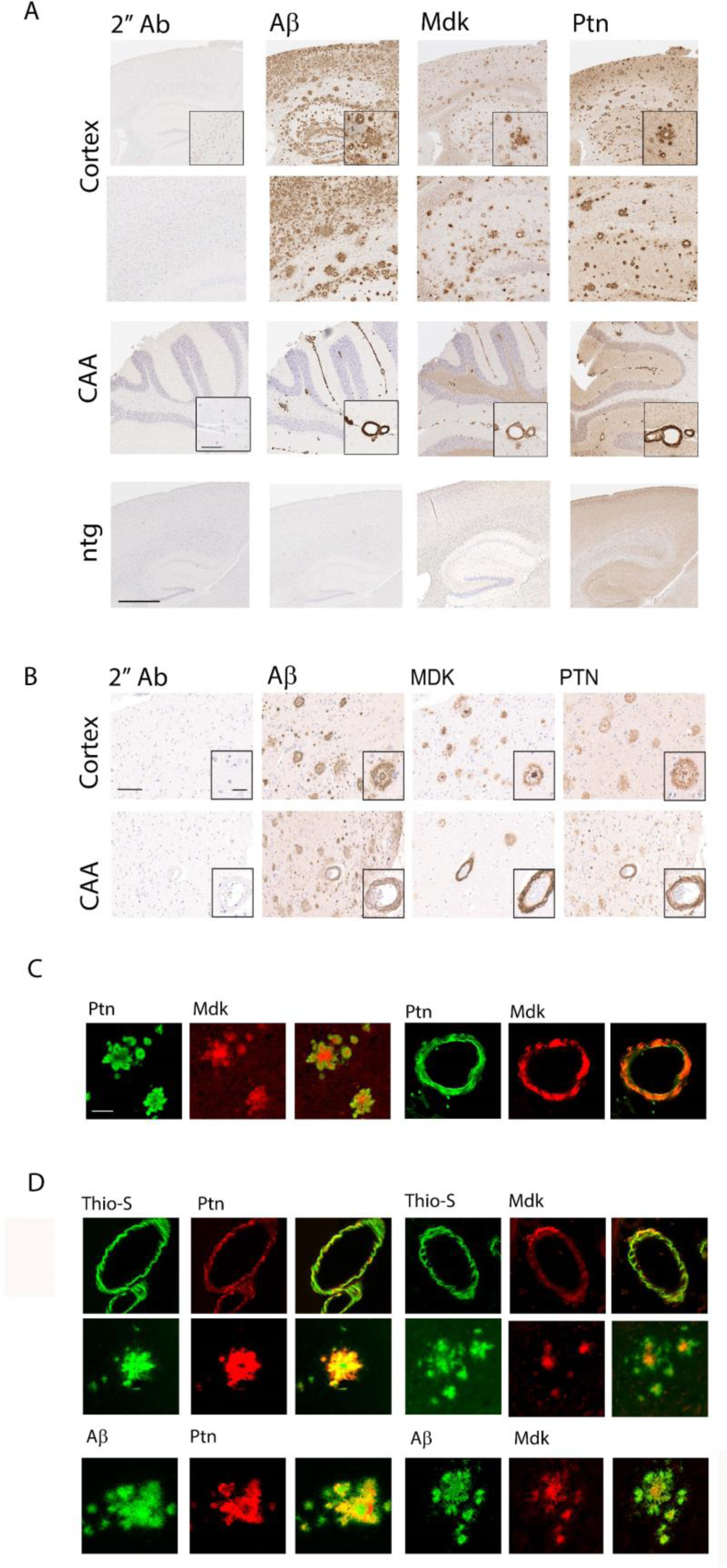
Mdk/MDK and Ptn/PTN colocalize with Aβ in amyloid plaques and CAA in CRND8 mice and human AD. (A) Paraffin slides containing brain tissue from 15-18M CRND8 mice were stained with mouse anti-pan-Aβ, sheep anti-Mdk or rabbit anti-Ptn antibodies. Low magnification (scale 500 μm) images and high magnification (inset, scale 50 μm) were taken from cortex (plaques) and cerebellum (CAA). (B) Representative low magnification (scale 100 µm) and high magnification (inset, scale 30 µm) of postmortem paraffin embedded tissue sections of human frontal cortex from patients with high AD neuropathological changes stained for MDK and PTN. (C) Plaques on paraffin slides containing brain tissue from old CRND8 mice were stained with anti-Mdk and anti-Ptn antibodies and visualized with fluorescent secondary antibodies (anti-rabbit green and anti-sheep red), scale 50 μm. (D) Plaques and CAA on paraffin slides containing brain tissue from old CRND8 mice were stained with anti-pan-Aβ, anti-Mdk or anti-Ptn antibodies and visualized with Thio-S (green) or fluorescent secondary antibodies (anti-mouse red, anti-rabbit red, anti-sheep red), scale 50 μm.

Western blot analysis of 8M Urea extracts from 20M CRND8 mice brains shows an almost qualitative accumulation of both Mdk and Ptn in both male and female CRND8 brains, relative to the non-transgenic lines, suggesting that TMT-MS methodology may underestimate the true fold change in protein levels (Figure S6B) ^33^. As a characteristic feature of Aβ deposited as amyloid is lack of solubility by strong detergent extraction, we examined the crude solubility profiles of Mdk and Ptn in CRND8-Tg brains sequentially extracted with TBS, RIPA, 2% SDS and 8M Urea (Figure S6C). Though relatively low levels of Mdk and Ptn are detected in the TBS and RIPA extracts, the largest amounts were present in the 2% SDS fraction and none was detected in the 2% SDS insoluble urea soluble fraction. Notably, the relative proportion of Mdk or Ptn appearing in each extraction was similar in both the CRND8 Tg and non-Tg brains. This lack of change in gross solubility as the proteins accumulate suggests that they are not undergoing an amyloid-like structural change, which is typically associated with detergent insolubility ^34^.

### Validation of additional M42+ DEP changes

We initially screened many commercially available antibodies to proteins in M42, few of these antibodies worked in paraffin embedded tissues (Tables S11, S12); therefore, we generated many new rabbit polyclonal antibodies. Data showing these antisera can detect the target protein, is provided in Figure S8A. In mice, we evaluated antibody staining of Tg and non-Tg brains in formalin, paraformaldehyde, and ethanol fixed sections. As shown in Figure 5A, we find that many of these antibodies stain amyloid plaques in CRND8 mice (non-Tg controls are shown in Figure S7A), suggesting that many M42+ proteins accumulate, at least in part, due to sequestration within plaques. In mice, a curious feature of these studies is that the localization of the proteins with the plaques is often morphologically distinct. Slit2, Vtn, Olfml3, Ndp, Rev3l, Ntn1, Hhipl1, Bmp6, Ctss, Frzb, Sema3e, Smoc2 and Nog are preferentially localized within the plaque center as observed for Mdk. Col25a1 and Dmp1 appear in the more peripheral regions of the fibrillar plaque akin to Ptn. Another subset shows more diffuse homogenous staining of the plaques (e.g., Vtn, Sdc4, S100a6). Still others show a patchy, nonhomogeneous, staining of the plaques (e.g., Hgf, Spock1, ApoD, Egfl8, C1qTnf4, Slit3), which may in part be attributable to staining of dystrophic structures around or within the plaque. In addition, we find that fixation conditions can dramatically impact the staining pattern observed and that the effects of fixation are antibody dependent (Figure S7B). Dual labeling studies with Aβ or Thio S and a subset of these proteins reinforce the different location within the plaque of select M42 proteins (Figure S7C). Finally, except for Mdk, Col25a1 and Ptn, which appear to stain most non-diffuse plaques, we note that only a subset of the plaques detected by Aβ immunohistochemistry is stained (Figure 5A). As the anti-Aβ monoclonal used for staining has subnanomolar affinity, additional studies will be needed to further evaluate the extent of co-localization using additional immunologic reagents.

**Figure 5.**
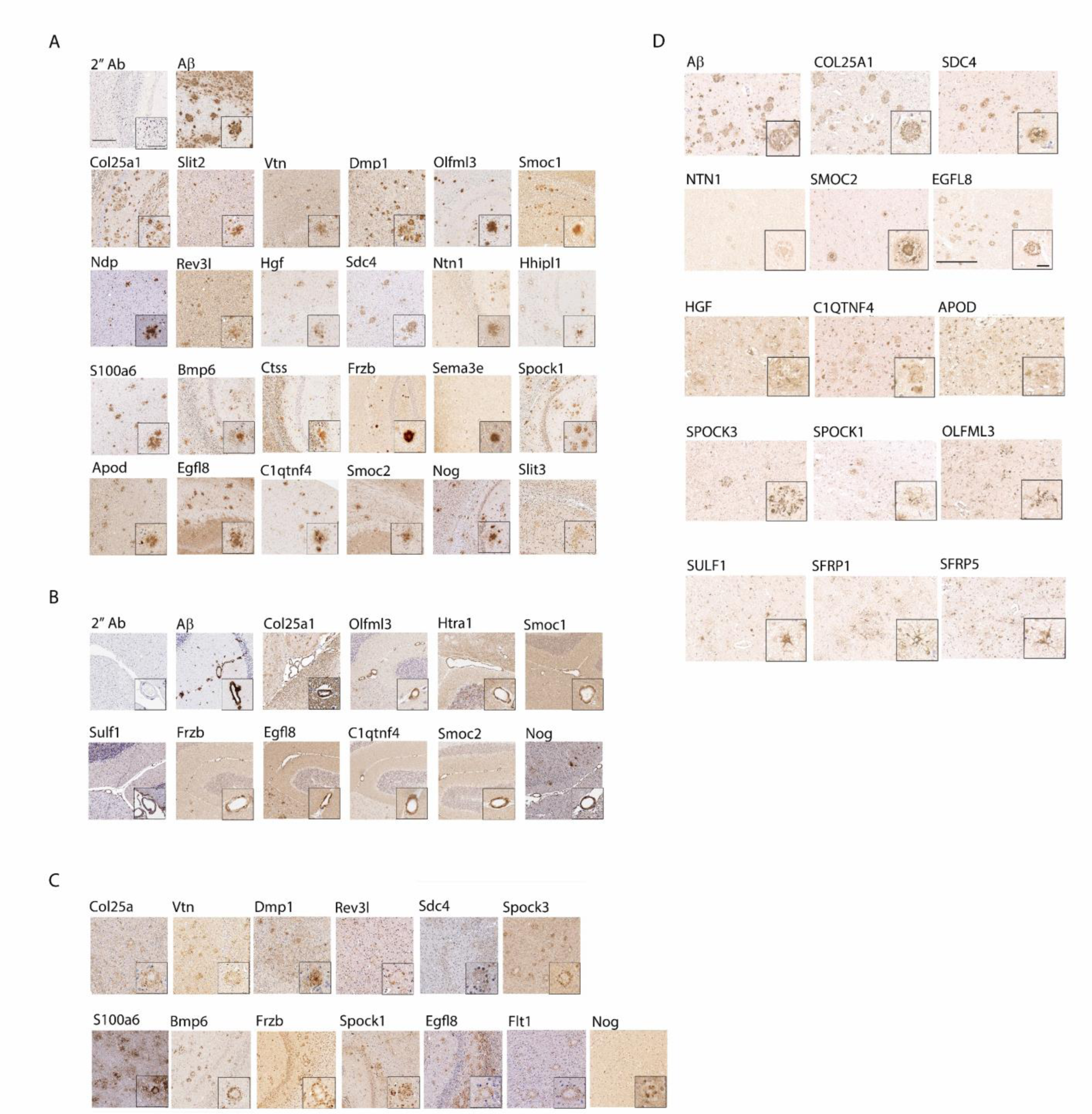
M42+ DEPs colocalize with amyloid pathology. Brain tissue from 15-18M CRND8 mice were stained with the anti-sera against the indicated proteins. Anti-pan-Aβ antibody served as a reference for amyloid pathology, secondary antibody alone – as a negative control. All proteins are listed in order of log 2 Fc (highest to lowest). Low magnification (scale 200 μm) images and high magnification (inset, scale 50 μm). See Figure S7 for non-Tg staining. (A) DEPs in amyloid plaques in the CRND8 cortex. (B) DEPs detected in CAA in the cerebellum. (C) DEPs detected in dystrophic processes surrounding amyloid plaques in the cortex. (D) Representative low magnification (scale 100 µm) and high magnification (inset, scale 30 µm) of postmortem paraffin embedded tissue sections of human frontal cortex from patients with high AD neuropathological changes stained for Aβ, EGFL8, C1QTNF4, COL25A1, HGF, APOD, SDC4, NTN1, SMOC2, OLFML3, SPOCK3, SPOCK1, SULF1, SFRP5 and SFRP1. There is selective staining of plaques by antisera for COL25A1, SDC4, NTN1, SMOC2, and EGFL8. Weaker staining of plaques and cellular staining is observed for HGF, C1QTNF4 and APOD. Dystrophic processes are stained with anti-serum to SPOCK3, SPOCK1 OLFML3. Glial cells are detected with anti-sera to SULF1, SRRP5 and SFRP1.

CAA Aβ deposition is easily assessed in the cerebellum of 12M CRND8 mice. In Figure 5B we show the subset of antibodies that stain CAA. In addition to Mdk and Ptn a smaller group of the M42+ DEPs are detectable in CAA Aβ deposits (Col25a1, Olfml3, Htra1, Smoc1, Sulf1, Frzb, Egfl8, C1qtnf4, Smoc2, Nog). All of these can also be shown to co-localize with Thio S staining in the vessels (Figure S7C).

Several M42 CRND8 DEPs show staining adjacent to the fibrillar plaque (Figure 5C). Spock3, Bmp6, Frzb, and Nog, show staining patterns highly consistent with the classic dystrophic neurites (DN) often visualized with anti-ubiquitin or other DN markers. In contrast, Col25a1, Vtn, Rev3l, Sdc4, S100a6, Egfl8, Spock1, and Flt1 show peri-plaque staining that is distinct from the classic DN pattern. Selective staining of reactive astrocytes is noted for Htra1, Sulf1, Sfrp1, HhIpl1, Spock3, Sema3d, and Smoc2 (Figure S7D).

We have been able to localize a smaller subset of the M42 proteins in AD postmortem brain tissue (Figure 5D). EGFL8, C1QTNF4, COL25A1, HGF, SDC4, NTN1 and SMOC2, all show staining in plaques. We would note that EGFL8 was not detected in the TMT-MS studies in human brain, but Egfl8 was detected as a DEP in CRND8 mice. OLFM3, SPOCK3, SPOCK1, show staining of dystrophic structures around plaques. SULF1, SFRP5 and SFRP1 stain reactive astrocytes.

In supplementary Table S9, we compile the staining observed and annotate our findings versus previous studies in the field, we also classify the cell type expression observed for the proteins that we detect in these immunohistochemistry studies using data available at the Allen brain atlas ^35^. The extended M42+ members appear to arise from multiple cell types representing all major brain cell types. This observation is consistent with the biological annotations of M42 (matrisome, ECM) and likely reflects extracellular processes regulating or responding to abnormal extracellular protein accumulation. Many of the mRNAs encoding these proteins are expressed at low levels (< 5 Fpkm). Further within the M42 DEPs only Olfml3, which is selectively expressed by microglia, shows significant increase in mRNA levels in the 18M RNAseq data. In previous RNAseq studies of CRND8 mice brains, we have found that Olfml3 mRNA levels are increased at 6, 12 and 20M ^30^. Ctss and S100a6 also show significant increases in mRNA levels, but as noted previously, are not part of the core M42 module; the associated increase in these mRNAs may be due to the reactive gliosis in the CRND8 brain. In addition to these immunohistochemical studies and data on Mdk and Ptn, we can also demonstrate increases in the brains of older CRND8 mice of Vtn, Htra1, Smoc1, Frzb, ApoD and Nog western blot analysis of the CRND8 mice (Figure S8B).

### Mdk and PTN overexpression increase parenchymal amyloid and CAA in CRND8 mice and accelerate amyloid aggregation in vivo

Given the robust data showing the co-localization of Mdk and Ptn with both plaques and CAA Aβ deposits and the early increase observed in the 6M TMT-MS data for Mdk, we evaluated the impact of rAAV-mediated overexpression of Mdk or PTN on amyloid deposition in 2 cohorts of CRND8 mice. The first study examined the impact of P0 ICV injection of rAAV2/8 vectors encoding Mdk or PTN after 3 months of age, the second at 6M of age. In both cohorts, similar impacts are observed, for brevity we will primarily describe the 6M analysis here (Figure 6A-C). The 3-month overexpression data is presented in Figure S9A-C. Widespread over expression of Mdk and PTN is detectable in the brain in both cohorts. Even in the small early compact plaques in the 3M cohort it is possible to detect Mdk and PTN staining within the plaques (Figure S9A). Mdk and PTN both increased parenchymal plaques (Figure 6A) and CAA (Figure 6B) as assessed by immunohistochemistry. In previous studies of CRND8 mice we have not noted sex differences in amyloid deposition or other phenotypes, though these studies were not designed to be powered to detect sex differences, post hoc analyses do not reveal any major impact of sex. Both also increased Aβ levels in SDS and Formic Acid (FA) fractions, but not in the RIPA fractions (Figure 6C). These studies establish that overexpression of either Mdk and Ptn can accelerate amyloid deposition into fibrillar plaques and CAA in two cohorts. In the 6M cohort, we also evaluated reactive gliosis (Figure S9D, E). Both Mdk and PTN increased astrocytosis assessed by Gfap staining, and PTN increased microgliosis as assessed by Iba1 staining. Mdk overexpression also increased Iba-1 staining but this elevation was not significant. We also assessed the impact of recombinant PTN and MDK on amyloid aggregation *in vitro* using standard ThT fluorescence assays (Figure 6D). Both reproducibly and significantly accelerated Aβ42 aggregation. Mdk also increased Aβ40 aggregation, whereas only a small impact for PTN on Aβ40 aggregation was observed. End of assay total ThT fluorescence was also increased in all cases except for the Aβ40 PTN study, suggesting increased β-sheet formation.

**Figure 6.**
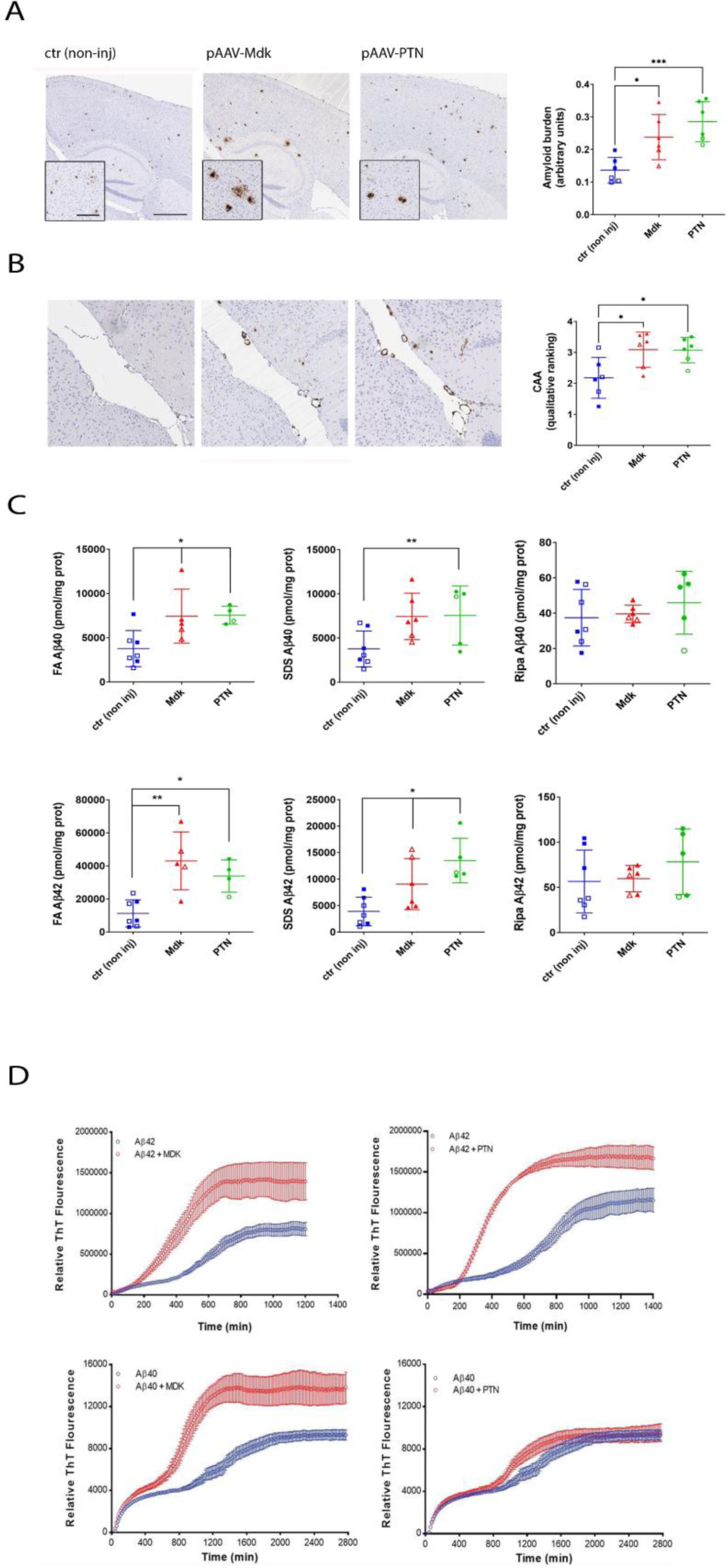
Mdk and PTN overexpression increase parenchymal Aβ amyloid deposition and CAA in 6M CRND8 mice. Mice were intracerebrally injected with rAAV2/8-Mdk, rAAV2/8-PTN or not injected (control) at P0 and aged 6 months. (A, B) Representative images of cortex and hippocampus 9A) or CAA (B) stained with biotinylated anti-Aβ mAb 33.1.1 (anti-Aβ 1-16). Scale bar: 500 µm, 50 µm (inset). Quantification data of the entire brain plaque count in three non-consecutive sections represented by a scatter dot plot of male (closed circle/square) and female (open circle/square) ± standard error of the mean. n=5-15. Statistical analyses by one-way ANOVA test (*, p<0.05; ***, p<0.001). (C) RIPA, 2% SDS and 70% formic acid (FA) extracted Aβ42 and Aβ40levels were detected by ELISA and plotted as scatter dot plot of male (closed circle/square) and female (open circle/square) ± standard error of the mean. n=5-12. Aβ42 and Aβ40 levels were quantified with corresponding one-way ANOVA and paired comparison test (*, p<0.05; **, p<0.01, ***, p<0.001). (D) MDK and PTN accelerate amyloid aggregation in vitro as detected by real time Thioflavin T (ThT) assay.

### Select M42+ proteins bind to Aβ fibrils and an amyloid derived from a non-human peptide, non-homologous peptide

Many M42 proteins are known or can be inferred by study of other family members to be HS/HSPG binding proteins. We have performed limited studies on heparin interaction with Mdk and Ptn and can confirm strong interactions with immobilized heparin (Figure S10). We evaluated a larger number of these for the potential to interact with Aβ42 fibrils and a non-homologous amyloid formed from a 12 amino acid adenovirus shaft peptide (AVS fibrils) ^36,37^. For these studies we turned to a simple amyloid-pulldown assay to screen for interactions with insoluble amyloids ^38-40^. As M42 proteins contain numerous intrachain disulfide bonds and other post translational modifications (PTMs), we expressed a subset of these proteins in HEK293 cells and collected conditioned media containing the secreted proteins. Aβ42 fibrils or the AVS fibrils were then incubated and spun down to evaluate potential interactions with amyloid structures. For the subset of proteins which we were able to detect in the media, we can show that APOE4, Mdk, Vtn, Ptn, Olfml3, Htra1, Smoc1, Sulf1, Frzb, ApoD, C1qTnf4, Smoc2, and Nog bind both Aβ42 and AVS amyloids (Figure 7A). Sfrp1 is a notable exception as we only observed it binding to Aβ fibrils in this assay. Such studies provide initial data that many of these proteins bind a generic amyloid structure.

**Figure 7.**
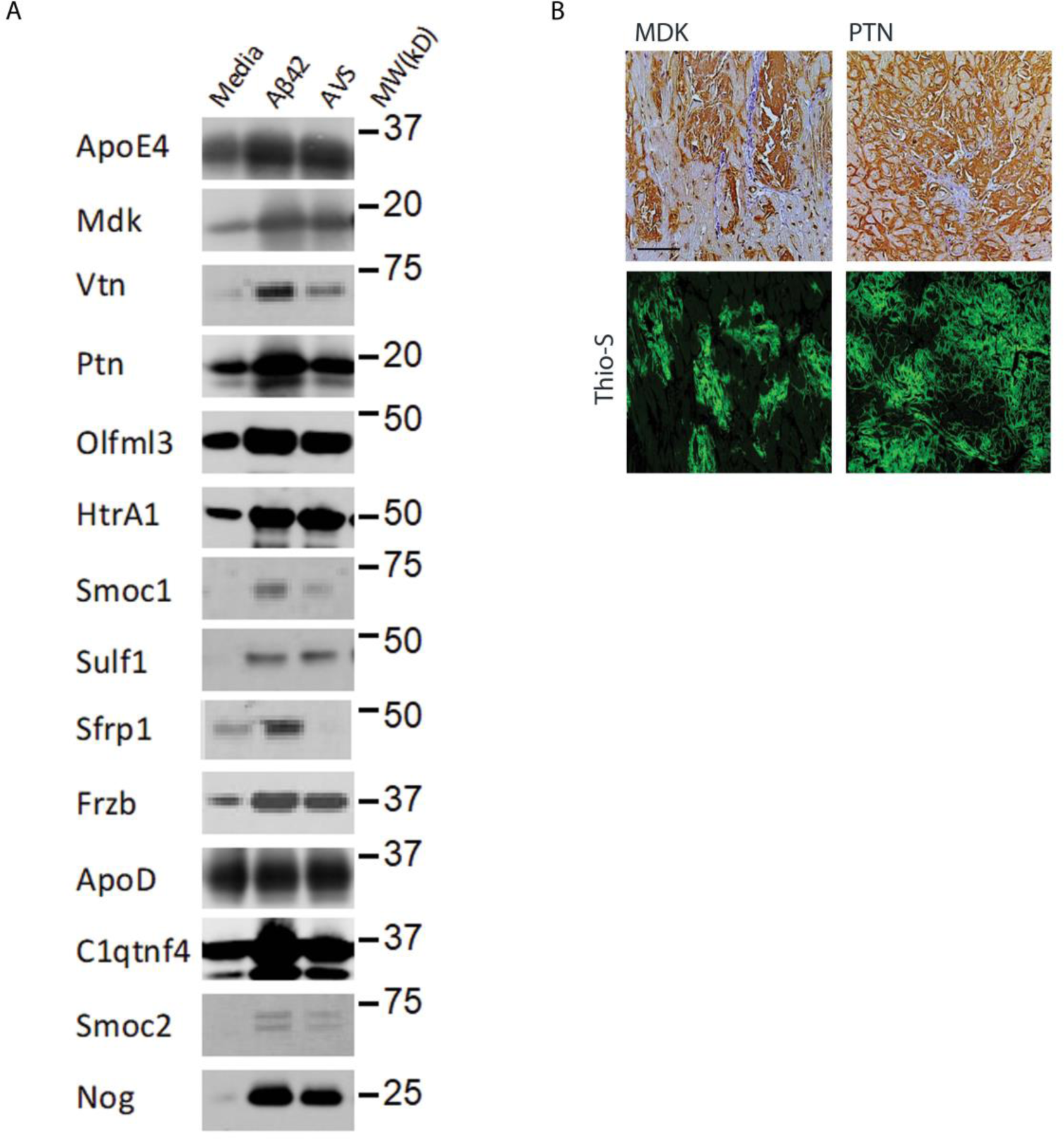
Amyloid binding and co-localization in TTR amyloids. (A) M42+ proteins secreted into the media of transiently transfected HEK cells bind to Aβ42 and AVS amyloids. Data are representative of three independent experiments. (B) Representative images of immunohistochemical stains for MDK, PTN or Thio-S on human cardiac amyloidosis sections. Scale bar, 50 µm.

### PTN and MDK are colocalized with transthyretin (TTR) amyloid in the human heart

Though the amyloid pulldown assay shows the ability of multiple M42 proteins to interact with a generic amyloid structure, we wanted to orthogonally test this hypothesis by evaluating whether some of these proteins may accumulate in other human amyloid diseases. For these studies we have focused on in *situ* detection of MDK and PTN, as the antibodies for these robustly detect these proteins in formalin fixed human tissues. As shown in Figure 7B we find that both MDK and PTN co-localize with TTR amyloid in the heart (the third most prominent human amyloid disease) ^41^. A series of additional TTR cardiac amyloid cases show consistent co-localization of PTN and CLU, and more variable co-localization of MDK, with the TTR amyloid fibrils (Figure S11).

### DEPs in CRND8 mice in the context of AD genetic studies

Previous transcriptomic studies of APP mice have suggested that very highly upregulated DEGs are enriched for genes under AD GWAS loci or known genetic risk factors for AD ^42^. To examine whether DEPs in the CRND8 mice might provide similar insights, we integrated data from three previous GWAS studies of AD to evaluate whether DEPs in CRND8 mice reflect some aspects of the genetic architecture of human AD (Table S1, Figure S12A) ^43-45^. Integration of the 18M CRND8 TMT-MS data with these AD genetic studies show that many established AD risk genes and prioritized GWAS loci (e.g., *APOE, BIN1, TREM2, CLU, SORL1, ACE, ADAM17, SORT1, INPP5D)* are DEPS in the 18M CRND8 mice. Abca1, recently implicated as genetic risk factor for AD, is also a DEP in both the 12m and 18M CRND8 cohorts ^46^. Notably multiple M42+ proteins besides APOE show nominal significance in the GWAS studies including *COL25A1, SMOC1, HTRA1, DMP1, MDK, SDC4, SPOCK2, NXPH1*, and *C1QTNF4*, the latter of which lies in the *CELF1* locus and shows the highest p value of M42 members for AD association. Finally, we note that Grn, Sppl2a, Tmem106b and Notch3 are all DEPs in the 18M CRND8 proteome. *GRN* and *TMEM106B* are genetically associated with FTD-GRN, and SPPL2A is known to cleave TMEM106B potentially linking it to the biology of FTD-GRN ^47,48^. Further, *NOTCH3* mutations are associated with CADASIL ^49^. Given debate over genetic pleiotropy or inclusion of non-AD dementia in the large GWAS studies (clinical phenocopy inclusion) ^50,51^, these data suggest there are biological links between proteins implicated in FTD and CADASIL and amyloid pathology in AD. A final intriguing observation from these data is that multiple proteins near the *APOE l*ocus are DEPs in the CRND8 mice. There is conservation of the extended *APOE* locus between mice and humans, and such data may point to complex biology around the *APOE* locus that could contribute to AD risk.

### DEPs in CRND8 mice in the context of AD CSF biomarker studies

We also evaluated the 18M cohort CRND8 DEPs in the context of human AD CSF biomarker studies, for this integration we chose a recent data set in which AD and control CSF were compared using a highly similar proteomic pipeline for both TMT-MS data generation and analysis ^52^. However, we would note that this data shows significant overlaps with other large scale proteomic analyses of AD versus control CSF ^52-62^. Of the 1839 in the human CSF proteomes 257 of these are significantly altered and of these 62 are significantly changed in the 18M CRND8 mice brains. 39 of these changes are congruent in directionality between the mouse brain and human CSF and 23 are discordant in directionality (Figure S12B, Table S13). Notably, 29 of the 18M DEPs are significantly altered in the same direction in human AD brain as in the mouse model, irrespective of direction of change in CSF, and only 5 show a discordant directionality with human AD brain change. Overall, these data support a hypothesis that some significant and reproducible changes in human CSF may reflect changes in the brain related to amyloid.

## DISCUSSION

By integrating newly generated deep TMT proteomic data from CRND8 mice brains with previously generated large-scale proteomic data from human AD brain, we have begun to define the Aβ/Aβ amyloid “responsome” that is partially conserved between this mouse model of Aβ deposition and human AD ^9,10^. Within select protein modules, DEP and overall protein changes are highly concordant; however, the presence of multiple discordant protein network modules and modules that are simply not concordant, reinforce the concept that these transgenic mice are incomplete models of AD ^9,10,30^. Nevertheless, these data and analyses establish a paradigm for contextualizing changes that occur in human AD and provide an unbiased view of the degree to which a model system reflects, or does not reflect, the complex proteomic changes in the human AD brain. Indeed, there is much debate about what defines a “better mouse model”, these data suggest that a comparative omics approach particularly those that analyze the proteome, can provide a less biased manner to address this question ^63,64^.

As discussed in detail below, we have primarily focused validation studies on M42, the most concordant module between human and mice, a module that in humans correlates best with amyloid and tau pathologies and strongly correlates with cognitive status ^9,10^. However, we document other concordant modules and discordant or non-concordant modules as well. This modular framework may offer initial insights into what aspects of AD pathology the mice mimic and what modules they do not, but validation and contextualization of these will be required to better understand how these modules fit into the complex architecture of AD pathology ^2^.

As CRND8 mice do not show overt neuronal loss ^65^, the finding that numerous modules relating to neuronal and synaptic function (M1, M4, M5, M22, M33) show conservation is intriguing. Numerous APP models, including CRND8 mice have been shown to have synaptic alterations ^66-68^, and in many cases these alterations occur at relatively early ages ^63,69,70^. Notably, our data show a progressive impact on synaptic protein modules as assessed by eigenprotein levels. However, for the majority of human and mouse proteins in these modules the magnitude of change as assessed by TMT-MS is quite small. Of interest, neuritin (Nrn1) and small glutamine-rich tetratricopeptide repeat-containing protein β (Sgtb) which are implicated in cognitive resilience to AD pathologies are both consistently downregulated DEPs in the older CRND8 cohorts ^71,72^. Nevertheless, these studies highlight the disconnect between Aβ amyloid deposition models and human AD, as these models do not progress to an overt neurodegenerative state and lack neurofibrillary tangle (NFT) pathology.

Several of the eigenprotein changes in the CRND8 brains assessed in the context of the human AD modular framework show a consistent alteration in directionality and an increased separation between Tg and non-Tg from 6M to 18M (e.g., M42, M28, M11, M12, M4, M23). Other modules, such as M3 (Myelination, down at 6M and up at 12/18M) and M15 (Kinase Activity, up at 6M and down at 12/18M) show discordant directionality across the cohorts. Notably, for myelination there is evidence in other APP mice models that there are temporal changes in oligodendrocyte lineages and myelin that may be consistent with this data ^73-75^ and, more generally, there is growing evidence for oligodendrocyte lineage and myelin alterations in AD ^76-78^ and other tauopathies ^76,79^. However, these specific proteomic changes need to i) be validated in this model and ii) further contextualized by refining these proteomic and validation studies through inclusion of a broader cross-sectional series of ages and additional models with differential temporal sequence of amyloid deposition.

We also may need to refine how we view concordant, non-concordant, and discordant changes between the model and human protein modular networks. A good example of this is M21, the MHC/immune module, though the correlation within this module not significant, in general most proteins in this module are upregulated in both human and mice, however, in humans the protein upregulation of immune module is much less robust, although transcriptome upregulation is significant in both human and mouse brains in the context of AD ^11,80^. This data suggests that mice respond to amyloid deposition with a more robust innate immune response than humans, perhaps because of i) the compressed time frame for amyloid deposition in these models ii) impacts of aging on the human brain which may alter baseline immune status ^81,82^ or iii) inherent interspecies differences in synthesis and clearance of these proteins.

More broadly, future studies will be needed to guide our understanding of the proteomic changes in mouse models that mimic human AD and those that do not. Similar deep proteomic studies on tau mice might inform on protein modules more related to NFT pathology. We also should consider that discordant modules in mice such as upregulation of M9 (Golgi) that includes ER to Golgi trafficking proteins and lysosomal hydrolases and proteases, might serve protective functions that could impact the appearance of downstream pathologies and overt neurodegeneration in the models ^83,84^.

To build a highly significant deep TMT-MS proteome from human AD brain required hundreds of samples ^9,10^, in contrast using small cohorts of CRND8 mice we can generate highly reproducible data. Further, by looking at cohorts of different ages, we can infer changes that might be attributable to transgene overexpression versus increasing level of Aβ deposition. The CRND8 mouse is a robust and aggressive model of amyloid deposition ^65^. CRND8 mice have shown a very reproducible phenotype across many studies, with no easily discernable sex-based impacts on pathological phenotypes ^39,68,85^. Notably our data is highly consistent with previously published proteomic data on 5X FAD mice, but because of the depth, multiple ages, and larger groups sizes we appear to be cataloging a larger number of proteomic changes ^14,29^. From a translational point of view, these data reinforce the notion that preclinical studies probing impacts of therapies targeting amyloid or pathologies downstream of amyloid should not just be tested in younger mice with limited amyloid deposition but should also be evaluated in the context of impacts on older mice with extensive amyloid deposition.

For the reasons described previously, we focused validation efforts on the M42+ module. We had known from previous studies that several of the core M42 members including APP, APOE and COL25A1 accumulate in amyloid plaques ^86-90^. Here we provide additional evidence that many of the core human M42 module are present in plaques in mice (Mdk, Ntn1, Smoc1, Sdc4, Olfml3, Sfrp1, Spock1, Frzb, Htra11, Slit2), we also show that many other proteins emerging from the CRND8 proteome (Vtn, Dmp1, Ndp, Rev3l, Hgf, HHipl1, S100a6, Bmp6, Ctss, Sema3e, ApoD, Egfl8, C1qtnf4, Smoc2, Nog and Slit3) are also present in the plaques in mice. A subset of these M42+ proteins also are found in CAA, others in dystrophic processes around plaques and others in reactive astrocytes. We have successfully localized a smaller number of these proteins to plaques and other pathological features in humans. Given the conserved protein changes in mice and humans seen by proteomic analyses, this discrepancy may be attributable to fixation conditions and epitope exposure, which may differ between mice and humans. Our current data showing how fixation conditions can alter the pathologies detected by select antibodies in mice, highlight the importance of developing multiple high-quality antibodies to extend and further validate these initial pathological assignments.

Notably, two proteins Egfl8 and Hgf were only detected and present as DEPs in the CRND8 proteome, but both HGF and EGFL8 appear to be plaque components in humans. Such findings serve as an exemplar of the power of utilizing a model system to inform on human pathology. Further, these studies broadly serve as a strong affirmation of the power of WGCNA to inform pathobiology of a complex disease ^91^. Indeed, M42 is the most conserved protein module network between CRND8 mice and human AD and shows the strongest correlation with amyloid in humans ^9,10^; these validation studies highlight its direct links relevant to amyloid pathology or response to amyloid pathology.

Some of the more nuanced data emerging from these validation studies relate to the unique patterns of localization of these proteins within fibrillar plaques. This feature has been most rigorously studied for Mdk and Ptn which show unique and reciprocal distributions in the CRND8 plaques, and distinct distributions in human plaques. Overall, there appear to be four general patterns of staining. The first pattern is like Mdk, whereby the core of plaques is labeled most intensely, the second pattern, like Ptn, shows more intense peripheral staining of the plaque, a third pattern, such as seen with Sdc4, shows a very homogenous staining of the entire plaque and a fourth pattern, such as seen with Egfl8, shows patchy non-homogenous staining. Such data along with data showing that only a subset of parenchymal plaque associated proteins are detected in CAA, raises many questions about the nature of these interactions and the differences and similarities of not only CAA and plaque amyloid, but also even the concept that structural interactions may vary even within a single plaque. Direct studies of the human CAA proteome have also found increased levels of OLFML3, PTN and HTR1A, along with other known CAA components (e.g., APOE, CLU, APCS) ^92,93^. Further study of CAA and cerebral autosomal-dominant arteriopathy with subcortical infarcts and leukoencephalopathy (CADASIL) vessels’ proteomes shows overlap with each other, but also reveal extensive and broad overlap with many M42 DEPs ^92,94^. As Notch3 and Htra1 are increased DEPs in CRND8 mice, and we have been able to detect Htra1 in vessels, further study of CAA and its impact on the proteome is warranted. Indeed, there is emerging evidence that sequestration of HTRA1, a serine protease, with NOTCH3 in CADASIL leads to HTRA1 loss of function and accumulation of select HTRA1 substrates ^92,94^. Such data could account for the similarities between CADASIL and cerebral autosomal recessive arteriopathy with subcortical infarcts and leukoencephalopathy (CARASIL), the latter of which is caused by heterozygous loss of function mutations in *HTRA1*. Such data suggest that HTRA1 accumulation in CAA and in plaques might have similar impacts and contribute to CAA driven vascular dysfunction in AD.

The data showing that overexpression of Mdk or PTN in the CRND8 brain results in increased CAA and parenchymal Aβ amyloid suggests that these proteins have a direct impact on amyloid deposition, and therefore represent novel therapeutic targets in AD. As both proteins can accelerate aggregation of Aβ and bind preexisting amyloid fibrils *in vitro* we hypothesize they act as amyloid chaperones likely directly impacting the rate of Aβ aggregation. However, Mdk is known to have impacts on myeloid cell lineages and its co-accumulation in plaques could theoretically impact microglial mediated clearance either through gain or loss of function ^95,96^ Ptn, which signals largely through the same receptors as Mdk, could have similar impacts as well ^97^. Thus, it is also plausible that these proteins may also influence cell mediated clearance of Aβ. More broadly, additional studies will be needed to evaluate the impact of not only Mdk and Ptn but the many co-accumulating proteins on not amyloid deposition itself, but also the cellular responses to the deposits. Though a previous study had noted that a Mdk knockout crossed into an APP transgenic line resulted in increased amyloid deposition, we would note this data is simply anecdotal ^98^. Non-Tg mice Mdk and Ptn knockouts were also studied, and claimed to increase amyloid deposition, but in that study the antibody 6E10, which is highly selective for human Aβ was used to stain “plaques”. Thus, such results are likely artifactual.

HS and HSPGs along with chondroitin sulfate proteoglycans (CSPGs) have long been implicated in the biology of AD and other amyloid deposits ^99^. HS and select HSPGs core proteins (e.g., HSPG2, ACAN) have been shown to accumulate in plaques or dystrophic neurites. HSPGs have also been reported to accelerate Aβ aggregation and mediate internalization of extracellular tau ^100^. In these proteomic studies, we detect HSPG2, ACAN, and multiple CSPGs but they are minimally altered in either mice or the human AD proteome (Table S1). In contrast, in human AD, GPC4,5; SPOCK1-3; and SDC2,4 are much more robust DEPs and many of these are core M42 module members. In CRND8 mice Gpc1,5; Spock1-3 and Sdc2,4 are also robust DEPs (Table S1). Further, gene ontology strongly demonstrates heparin and glycosaminoglycan biology related biological processes upregulation in aged CRND8 mice brains (Figure 1).

We have not systematically evaluated heparin binding as a surrogate for HS/HSPG binding of many of the M42+ proteins studied here, but we find MDK and PTN secreted from mammalians cells bind robustly to heparin. We do find that a subset of the M42+ proteins secreted by human embryonic kidney 293 (HEK293) cells bind Aβ42 amyloid and a non-human, non-homologous amyloid fibrils ^36,37^. This subset of M42+ proteins was pragmatically chosen simply based on proteins which were secreted and detectable in the media of transiently transfected HEK293 cells. This data led us to explore whether MDK and PTN might be components of peripheral amyloids and we find both accumulate in TTR cardiac amyloid. Neither have been previously detected, even in extensive laser capture proteomic studies of cardiac amyloid deposits ^101^. As HS/HSPGs are common components of amyloids, these studies suggest that there may be very complex multipartite structural interactions between amyloid, HS/HSPGs and these co-accumulating proteins which may prove challenging to unravel.

Given the intensive studies of both Aβ amyloid and peripheral amyloids, it is somewhat surprising that we and others are finding numerous additional proteins co-accumulating with amyloid ^14,28,29,101,102^. In both the human and mouse studies brain tissue was extracted with 8M urea; thus, we are studying the total detectable urea solubilized proteome and not the detergent insoluble proteome. Indeed, for Mdk and Ptn in mice brains we find no gross change in solubility, despite the large increase in the Tg mouse brains. Following SDS-extraction, no residual protein is detected in the Urea fraction. These initial solubility studies likely explain why many of these proteins are not detected in detergent insoluble fractions typically used to isolate amyloid, they simply may not be there.

Our current data and data from others reveal numerous proteins that co-deposit with Aβ amyloid. Further, given the large number of proteins showing age progressive increases in the CRND8 mice brain, it is highly likely that over time many additional proteins will be found to accumulate with Aβ amyloid and perhaps other disease relevant amyloids as well. Indeed, two recently identified components of Aβ plaques, Pycard and Mfge8, are increased DEPs in our data, but in the 18M old CRND8 cohort rank 296th and 300^th^ among all the increased DEPs in terms of log2 Fc ^103,104^. In the human AD proteome, PYCARD is significantly increased (log2 Fc=0.06), and MFGE8 is detected but not increased. Cleary not all proteins upregulated will be components of plaques, but these data clearly point to biology of amyloid that is much more complex than previously appreciated, and that the response to amyloid deposits may also be equally as complex.

Despite thousands of studies and numerous hypotheses, there is ongoing debate as to how amyloid or Aβ aggregate pathology drives cellular dysfunction that leads to the neurodegenerative phase of AD ^2,7,8^. More generally, this debate extends to all amyloid disorders and how amyloid accumulation causes tissue damage ^34,41,105^. Indeed, extensive efforts have been made to identify more toxic species or conformers (e.g., oligomer, protofibril, dimer, specific fibrillar conformer) ^106^. Our current data, along with previous studies of proteins that co-accumulate with amyloid, offer an alternative hypothesis to the concept that amyloid or some distinct amyloid-like aggregate is a direct toxin ^70,107^. We term this the “amyloid scaffold” hypothesis (Figure S13) and posit that if amyloid deposits can scaffold the accumulation of numerous proteins than perhaps the accumulation of these proteins mediates downstream pathophysiology. Both toxic gain of function and loss of function mechanisms need to be considered. Accumulation of proteins within amyloid deposits could lead to gain of normal function if these proteins retain activity or if such binding activates a novel signaling pathway. Sequestration in the deposit could result in loss of function and reduced levels in the surrounding tissue. If solubility of select proteins accumulating in the amyloid deposit is not changed, then these proteins may retain signaling function. Indeed, many of the proteins we detect in plaques and CAA are known signaling molecules and through either loss or gain of function mechanisms their accumulation could impact dystrophic neurite and reactive glia in the local plaque microenvironment or cells in the vessel wall. In the heart MDK, though shown to have some cardioprotective effects, can also promote intimal hyperplasia whereas PTN have been shown to have angiogenic effects ^108-110^. Thus, this hypothesis may have relevance for peripheral amyloidoses as well. There are also intriguing parallels between this concept of amyloid as a scaffold for pathological process and functional amyloids which have been proposed to mediate their biological effects in part through a scaffolding mechanism (reviewed in ^111^).

As with many biologic responses, it is important to consider that the accumulation of proteins in the plaque is not simply a bystander effect, but part of the response to amyloid as a danger associated molecular pattern ^81,112^. The amyloid “scaffold” and amyloid (or oligomer) as “direct toxin” hypotheses are not mutually exclusive. Amyloid fibrils and oligomers are potent activators of microglia and myeloid cells and soluble oligomeric and protofibrillar aggregates can mediate many impacts on different cells ^30,70^. Proteins that interact with an amyloid or oligomer structure could be actively involved both in clearance, coating, and neutralization of the structure to reduce toxicity, or some combination of these. Other components may simply be bystanders, at least with respect to impacts on amyloid deposition, but once present could still mediate pathology by altering signaling gradients. er.

There is extensive data that APOE and CLU interaction with Aβ/Aβ amyloid deposits alters the local cellular response, and influence amyloid deposition itself^86-88,113,114^ Because of their genetic association with AD, they are also considered therapeutic targets, though it has been challenging to develop therapeutic approaches that modulate APOE or CLU and show robust preclinical disease modifying impacts ^115^. We have not yet explored studies in knockout mice, however, overexpression studies suggest that Mdk and PTN, like APOE and CLU, can increase amyloid deposition and both also increase CAA. It will be intriguing to evaluate the impacts of the larger set of proteins that are now being shown to co-accumulate. We might speculate that some will promote amyloid deposition, others might block it and still others might simply bind without altering kinetics of deposition. Our initial data suggest that many of these co-accumulating proteins could be considered as novel therapeutic targets, but additional probes of their biology will be necessary to determine their role in disease and their druggability.

Integrating data from genetic studies of AD and the DEPs in the CRND8 mouse brain, reinforces links between many established AD genes or prioritized genetic loci and AD risk genes and regulation of Aβ catabolism and deposition or response to Aβ deposition. This integration also provides an additional criterion that might be considered when prioritizing studies in loci with multiple candidate genes. Indeed, multiple proteins that are within M42, co-deposit with amyloid, or both and show nominally significant associations with AD. Integration of CRND8 proteome with AD CSF biomarkers shows that many proteins significantly altered in human AD CSF are DEPs in the brains of an amyloid depositing mouse model. Again, these data suggest that these CSF biomarkers might reflect the impacts of amyloid pathology on the AD brain. With the approval of immunotherapies that can dramatically impact amyloid deposition it will be intriguing to determine if this subset of CSF AD biomarkers is altered in response to such therapy. Recently SMOC1 and SPON1, two core M42 members have been found to be altered in concert with decreases in Aβ42/40 ratios in the CSF of individuals with dominantly inherited AD ^57^. Further, Smoc1 is a highly reproducible highly upregulated AD CSF biomarker ^53,57,61,116^.

In summary, this contextualized understanding of AD and APP mouse model proteomic changes provides insights into the true molecular complexity of AD pathogenesis and informs on biomarkers and novel therapeutic targets related to amyloid deposition. Though we have i) validated some of the more robust protein changes in the proteomic analyses, ii) provided pathological insight related to a subset of these proteins, and iii) generated initial modeling and biochemical data to support potential pathobiological roles, there is still much to be learned about amyloid and its role in AD pathophysiology and in other amyloid disorders. Clearly, future integrative omic studies can build off and refine these current studies, especially by incorporating models of additional pathologies relevant to AD and other amyloid disorders.

### Limitations of the study

We commented on the many limitations of this multifaceted study in the results and discussion, but several overarching limitations are highlighted here. First, the pathological validation studies vary with respect to the confidence in the findings. Classic criteria suggest two antibodies or antisera to two distinct epitopes of the target protein are needed to confirm presence in pathological setting. A subset of the pathological findings is based on a single antibody or antisera. However, we would note that this criterion was often utilized in an era when additional data (e.g., proteomic based) was lacking. Second, we do not have extensive proteomic data from human brains with amyloid deposition prior to the presence of other pathologies. Thus, we have limited our integration of the mouse data to human brain data which has a spectrum of AD pathologies. This fact limits inference as to which proteomic changes in humans are truly amyloid driven. Third, the lack of analyses of a group young enough to lack amyloid deposition or alternatively a manipulation that alters amyloid loads, impose some interpretative limitations on the data with respect to changes driven by amyloid or mutant APP overexpression. Nevertheless, given the multiple layers of orthogonal and integrative data used in this study we believe that this study begins to capture and contextualize proteomic changes in AD and a mouse model of amyloid deposition in a way that will inform downstream studies for many years.

## METHODS

### Cloning and cell culture

All cDNAs used were synthesized by Genscript (Piscataway, NJ) and subcloned into pCDNA3.1 vector, the rAAV2-CBA expression vector pCTR4 ^117^, or both. All cDNAs incorporated a FLAG epitope tag (DYKDDDDK) on the-COOH terminal. To overexpress the genes, 2.7 µg plasmid were transiently transfected into HEK 293T cells in a 6-well dish using polyethylenimine (PEI). Media was replaced with FBS free Opti-MEM after 18 hr. incubation. Cells and conditioned media were collected after another 24 hr. incubation. Bis-Tris precast gels (Bio-Rad, Hercules, CA) were used for all SDS-PAGE. 50 µg cell lysate by total protein and 30 µl media of each sample were loaded. Western blot was developed using monoclonal anti-FLAG M2 antibody and anti-β-actin (Sigma-Aldrich, Inc, St. Louis, MO).

### Animal models and neonatal injections

All animal procedures were approved by the Institutional Animal Care and Use Committee in accordance with NIH guidelines. CRND8 mice were bred in house and housed three to five to a cage and maintained on ad libitum food and water with a 12hr light/dark cycle√. Intracerebroventricular injections of recombinant adeno-associated virus (rAAVs) were carried out on day P0 as described previously ^118^. Two microliters of rAAV2/1 encoding Mdk and PTN was administrated bilaterally into the cerebral ventricle. Mice injected with rAAV encoding MDK and PTN were aged 3 or 6 months and euthanized, brains were harvested and one hemibrain was fixed overnight in 4% paraformaldehyde at 4°C followed by processing and paraffin embedding for immunohistochemical staining. The other hemibrain was snap frozen in isopentane on dry-ice and then stored at −80°C until it was thawed and homogenized for ELISA measurements of Aβ peptide levels.

### Mouse Brain Histology and immunohistochemistry

Paraffin sections (5 μm) were used for all histology and immunohistochemistry studies. For various antibodies screens two hemibrains from the same mouse were fixed in EtOH or in 4% PFA (alternatively, in 10% Formalin) and embedded into the same block. Antibodies used are listed in Table S11. The slides were scanned by Aperio XT System (Leica Biosystems, Buffalo Grove, IL). Thio-S tissue staining methods were performed as previously described ^119^. For studies in mice injected with pAAV-Mdk or pAAV-PTN, embedded sections were immunohistochemically stained with a biotinylated pan-Aβ antibody Ab5 ^120^ (1:500) or anti-FLAG antibody (Sigma-Aldrich Inc, St. Louis, IL) and developed using Vectastain Elite ABC Kit (Vector Laboratories, Burlingame, CA) followed with 3,3’- diaminobenzidine (DAB) substrate (Vector Laboratories, Burlingame, CA) and counterstained with hematoxylin. The slides were scanned by Aperio XT System and analyzed using the ImageScope program. In brief, at least three sections per sample, at least 30 µm apart, were imaged and plaque burden was quantified. For three-month old mice, plaque number was calculated by two independent observers.

### Human Tissue Histology and immunohistochemistry

5 μm thick sections of formalin fixed or ethanol fixed, paraffin embedded postmortem brain tissue was deparaffinized in xylene twice for 5 minutes and followed by rehydration in descending ethanol series (100%, 100%, 90%, 70%) for 1 minute each. For Formic acid antigen retrieval, sections were immersed in a 70% Formic acid solution for 20 minutes (where indicated in Table S12). Sections were incubated in 0.1M Tris and 0.005% Tween (or Citrate buffer, where indicated in table below) at high pressure in a pressure cooker for 15 minutes, followed by incubation in a PBS/H2O2 solution with 10% Triton-x for 20 minutes to quench endogenous peroxidase. The sections were washed with tap water and then equilibrated to 0.1M Tris for 5 min before blocking in normal horse serum for 20 minutes, followed by blocking in 2% FBS/0.1 M Tris, (pH 7.6) for 5 minutes. After that, primary antibody, diluted in 2% FBS/0.1 M Tris, (pH 7.6) was applied to the section and incubated overnight at 4 °C. Dilutions for each antiserum is listed in Table S11. The next day, sections were quickly rinsed in 0.1M Tris and blocked in 2% FBS/0.1 M Tris, (pH 7.6) for 5 minutes before incubating with the secondary antibody (HRP-conjugated ImmPRESS Polymer Reagent, Vector Labs) for 30 minutes at room temperature. Following a quick wash in 0.1M Tris, the signal was developed using 3,3’-diaminobenzidin (DAB, Vector Lab SK-410) for 1-2 minutes and then sections were counterstained with hematoxylin (Mayer’s version, Sigma Aldrich) for 1 minute. Next, sections were washed in tap water, and then dehydrated in an ascending series of ethanol (70%, 90%, 100%, 100%) for 1 minute each, followed by xylene (2 x 5 mins). Sections were cover slipped using Cytoseal 60 (Thermo Fisher) mounting media and dried overnight.

### Aβ ELISAs

After tissue harvesting, the left hemisphere was flash-frozen in isopentane. The frozen cortex was sequentially extracted with protease inhibitor cocktail (Roche, Basel, Switzerland) containing RIPA buffer, 2% SDS, and 70% formic acid (FA) as described previously at a concentration of 150 mg/ml. Aβ levels from the 2% SDS– and 70% FA–extracted samples were quantified using end-specific sandwich ELISA as previously described ^121^. Aβ40 was captured with mAb 13.1.1 and detected by HRP-conjugated mAb5. Aβ42 was captured with mAb 2.1.3 and detected by HRP-conjugated mAb5. ELISA results were analyzed using SoftMax Pro software (Molecular Devices, San Jose, CA).

### Amyloid-β Aggregation Assay

The effect of midkine (MDK) and pleiotrophin (PTN) proteins on amyloid beta 1-42 (Aβ1-42) aggregation was measured by *in vitro* thioflavin T (ThT) fluorescence assay, as previously described ^122^. Recombinant human Aβ1-42 (20 ng/µL equivalent to 5 µM) from rPeptide (A-1170-1) was incubated in Tris-buffered Saline (TBS; 150 mM NaCl, 50 mM Tris-HCl, pH 7.6), and 20 µM ThT in the presence or absence of purified recombinant MDK (1.25 ng/µl equivalent to 47 nM) from Sino Biological (10247-HNAB) or PTN (5 ng/µl equivalent to 327 nM) from R&D Systems (R&D252-PL-050). The final volume within each well was 100 µL. The assay was conducted in quadruplicates using chilled (4°C) 96 well black clear bottom plates (Corning, #3904). Fluorescence was captured at 420 Ex, 480 Em for 20 hours at 15 min intervals at 37°C using Synergy H1 (Biotek) microplate reader. ThT alone (20 µM) was measured and subtracted as background fluorescence. A titration curve was also performed with fixed concentration of Aβ1-42 (5 µM) and an increasing concentration of MDK (12 nM, 23 nM, 47 nM and 93 nM) or PTN (82 nM 163 nM and 327 nM). Fluorescence intensities were graphed using GraphPad Prism.

### Pulldowns and Western Blot Analysis

Aβ1-42 fibrils prepared as described above (20 hr. incubation) were precipitated by centrifugation at 10,000 x g. The pellet was resuspended in 8M urea buffer (8 M urea, 100 mm NaHPO4, pH 8.5) and boiled in Laemmli sample buffer (BioRad, 161-0737) at 98°C for 5 min. Proteins were resolved on Bolt 4-12% Bis-Tris gels (Thermo Fisher Scientific, NW04120BOX) followed by transfer to nitrocellulose membrane using iBlot 2 dry blotting system (ThermoFisher Scientific, IB21001). Membranes were incubated with StartingBlock buffer (ThermoFisher, 37543) for 30 min followed by overnight incubation at 4° in primary antibodies including Aβ (Novus, NBP11-97929), MDK (Abcam, #52637), and PTN (Abcam, #ab79411) antibodies. Membranes were washed with TBS containing 0.1% (v/v) Tween 20 (TBS-T) and incubated with fluorophore-conjugated secondary antibodies (Alexa Fluor-680 or Alexa Fluor-800; Invitrogen) for 1 hr. at room temperature. Membranes were subsequently washed three times with TBS-T and images were captured using an Odyssey Infrared Imaging System (LI-COR Biosciences).

### Statistical analysis

Data were analyzed statistically according to the methods specified in each figure legend. Briefly, p values were obtained as specified by either a one-way ANOVA test with two groups, or for enrichment, by a Fisher’s Exact test. P values were corrected for multiple testing were indicated by Benjamini-Hochberg FDR correction. For significance of Pearson correlations, Student’s test was used.

### Proteomic Sample Processing

Each tissue piece was individually homogenized in 300 uL of urea lysis buffer (8M urea, 100 mM NaHPO4, pH 8.5), including 3 uL (100x stock) HALT protease and phosphatase inhibitor cocktail (Pierce). All homogenization was performed using a Bullet Blender (Next Advance) according to manufacturer protocols. Briefly, each tissue piece was added to Urea lysis buffer in a 1.5 mL Rino tube (Next Advance) harboring 500 mg stainless steel beads (0.9-2 mm in diameter) and blended twice for 5-minute intervals in the cold room (4°C). Protein supernatants were transferred to 1.5 mL Eppendorf tubes and sonicated (Sonic Dismembrator, Fisher Scientific) 3 times for 5 s with 15 s intervals of rest at 30% amplitude to disrupt nucleic acids and subsequently vortexed. Protein concentration was determined by the bicinchoninic acid (BCA) method, and samples were frozen in aliquots at −80°C. Protein homogenates (100 ug) were diluted with 50 mM NH4HCO3 to a final concentration of less than 2M urea and then treated with 1 mM dithiothreitol (DTT) at 25°C for 30 minutes, followed by 5 mM iodoacetimide (IAA) at 25°C for 30 minutes in the dark. Protein was digested with 1:100 (w/w) lysyl endopeptidase (Wako) at 25°C for 2 hours and further digested overnight with 1:50 (w/w) trypsin (Promega) at 25°C. Resulting peptides were desalted with a Sep-Pak C18 column (Waters) and dried under vacuum.

### Tandem Mass Tag (TMT) Labeling

Batches 1 and 2 were labeled using the TMT 11-plex kit (ThermoFisher 90406 – Lot# TG273545 and TG273555) while batch 3 was labeled with TMTPro (ThermoFisher A44520 – Lot# VH311511) according to manufacturer’s protocol. For batches 1 and 2, each sample (containing 100 μg of peptides) was re-suspended in 100 mM TEAB buffer (100 μL). The TMT labeling reagents were equilibrated to room temperature, and anhydrous ACN (256 μL) was added to each reagent channel. Each channel was gently vortexed for 5 min, and then 41 μL from each TMT channel was transferred to the peptide solutions and allowed to incubate for 1 h at room temperature. The reaction was quenched with 5% (vol/vol) hydroxylamine (8 μL) (Pierce). For batch 3, each sample (containing 100 μg of peptides) was re-suspended in 100 mM TEAB buffer (100 μL). The TMT labeling reagents were equilibrated to room temperature, and anhydrous ACN (200 μL) was added to each reagent channel. Each channel was gently vortexed for 5 min, and then 20 μL from each TMT channel was transferred to the peptide solutions and allowed to incubate for 1 h at room temperature. All channels for each batch were then combined (batch 1 used all 11 channels, batch 2 utilized only 10 channel and batch 3 utilized only 8 channels) and dried by SpeedVac (LabConco) to approximately 150 μL and diluted with 1 mL of 0.1% (vol/vol) TFA, then acidified to a final concentration of 1% (vol/vol) FA and 0.1% (vol/vol) TFA. Peptides were desalted with a 200 mg C18 Sep-Pak column (Waters). Each Sep-Pak column was activated with 3 mL of methanol, washed with 3 mL of 50% (vol/vol) ACN, and equilibrated with 2×3 mL of 0.1% TFA. The samples were then loaded, and each column was washed with 2×3 mL 0.1% (vol/vol) TFA, followed by 2 mL of 1% (vol/vol) FA. Elution was performed with 2 volumes of 1.5 mL 50% (vol/vol) ACN. The eluates were then dried to completeness.

### High pH Fractionation

For batches 1 and 2, dried samples were re-suspended in high pH loading buffer (0.07% vol/vol NH4OH, 0.045% vol/vol FA, 2% vol/vol ACN) and loaded onto an Agilent ZORBAX 300 Extend-C18 column (2.1mm x 150 mm with 3.5 µm beads). An Agilent 1100 HPLC system was used to carry out the fractionation. Solvent A consisted of 0.0175% (vol/vol) NH4OH, 0.01125% (vol/vol) FA, and 2% (vol/vol) ACN; solvent B consisted of 0.0175% (vol/vol) NH4OH, 0.01125% (vol/vol) FA, and 90% (vol/vol) ACN. The sample elution was performed over a 58.6 min gradient with a flow rate of 0.4 mL/min. The gradient consisted of 100% solvent A for 2 min, then 0% to 12% solvent B over 6 min, then 12% to 40 % over 28 min, then 40% to 44% over 4 min, then 44% to 60% over 5 min, and then held constant at 60% solvent B for 13.6 min. A total of 96 individual equal volume fractions were collected across the gradient and subsequently pooled by concatenation into 24 fractions. For batch 3, dried samples were re-suspended in high pH loading buffer (0.07% vol/vol NH4OH, 0.045% vol/vol FA, 2% vol/vol ACN) and loaded onto a Waters BEH 1.7 um 2.1mm by 150mm. A Thermo Vanquish was used to carry out the fractionation. Solvent A consisted of 0.0175% (vol/vol) NH4OH, 0.01125% (vol/vol) FA, and 2% (vol/vol) ACN; solvent B consisted of 0.0175% (vol/vol) NH4OH, 0.01125% (vol/vol) FA, and 90% (vol/vol) ACN. The sample elution was performed over a 25 min gradient with a flow rate of 0.6 mL/min. A total of 192 individual equal volume fractions were collected across the gradient and subsequently pooled by concatenation into 96 fractions. All fractions were dried to completeness using a SpeedVac.

### Liquid Chromatography Tandem Mass Spectrometry

For batches 1 and 2, each of the 24 high-pH peptide fractions was resuspended in loading buffer (0.1% FA, 0.03% TFA, 1% ACN). Peptide eluents were separated on a self-packed C18 (1.9 µm Dr. Maisch, Germany) fused silica column (50 cm × 75μM internal diameter (ID), New Objective, Woburn, MA) by an Easy nLC 1200 (Thermo Scientific) and monitored on a Q-Exactive HFX mass spectrometer (Thermo Scientific). Elution was performed over a 120-min gradient at a rate of 250 nL/min or 300 nL/min with buffer B ranging from 3% to 50% (buffer A: 0.1% FA in water, buffer B: 0.1 % FA in 80% ACN). For batch 1, the mass spectrometer was set to acquire data in positive ion mode using data-dependent acquisition with top 20 cycles. Each cycle consisted of one full MS scan followed by a maximum of 20 MS/MS. Full MS scans were collected at a resolution of 120,000 (400-1600 m/z range, 1×10^6 AGC, 100 ms maximum ion injection time). All higher energy collision-induced dissociation (HCD) MS/MS spectra were acquired at a resolution of 45,000 (1.6 m/z isolation width, 0.5 m/z offset, 32% collision energy, 1×10^4 AGC target, 86 ms maximum ion time). Dynamic exclusion was set to exclude previously sequenced peaks for 20 seconds within a 10-ppm isolation window. For batch 2, the mass spectrometer was set to acquire data in positive ion mode using data-dependent acquisition with top 10 cycles. Each cycle consisted of one full MS scan followed by a maximum of 10 MS/MS. Full MS scans were collected at a resolution of 120,000 (400-1600 m/z range, 3×10^6 AGC, 100 ms maximum ion injection time). All higher energy collision-induced dissociation (HCD) MS/MS spectra were acquired at a resolution of 45,000 (1.6 m/z isolation width, 0.5 m/z offset, 30% collision energy, 1×10^4 AGC target, 86 ms maximum ion time). Dynamic exclusion was set to exclude previously sequenced peaks for 20 seconds within a 10-ppm isolation window.

For batch 3, each of the 96 high-pH fractions was resuspended in loading buffer (0.1% FA, 0.03% TFA, 1% ACN). Peptide eluents were separated on a self-packed C18 (1.7 µm Water’s BEH) fused silica column (laser pulled 15 cm × 150μM ID) by Ultimate 3000 RSLCnano (Thermo Scientific). Elution was performed over a 36-min gradient at a rate of 1ul/min with buffer B ranging from 3% to 50% (buffer A: 0.1% FA in water, buffer B: 0.1 % FA in 80% ACN). Mass spectrometry was performed with a high-field asymmetric waveform ion mobility spectrometry (FAIMS) Pro frontend equipped Orbitrap Eclipse (Thermo) in positive ion mode using data-dependent acquisition with 1.5 second top speed cycles for each FAIMS compensative voltage (CV). Each cycle consisted of one full MS scan followed by as many MS/MS events that could fit within the given 1.5 second cycle time limit. MS scans were collected at a resolution of 120,000 (410-1600 m/z range, 4×10^5 AGC, 50 ms maximum ion injection time, FAIMS CV of −45 and −65). Only precursors with charge states between 2+ and 6+ were selected for MS/MS. All higher energy collision-induced dissociation (HCD) MS/MS spectra were acquired at a resolution of 30,000 (0.7 m/z isolation width, 35% collision energy, 1.25^5 AGC target, 54 ms maximum ion time, turboTMT on). Dynamic exclusion was set to exclude previously sequenced peaks for 20 seconds within a 10-ppm isolation window.

### Proteome Identification and Quantification with Proteome Discoverer

All raw files were analyzed using Proteome Discoverer v2.5.0.400 (ThermoFisher Scientific). MS/MS spectra were compared to theoretical spectra in a search performed by the Sequest HT search engine, with parameters specified as: fully tryptic specificity, a maximum of two missed cleavages, a minimum peptide length of six, fixed modifications for TMT or TMTPro tags on lysine residues and peptide N-termini (respectively +229.163 or +304.207 Da), carbamidomethylation of cysteine (+57.021 Da), and dynamic modification of methionine oxidation (+15.995 Da) and N-terminal protein loss of methionine, methionine loss plus acetylation, or acetylation (−131.040 Da, −89.030 Da, or +42.011 Da, respectively); precursor mass tolerance was 20 ppm, and Fragment tolerance 0.05 Da. The FASTA database searched was downloaded from UniProt on August 15, 2020, and contained 91,413 entries, plus one added for human amyloid beta residues 6-28 (HDSGYEVHHQKLVFFAEDVGSNK). Correction for stable isotope labeling impurity of the TMT tags used in each of the 3 batches was performed. Proteins reported in the consensus workflow were filtered by Percolator to 0.005 FDR-passed 1,266,002 PSMs, 0.01 FDR-passed 310,145 peptide groups, and 0.05 FDR-passed 49,656 proteins, including decoys in each case. Strict parsimony principles were followed to group peptides into proteins so that 11,681 protein parsimony groups represent the full results. A complete protein-level TMT reporter normalized abundance table is available in Table S3.

### Bioinformatic Analyses

Circular heatmap plot of eigenprotein-trait correlations (bicor), native eigenprotein −log10(p) for signed significance of AD vs control or synthetic eigenprotein Tg vs WT comparison, and the −log10(BH FDR) of cell type marker enrichment from the one-tailed Fisher’s test p value was visualized using the R circlize package suite of functions using an in-house script. Synthetic eigenprotein calculations used the top 20 percent of hubs ranked by kME, a minimum of 4 hubs per module, calculating the first principal component of variance in those hubs ^9^ after mapping cross-species using the biomaRt R package getLDS function. Effect size correlation plots leveraged the WGCNA verboseScatterplot function for Pearson rho, least squares fit line, and Student’s significance of the Pearson correlation. Boxplots were drawn using the base R boxplot function, with individual points overlain using the beeswarm package function of the same name. The stacked bar plot of percent differentially expressed proteins per module was executed using the DEXpercentStacked function available from https://www.github.com/edammer/parANOVA/. Ensemble human AD GWAS MAGMA p values (Table S1) were determined as the mean −log10(p) gene-level risk for all genes reaching nominal significance in any of the three GWAS studies considered ^43-45^, following rollup of SNP-level GWAS summary statistics to the gene-level p value using MAGMA v1.09b.

### Gene Ontology Analyses

Gene ontology enrichment analysis were performed with goseq ^123^ to identify enrichment in gene ontology categories and KEGG pathways. For DEGs, up- and down-regulated gene lists will be analyzed separately. For WGCNA, gene lists from each module will be used as input. Over-represented P-values will be adjusted for multiple comparisons using the Benjamini–Hochberg (BH) adjustments for controlling false-discovery rates. An enrichment score will be calculated using an observed-over-expected ratio for each gene list. GO-BP categories and KEGG pathways were plotted if their BH adjusted FDR reached >= 0.05 and the number of DEGs within each category/pathway was greater than 4. Individual GO terms were grouped based on their similarity of gene composition computed using the Jaccard similarity index where J(A,B) = |size of A intersection B| / |size of A union B|. These groupings were then expert-annotated based on the overarching main function and theme of their constituent members into supergroups. The mean enrichment and p-values for the union of all member GO terms for each supergroup were calculated for graphing.

### RNA extraction, sequencing and analyses for 18M mouse brains

RNA sequencing data for TgCRND8 transgenic mice was downloaded from Synapse (https://doi.org/10.7303/syn3157182). RNA was extracted using the RNeasy mini extraction kit with on-column DNase treatment (QIAGEN). RNA quantity is determined with the Qubit RNA HS assay. RNA quality will be checked via the RNA Integrity Number (RIN) on an Agilent Bioanalyzer 2100 with the Eukaryote Total RNA Nano chip. Libraries will be generated polyA enrichment using the Illumina TruSeq Stranded mRNA library prep kit. Libraries will be sequenced on paired-end, 100 bp runs on the Nextseq 2000 (Illumina) utilizing a pooling strategy that minimized batch effects from extraction, library preparation.

IFASTQ files were aligned against the mouse genome (GRCm39) and GRCm39.107 annotation using STAR^124^ to generate BAM files. Gene counts were generated from BAM files using Rsamtools (https://bioconductor.org/packages/release/bioc/html/SummarizedExperiment.html) and the summarizeOverlaps function with the GenomicAlignments package v1.36.0 ^125^. Differential gene expression analysis was performed with DESeq2 package v1.40.2 using the “DESeq” function with default settings ^126^ which fits a generalized linear model for each gene. Subsequent Wald test P-values are adjusted for multiple comparisons using the Benjamini–Hochberg method (adjusted P-value). Pair-wise changes in gene expression levels between groups were used to identify DEGs. DEGs will be defined as an absolute log2 fold change ≥0.5 and an adjusted P-value ≤0.05.IFASTQ files were aligned against the mouse genome (GRCm39) and GRCm39.107 annotation using STAR ^124^ to generate BAM files. Gene counts were generated from BAM files using Rsamtools (https://bioconductor.org/packages/release/bioc/html/SummarizedExperiment.html) and the summarizeOverlaps function with the GenomicAlignments package v1.36.0 ^125^. Differential gene expression analysis was performed with DESeq2 package v1.40.2 using the “DESeq” function with default settings ^126^ which fits a generalized linear model for each gene. Subsequent Wald test P-values are adjusted for multiple comparisons using the Benjamini–Hochberg method (adjusted P-value). Pair-wise changes in gene expression levels between groups were used to identify DEGs.

### AMP-AD brain transcriptome and proteomic datasets

Brain transcriptome dataset from Mayo Clinic on the AD Knowledge Portal was utilized for human vs mouse DEG comparisons). Human and CRND8 mouse brain RNA sequencing data was downloaded from Synapse (doi: https://doi.org/10.7303/syn5550404; https://www.synapse.org/#!Synapse:syn17008858). Mayo RNAseq dataset comprises transcriptome measures from temporal cortex (TCX, superior temporal gyrus) and cerebellum (CER) the former of which was utilized in this study. For Non-Tg mice and CRND8 mice a sagitally dissected whole hemibrain was used. RNA isolation, data collection, sequencing alignment, counting and QC has been described in detail elsewhere ^12,30,76^. EdgeR was used to reprocess both the mouse and human RNA sequencing data ^127^. Mouse and human orthologues were identified in a two-step process. Most orthologous genes were first identified using http://alliancegenome.org/downloads, Database Version: 5.4.0 on Apr 20, 2023, with “Orthology Filter: Stringent”. Then a manual curation of additional orthologues was conducted using the gOrth serac function on the g:profiler website (https://biit.cs.ut.ee/gprofiler/orth). Human brain proteomic data was utilized form previous publications. The human proteome data is available at Synapse (https://www.synapse.org/#!Synapse:syn20933797/wiki/596247).

## Availability of data and materials

Raw mass spectrometry data and pre- and post-processed plasma protein abundance data and case traits related to this manuscript are available at https://synapse.org/CRND8. The results published here are in whole or in part based on data obtained from the AMP-AD Knowledge Portal (https://adknowledgeportal.synapse.org). The AMP-AD Knowledge Portal is a platform for accessing data, analyses and tools generated by the AMP-AD Target Discovery Program and other programs supported by the National Institute on Aging to enable open-science practices and accelerate translational learning. The data, analyses and tools are shared early in the research cycle without a publication embargo on secondary use. Data are available for general research use according to the following requirements for data access and data attribution (https://adknowledgeportal.synapse.org/#/DataAccess/Instructions).

## AD Knowledge Portal: AMP-AD datasets

The results published here are in whole or in part based on data obtained from the AMP-AD Knowledge Portal (https://doi.org/10.7303/syn2580853). Mayo Clinic: The Mayo RNAseq study data was led by Dr. Nilüfer Ertekin-Taner, Mayo Clinic, Jacksonville, FL as part of the multi-PI U01 AG046139 (MPIs Golde, Ertekin-Taner, Younkin, Price). Samples were provided from the following sources: The Mayo Clinic Brain Bank and Banner Sun Health Research Institute. Data collection was supported through funding by NIA grants P50 AG016574, R01 AG032990, U01 AG046139, R01 AG018023, U01 AG006576, U01 AG006786, R01 AG025711, R01 AG017216, R01 AG003949, NINDS grant R01 NS080820, CurePSP Foundation, and support from Mayo Foundation. Study data includes samples collected through the Sun Health Research Institute Brain and Body Donation Program of Sun City, Arizona. The Brain and Body Donation Program is supported by the National Institute of Neurological Disorders and Stroke (U24 NS072026 National Brain and Tissue Resource for Parkinson’s Disease and Related Disorders), the National Institute on Aging (P30 AG19610 Arizona Alzheimer’s Disease Core Center), the Arizona Department of Health Services (contract 211002, Arizona Alzheimer’s Research Center), the Arizona Biomedical Research Commission (contracts 4001, 0011, 05-901 and 1001 to the Arizona Parkinson’s Disease Consortium) and the Michael J. Fox Foundation for Parkinson’s Research. For the human protein data All raw data, case traits and analyses (differential and co-expression) related to this manuscript are available at https://www.synapse.org/consensus. The results published here are in whole or in part based on data obtained from the AMP-AD Knowledge Portal (https://adknowledgeportal.synapse.org). The AMP-AD Knowledge Portal is a platform for accessing data, analyses, and tools generated by the Accelerating Medicines Partnership (AMP-AD) Target Discovery Program and other National Institute on Aging (NIA)-supported programs to enable open-science practices and accelerate translational learning. The data, analyses and tools are shared early in the research cycle without a publication embargo on secondary use. Data is available for general research use according to the following requirements for data access and data attribution (https://adknowledgeportal.synapse.org/#/DataAccess/Instructions). ROSMAP resources can be requested at www.radc.rush.edu.

## Author contributions

Conceptualization, TEG, NTS, SP, AIL, YL, ECBJ, ED; Methodology, TEG, YL, YR, DD, JG, NTS, MA, KD, DR, JT, CM, LP, AI, AE, BDM, KDD, AN, TI, FA, TL, KM, BM, MR, CF; Investigation, YL, EBD, YR, WT, DD, MA, JG, KL, JT-L, JP, AI, AE, BM, KM, AN; Formal Analysis, YL, KM, EBD, MR, CF, NTS, TEG, AN; Writing – Original Draft, TEG, YL, SP; Writing – Review & Editing, YL, EBD, DD, NET, JWK, AIL, KM, NTS, TEG, SP, FLH; Funding Acquisition, TEG, SP, NTS, YL AIL and; Resources, AIL, JWK, NTS, SP, TEG, ECBJ, FLH, JWK, JJ; All authors read and approved the final manuscript.

## Supplemental material

### Figure legends

**Figure S1.**
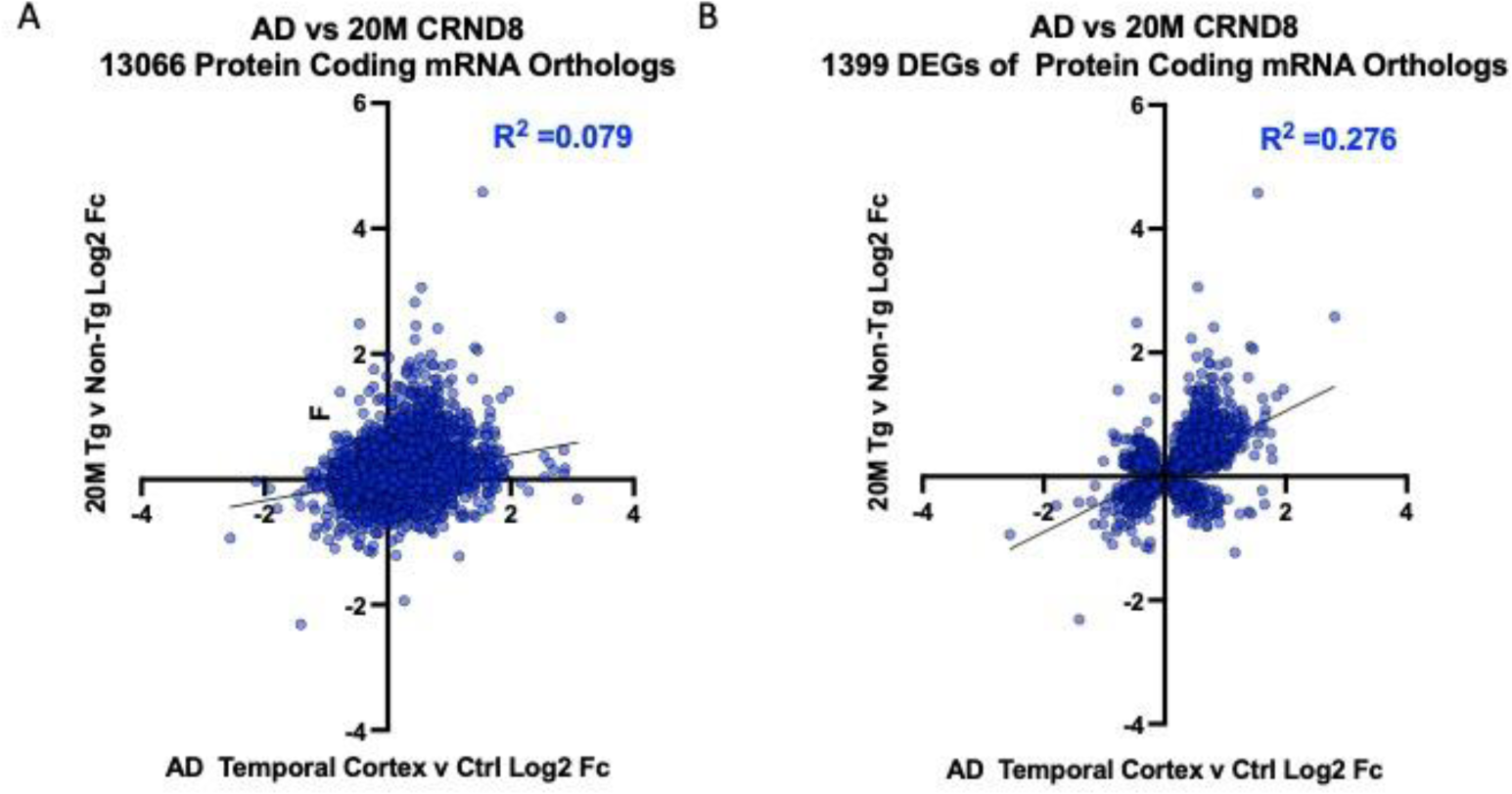
Correlation of protein encoding mRNAs in 20M CRND8 mice (Tg versus non-Tg) and AD versus Control human Temporal Cortex. Previously generated RNA expression data from human temporal cortex ^12^ and CRND8 mice brains ^30^ was reanalyzed using EdgeR and filtered for orthologues. (A) Plot of log2FC of human temporal cortex AD versus Control mRNA levels (x-axis) and 20M CRND8 of orthologous protein coding RNA (y-axis). (B) Plot of log2FC of human temporal cortex AD versus Control mRNA levels (x-axis) and 20M CRND8 of shared DEGs of protein coding genes (y-axis). Data used to generate these plots is found in Table S6.

**Figure S2.**
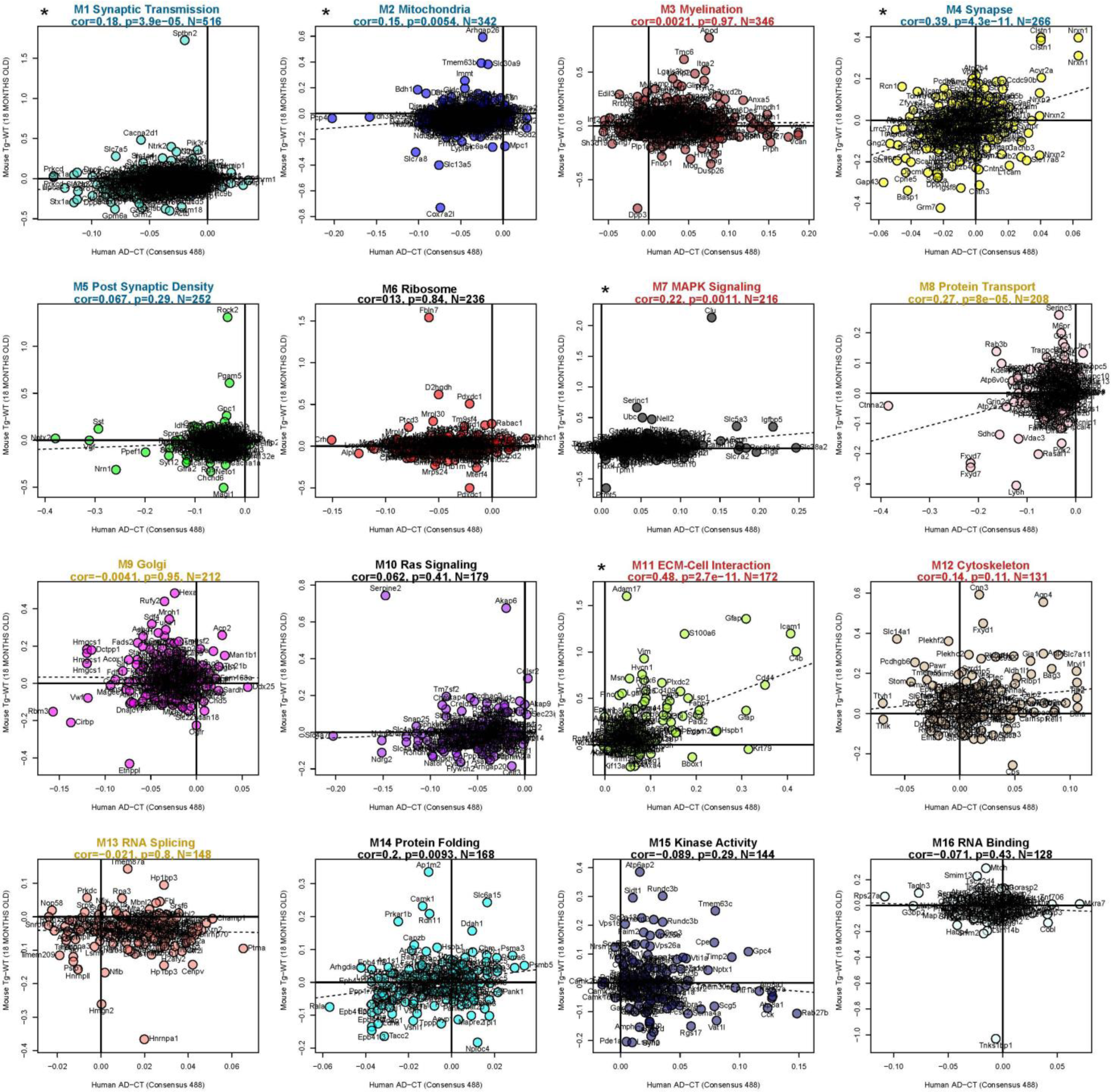
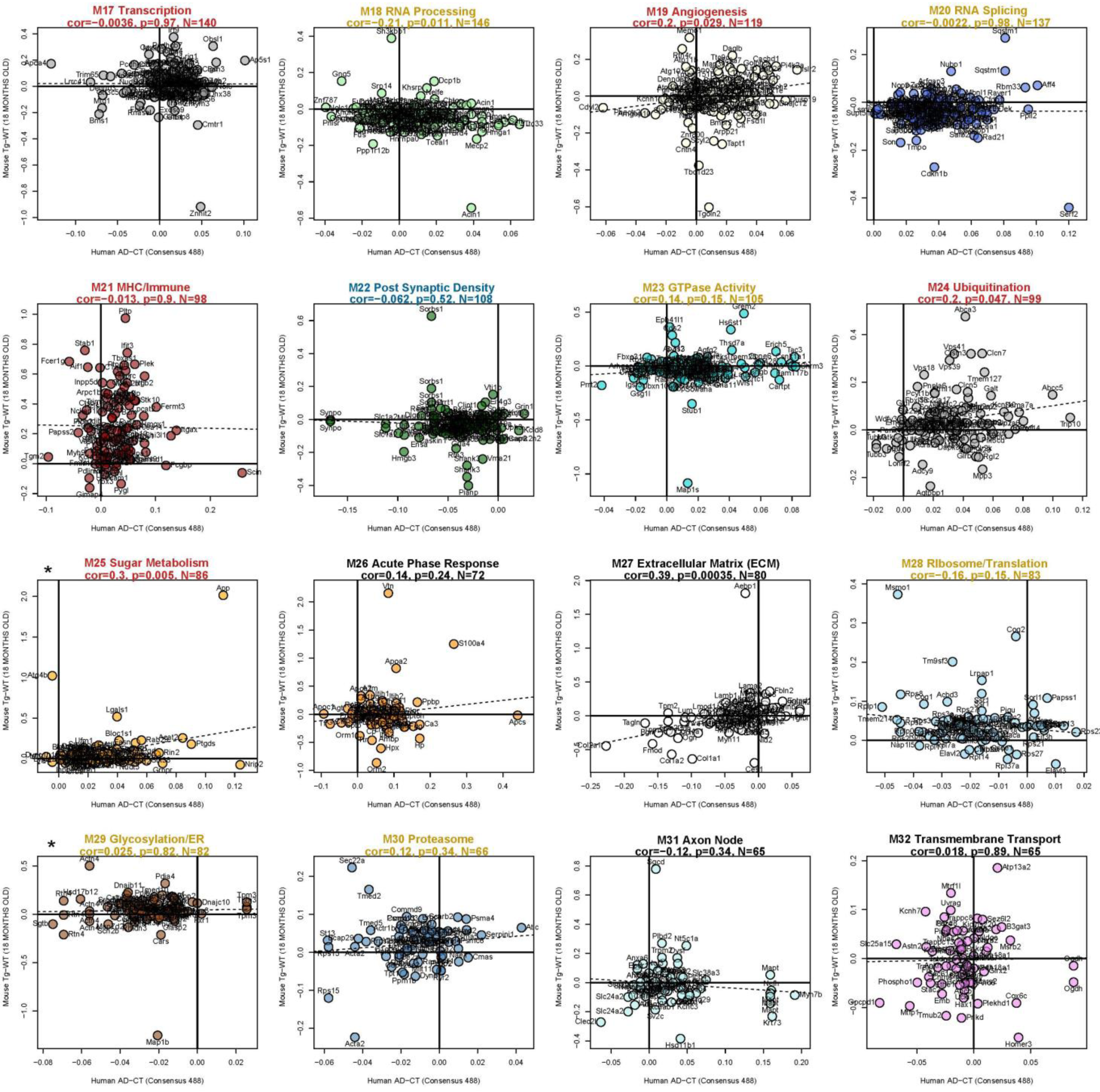
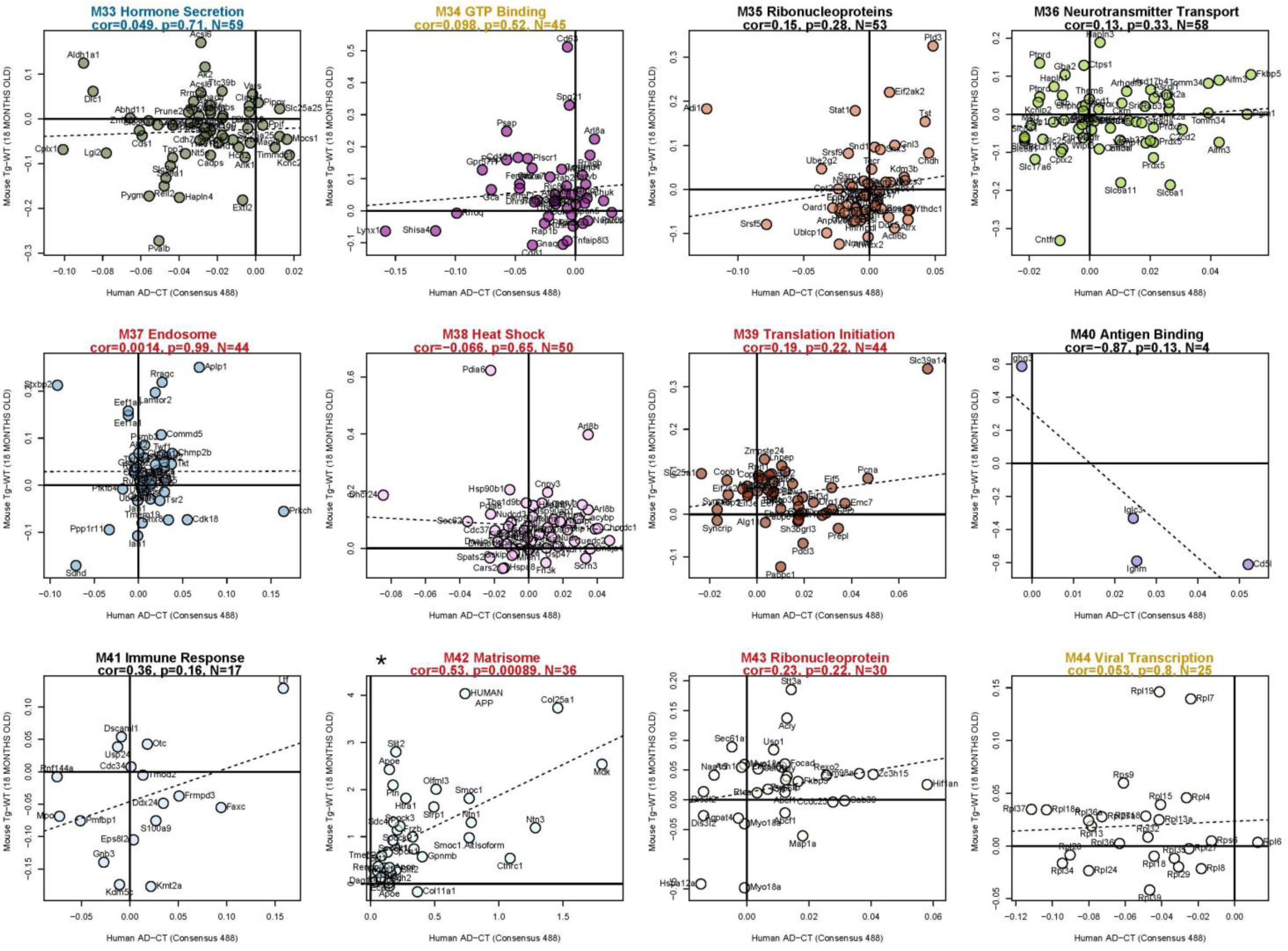
Effect size correlation for human AD vs control effect and mouse Tg effect at 18M shows varying concordance in each of the 44 consensus brain modules. Panels are in rank size order from M1 to M44, with each title providing module number, description, Pearson correlation (rho), Student’s significance of correlation, number of mouse orthologs mapped to the members of the human module, and all points labeled by the gene symbol of the mouse ortholog. *Modules also displayed in Figure 2 are labeled with an asterisk. Text color of each module plot title matches the scheme described in Figure 2F.

**Figure S3.**
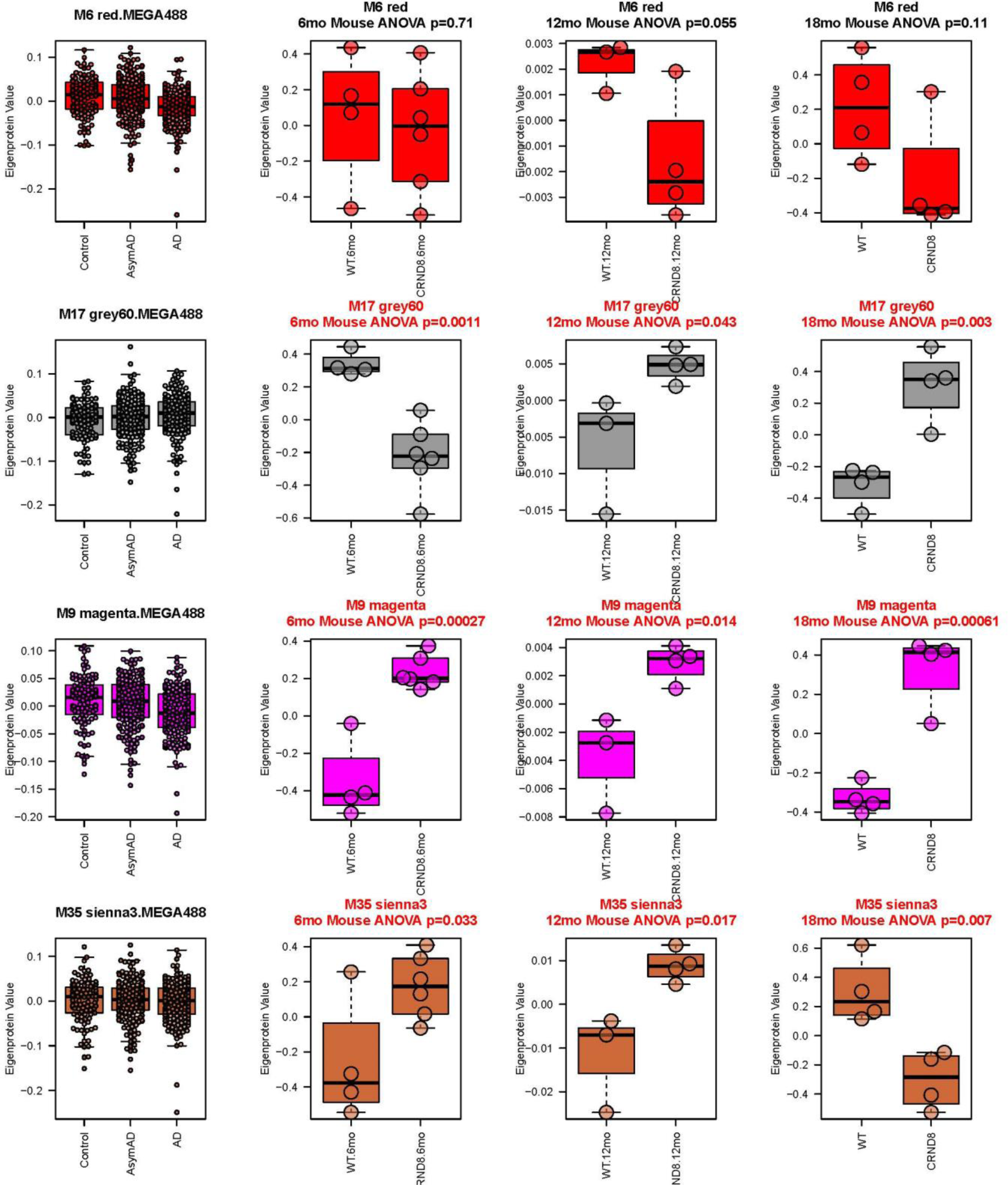
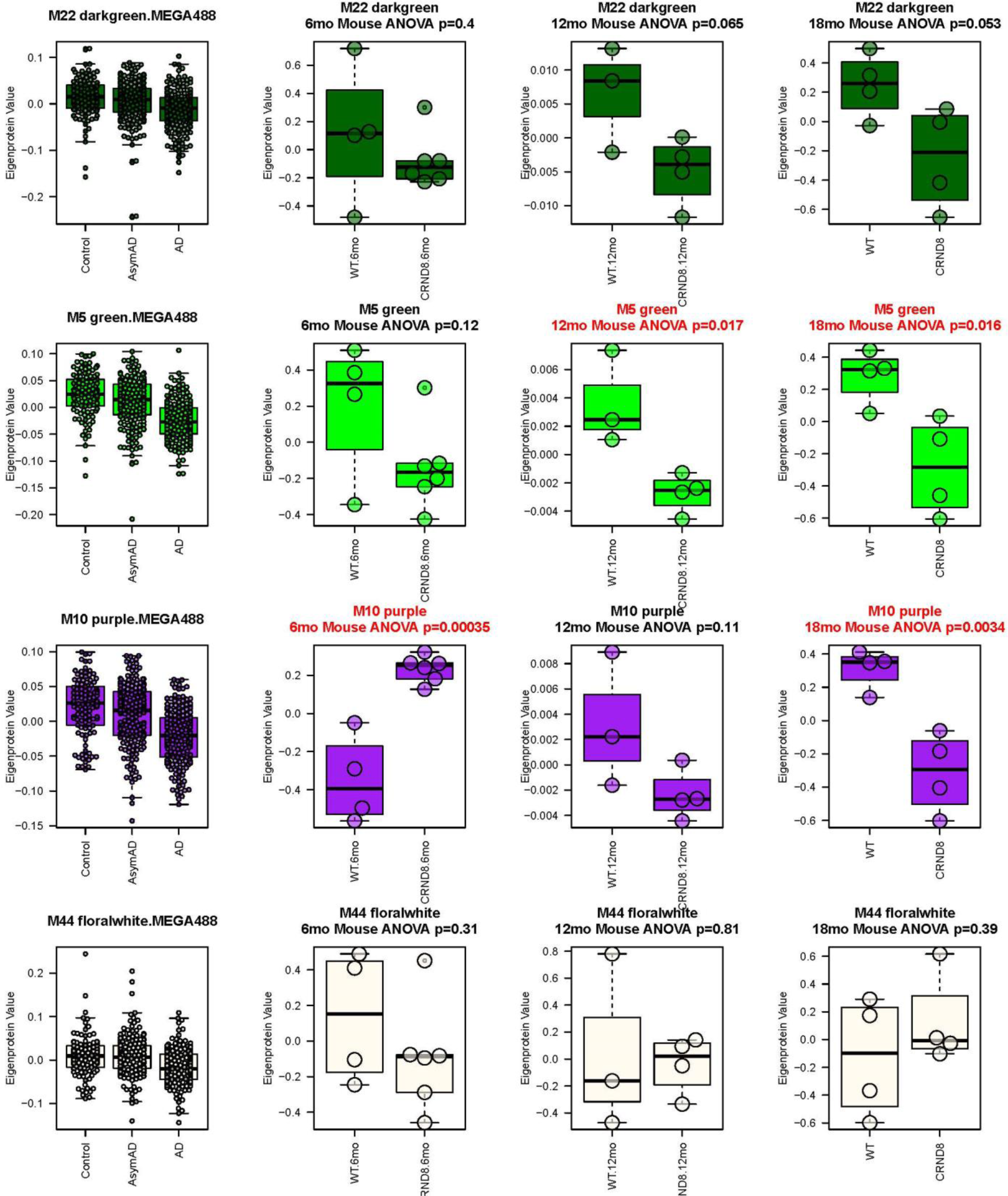
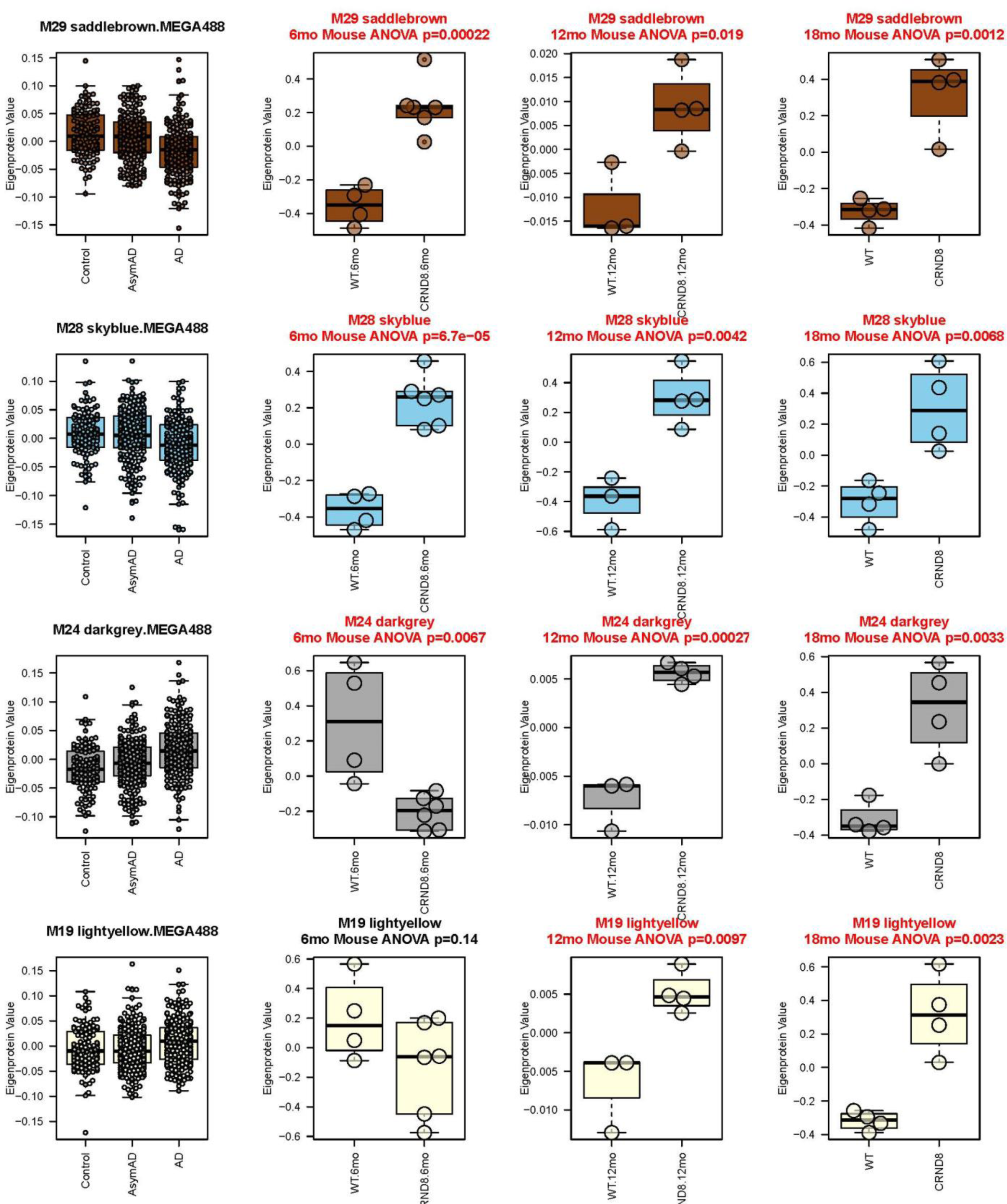
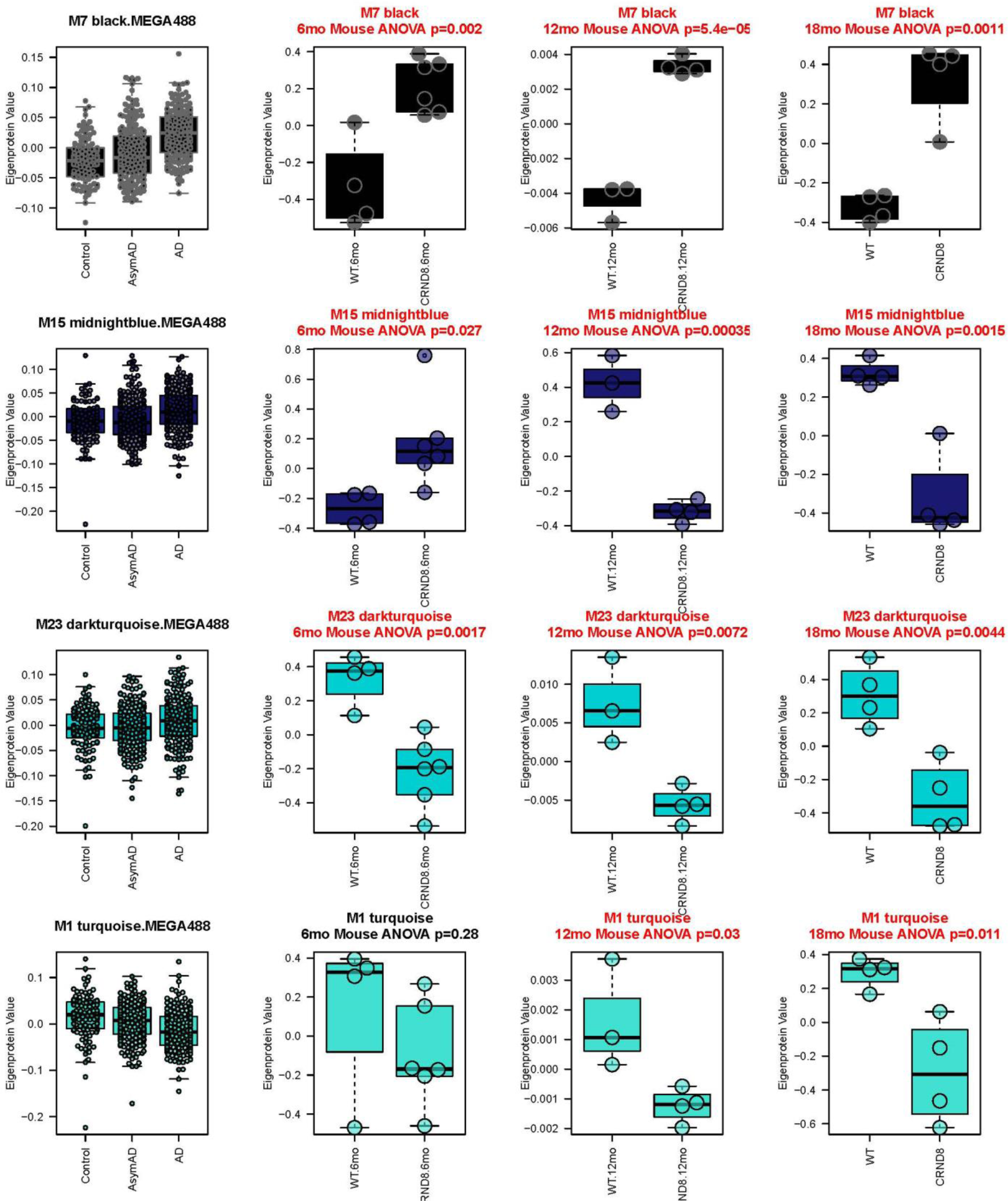
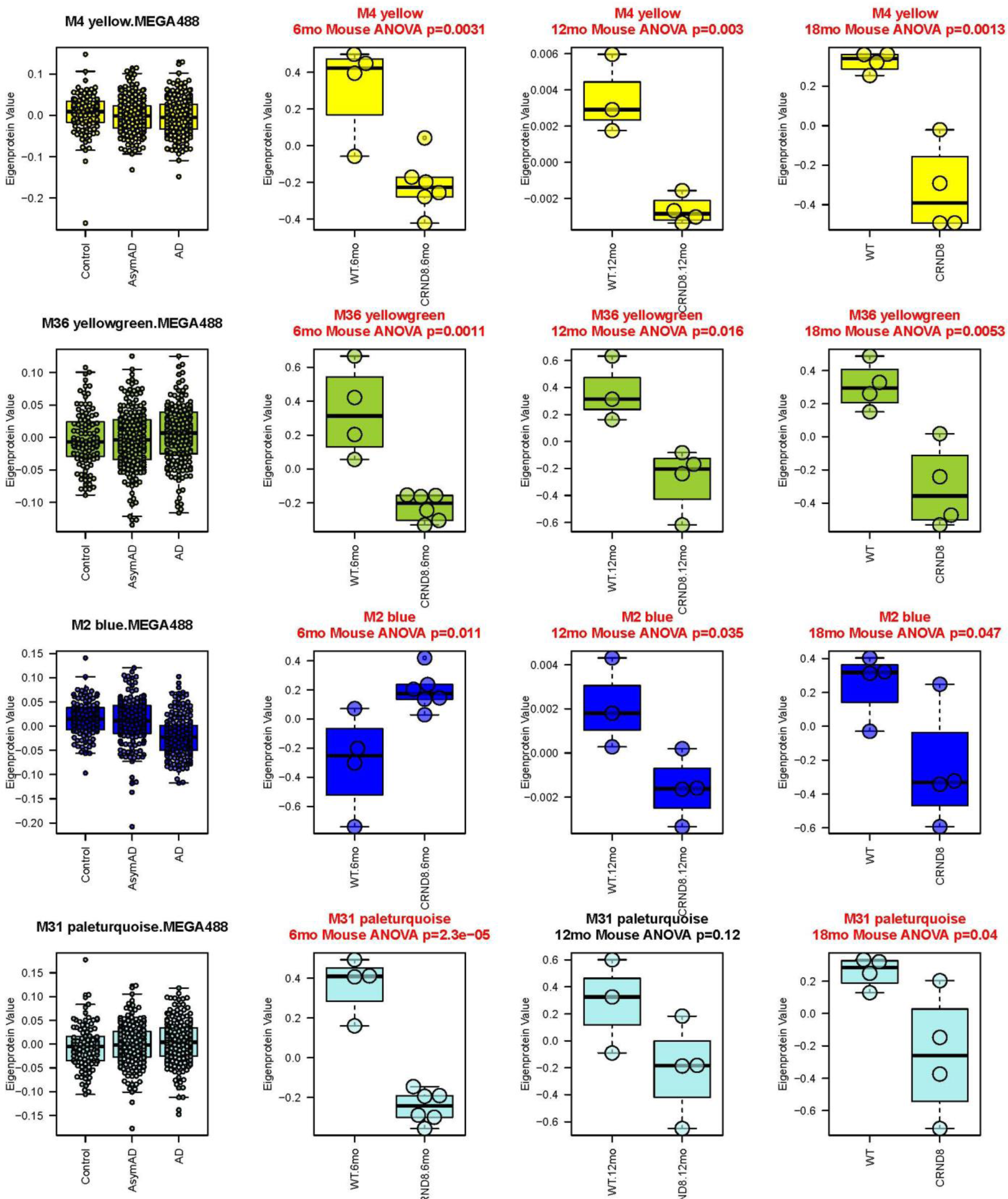
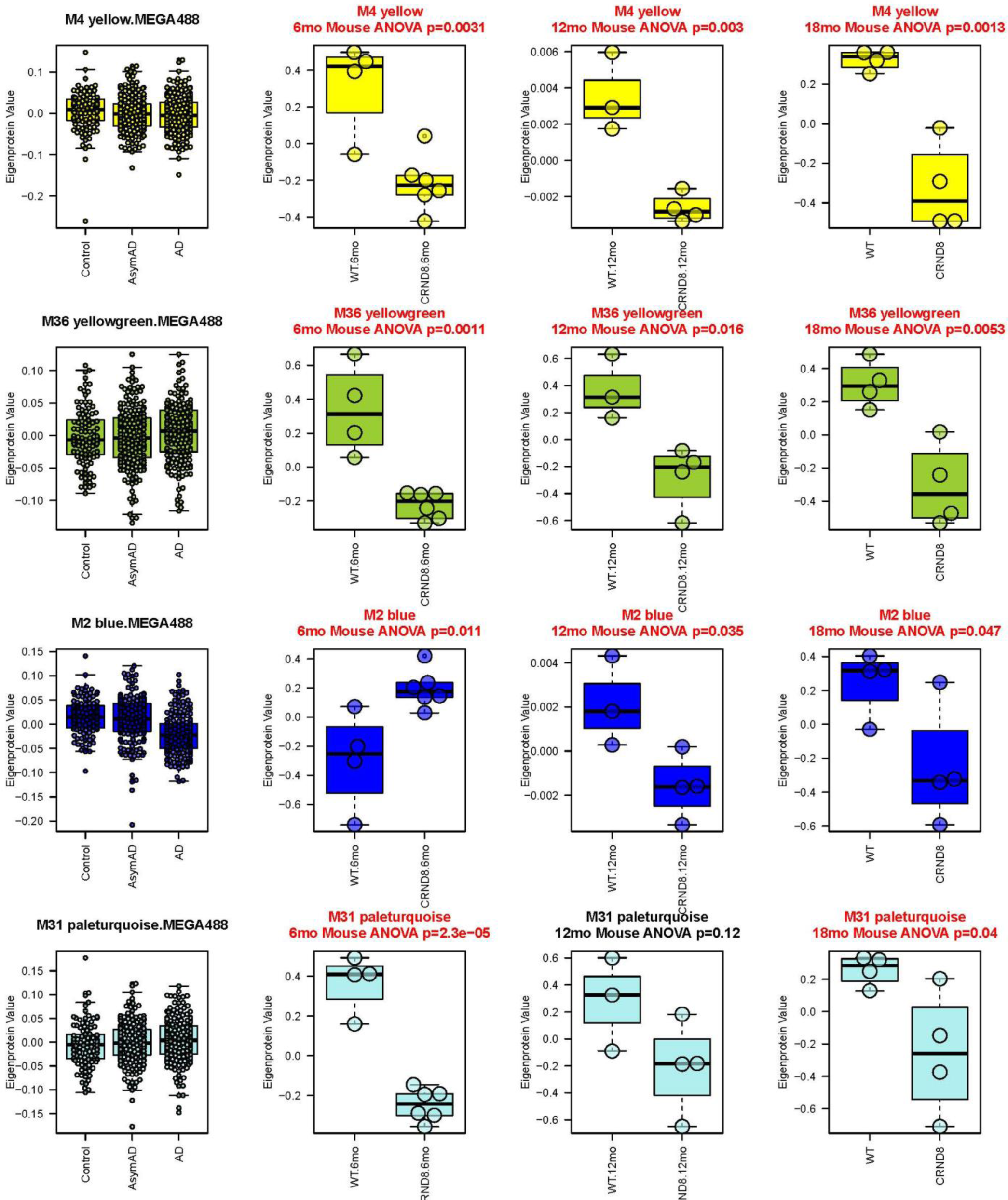
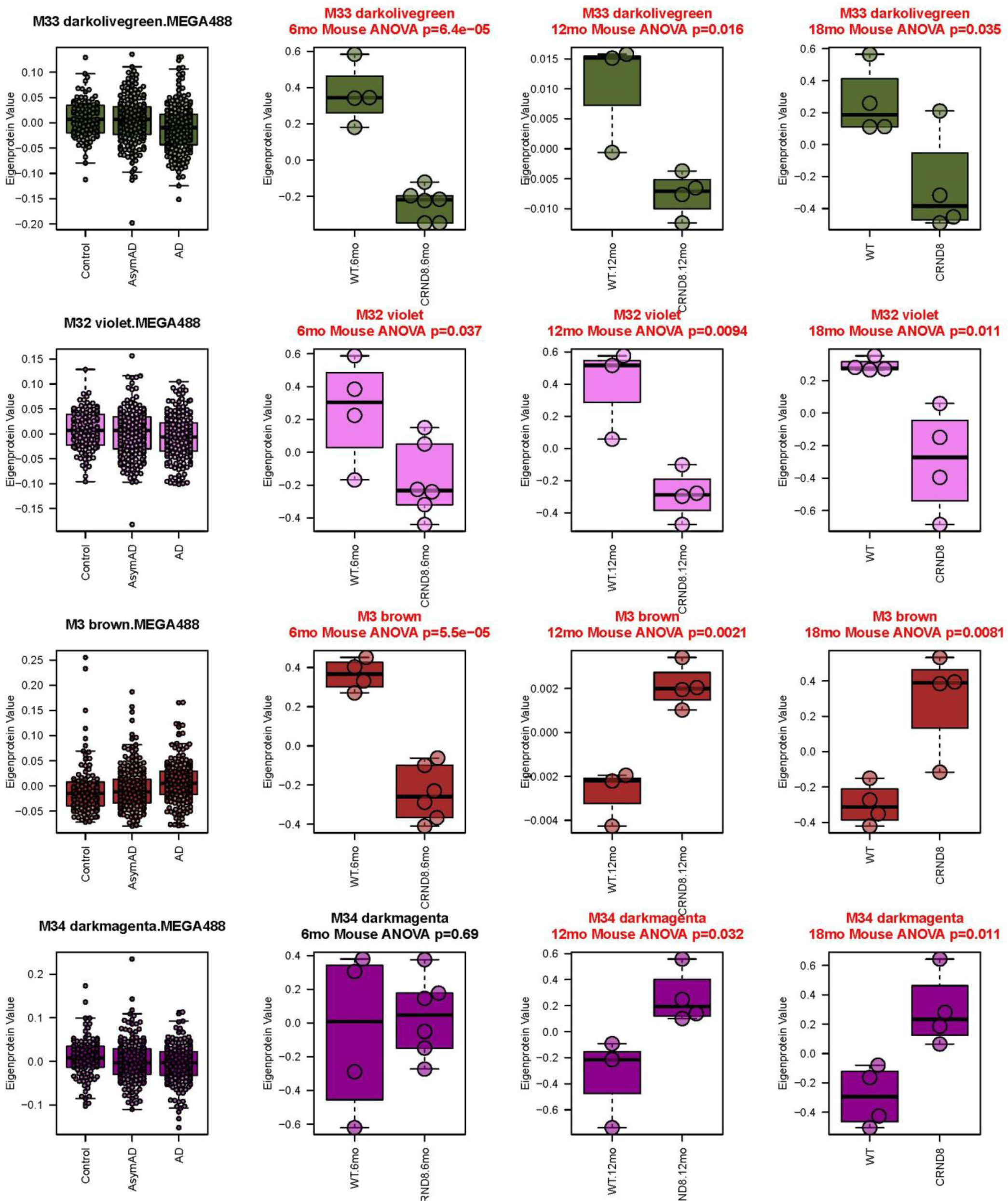
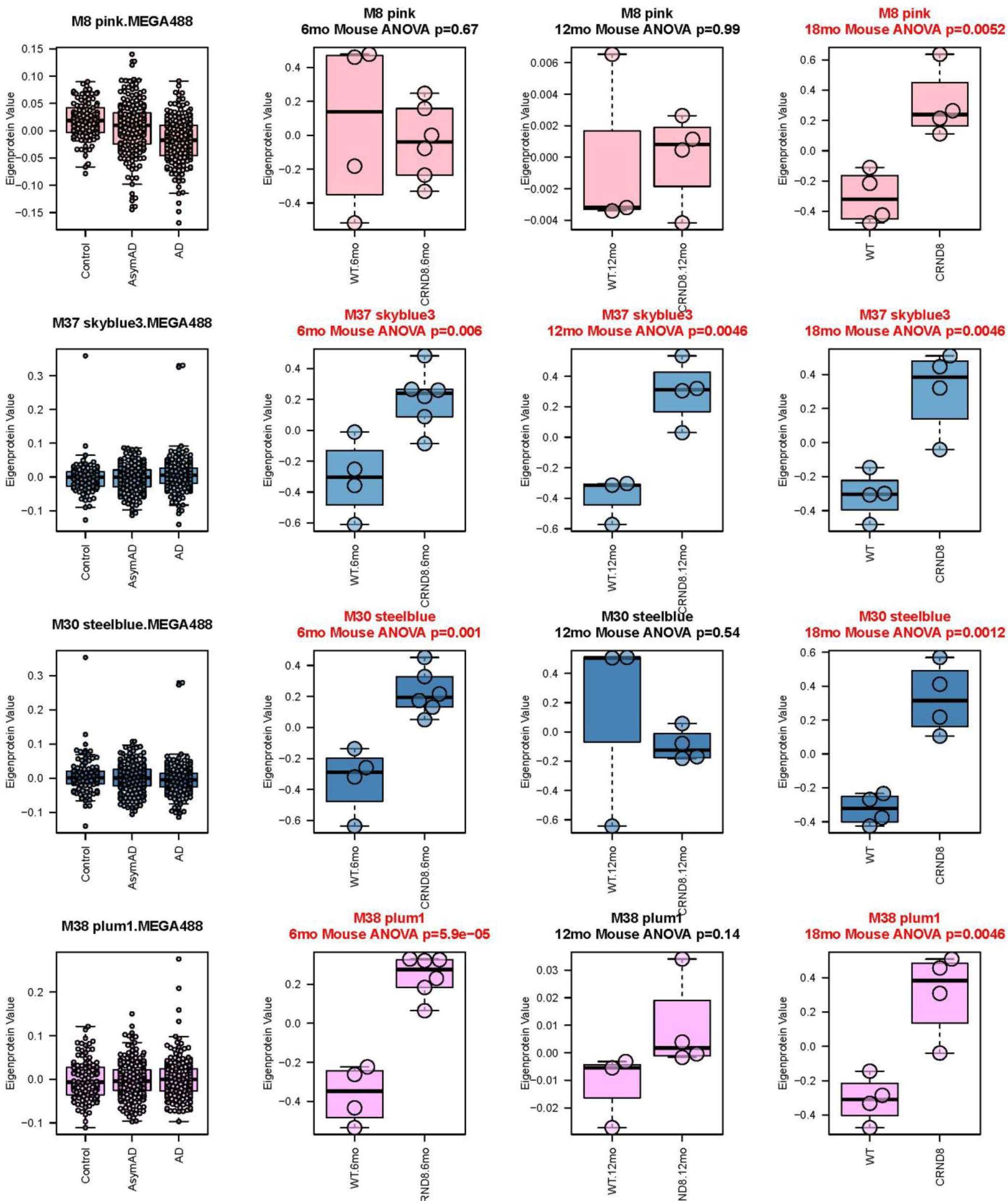
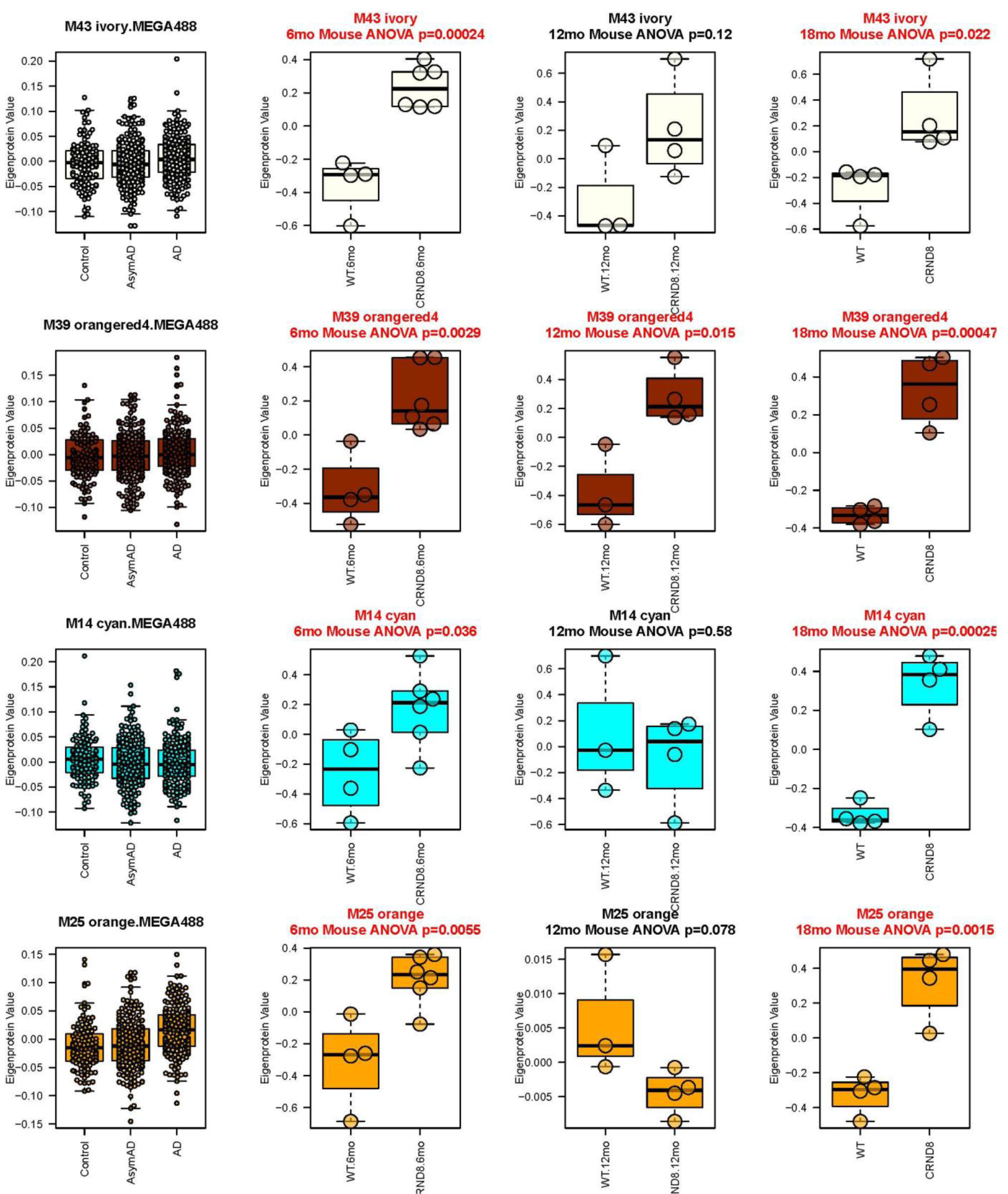
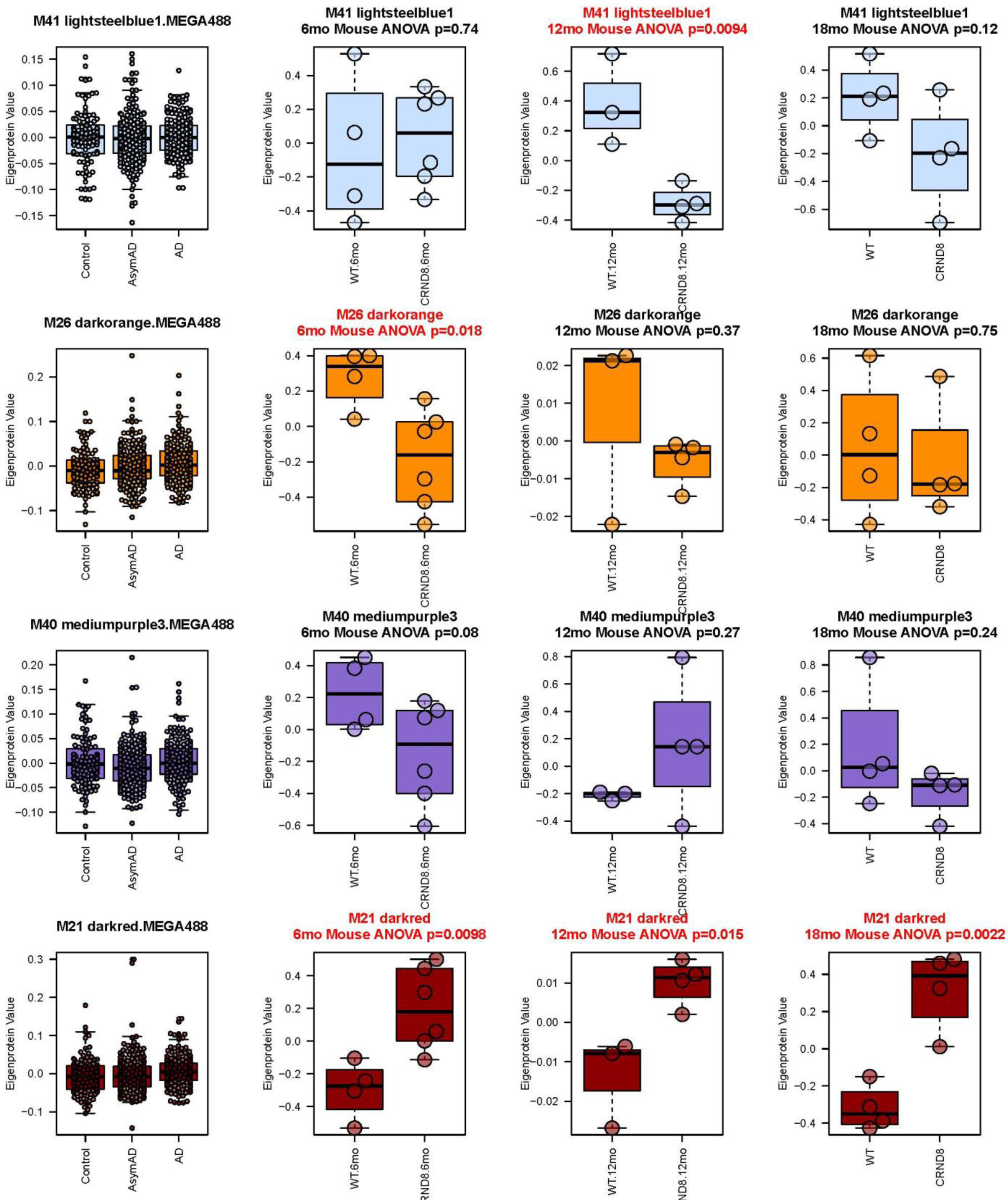
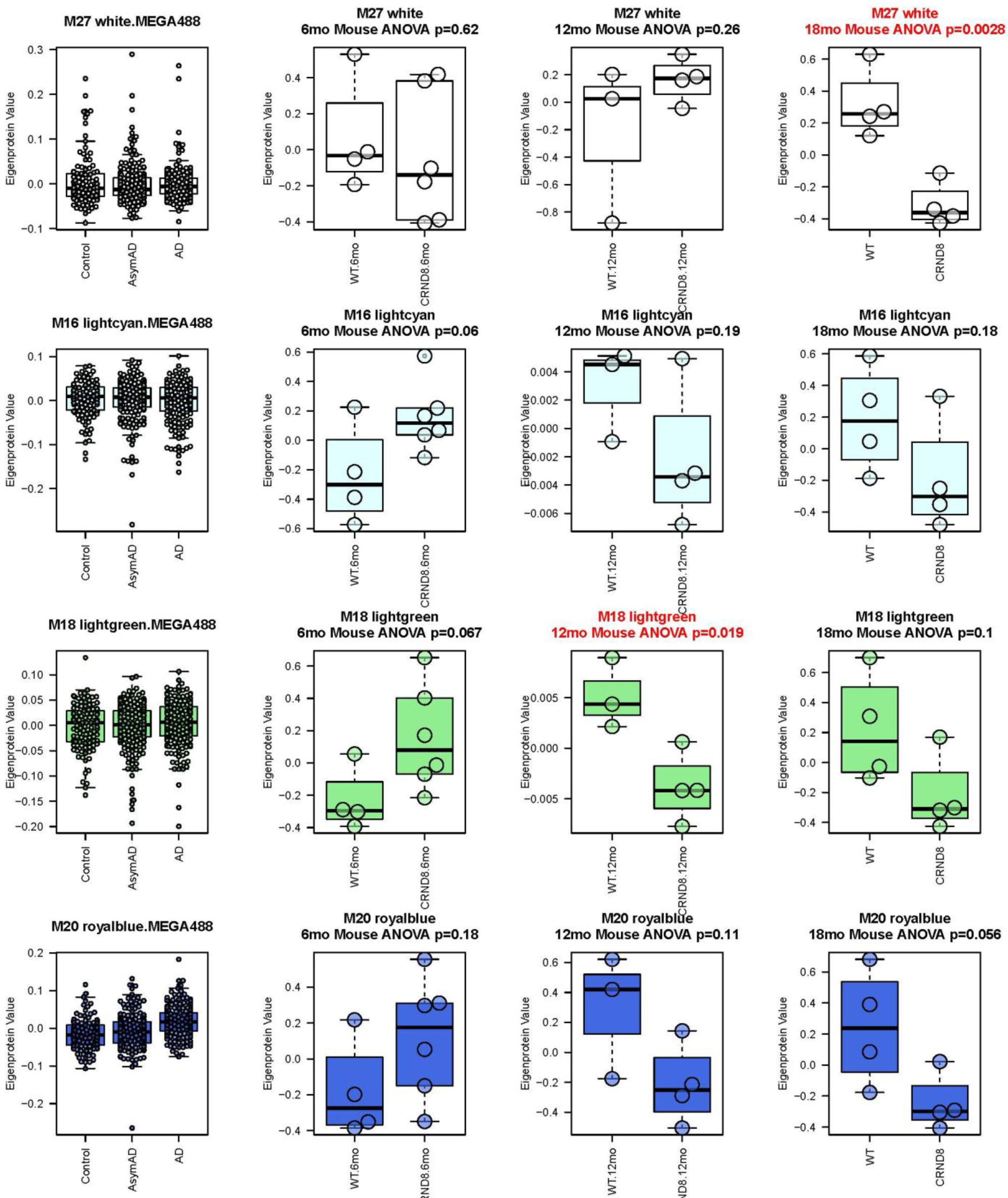
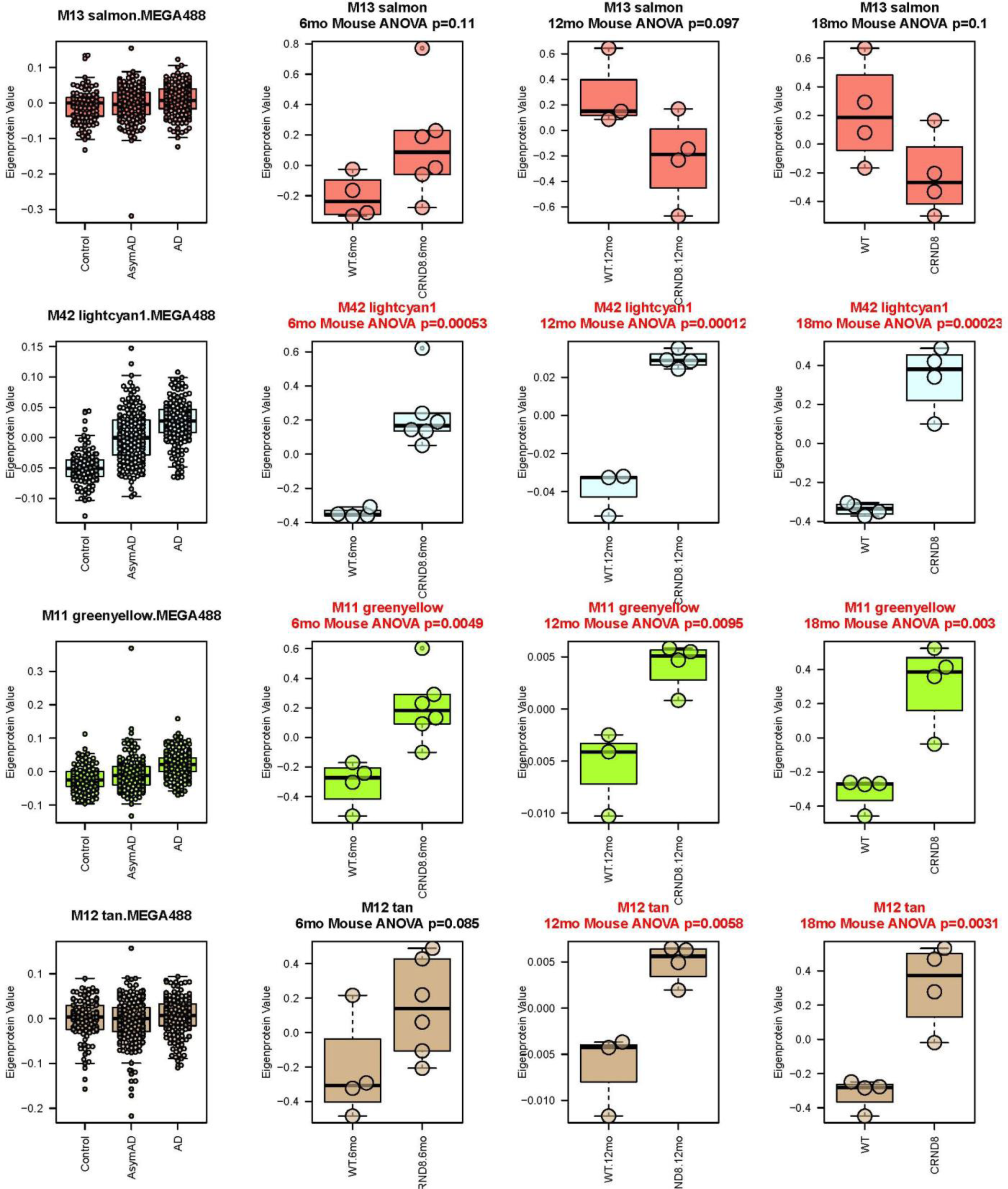
Synthetic eigenproteins (modules) show trends across the three cohorts of aged Tg and non-Tg mice for all 44 consensus brain modules. All 44 human modules were calculated as synthetic eigenproteins in the mouse proteomics and plotted as boxplots. Boxes represent the second to third interquartile range; whiskers represent the 5^th^ to 95^th^ percentiles; and the horizontal line within each box represents the median value.

**Figure S4.**
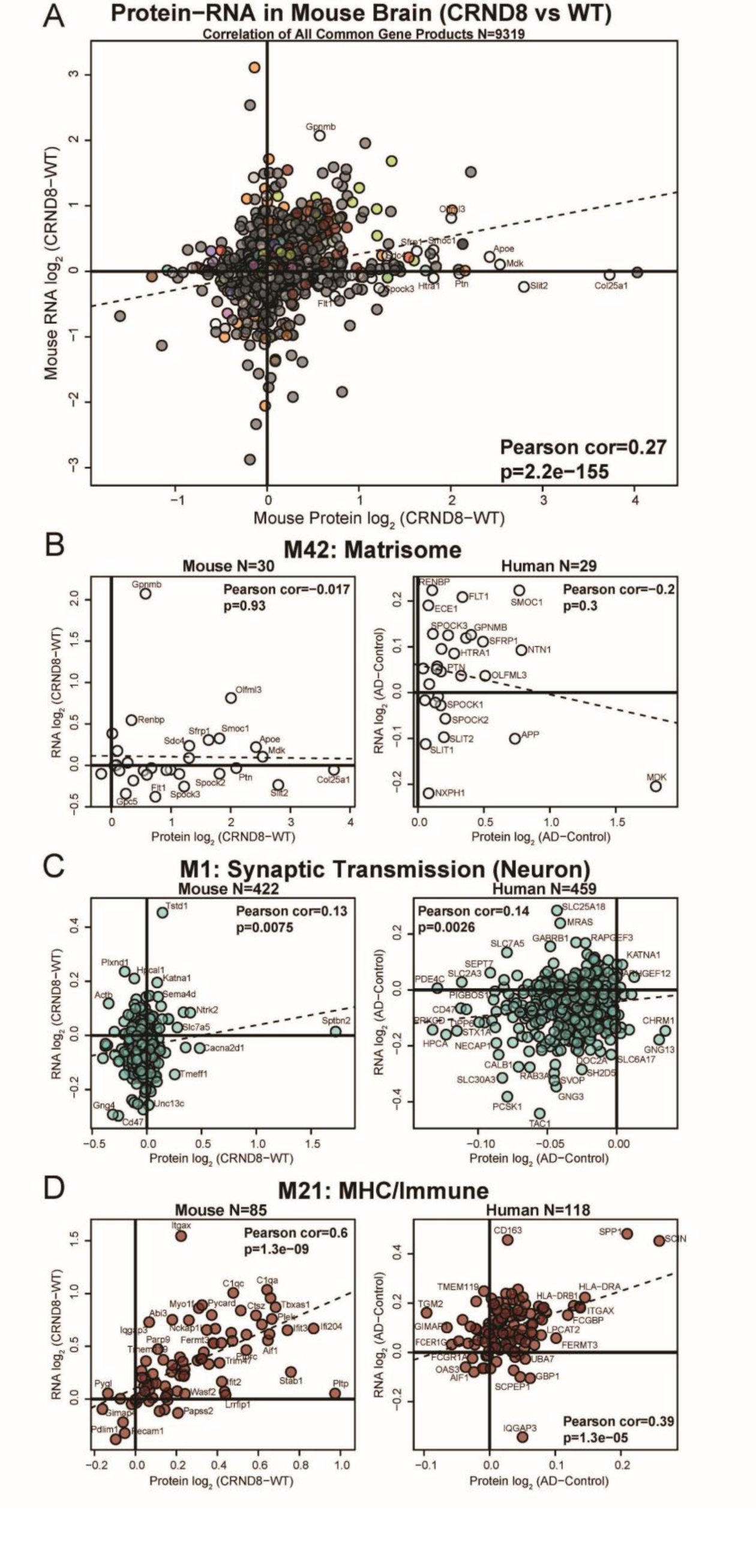
RNA-Protein AD or Tg Effect Size Correlation, in total and within modules. (A) The correlation of Tg effect size in mouse proteome (x) to transcriptome (y) shown with a least squares trendline over a scatterplot of the effect sizes for 9319 mouse gene transcripts and corresponding proteins (Table S8). When more than one splicing isoform protein was present, the one with maximum variance was retained. (B) M42 protein (x) to transcriptome (y) effect size scatterplot with least squares trendline, Pearson correlation (rho), and Student’s significance of correlation are shown for Tg vs non-Tg effect sizes (*left*), and in human AD vs. Control effect sizes (*right*). (C) M1 RNA-protein correspondence for Tg effect (*left*) and for the effect in human of AD vs. control (*right)*. (D) M21 RNA-protein correspondence for Tg effect (*left*) and for the human AD effect (*right*).

**Figure S5.**
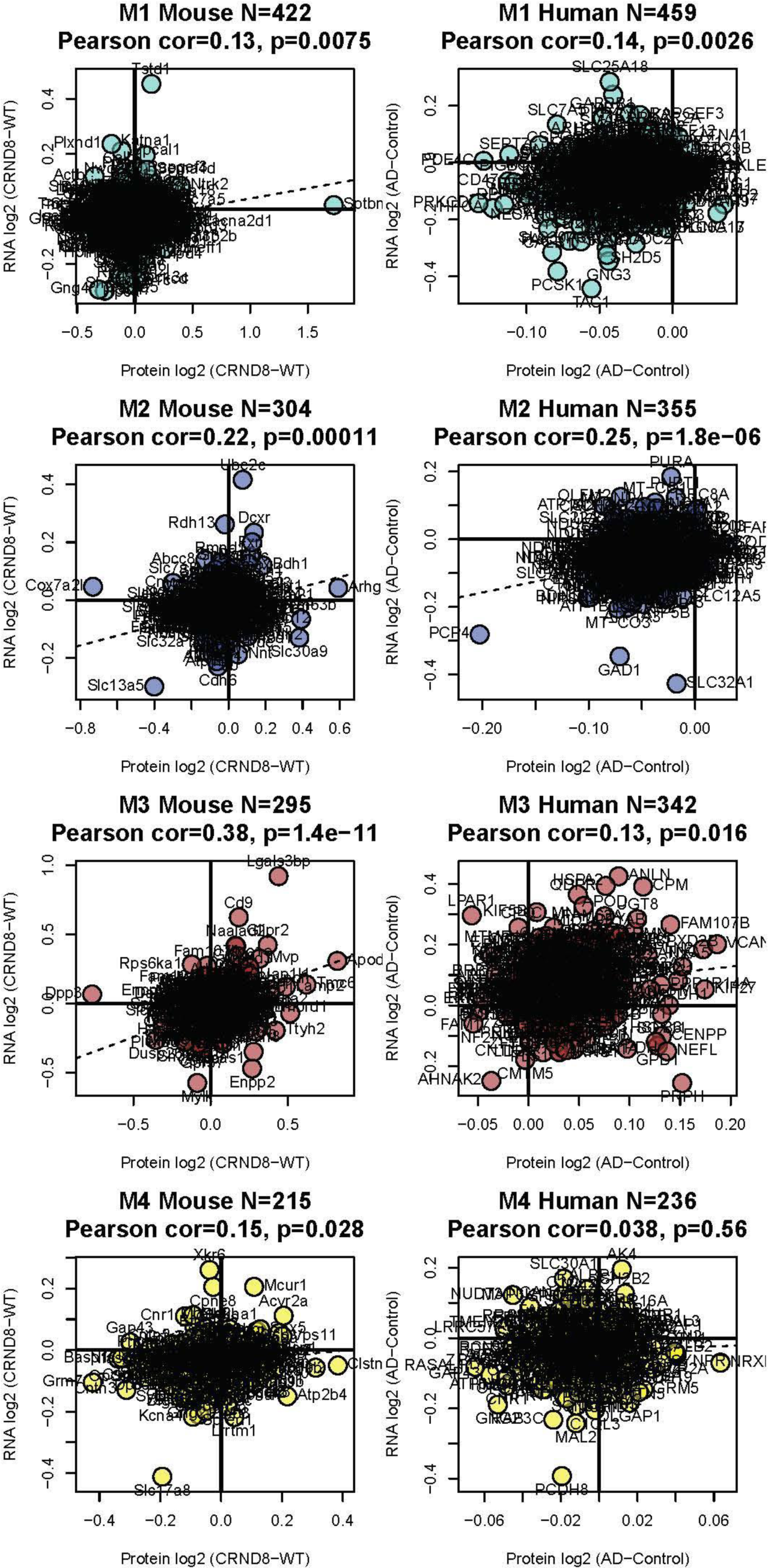
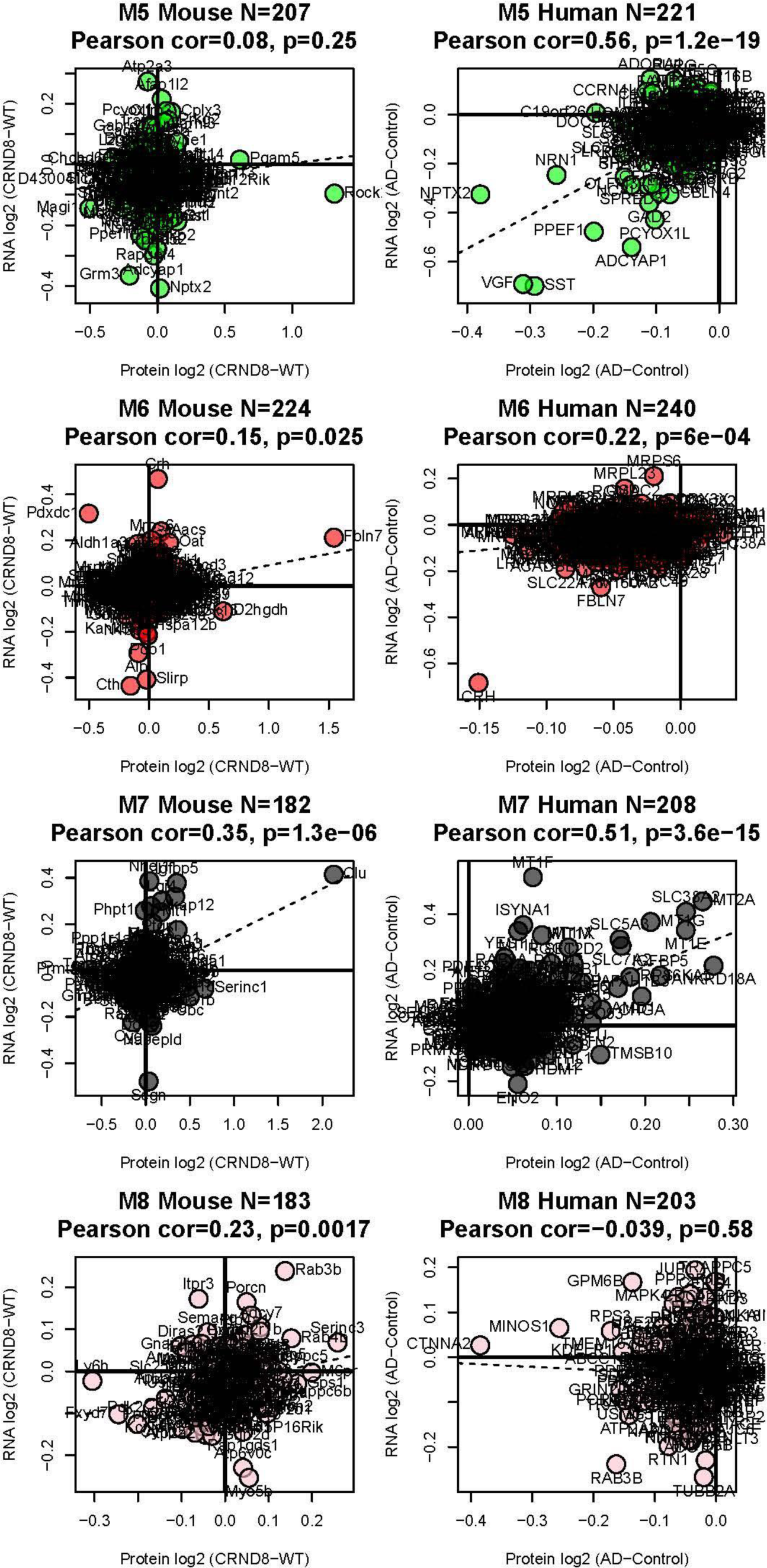
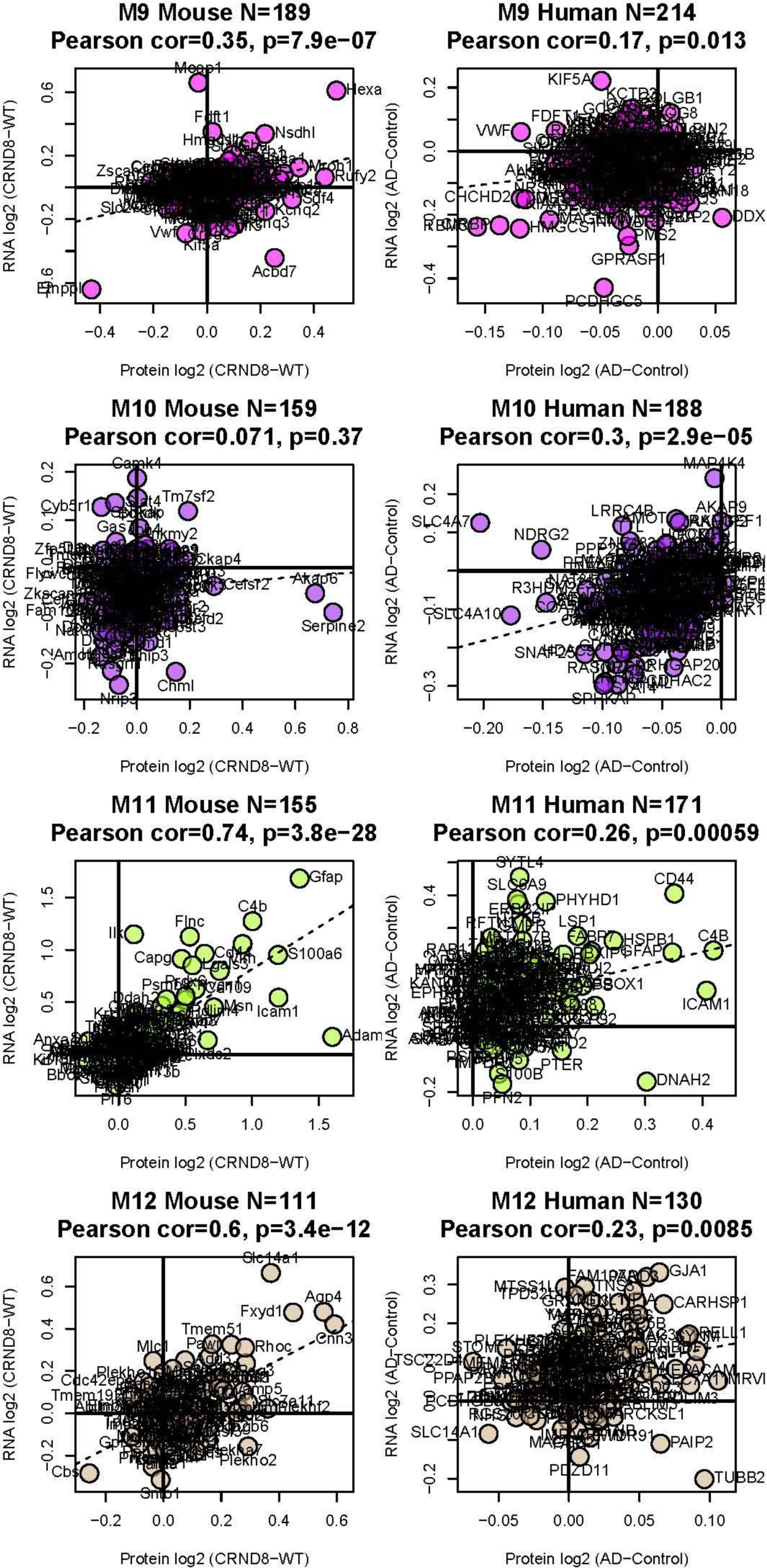
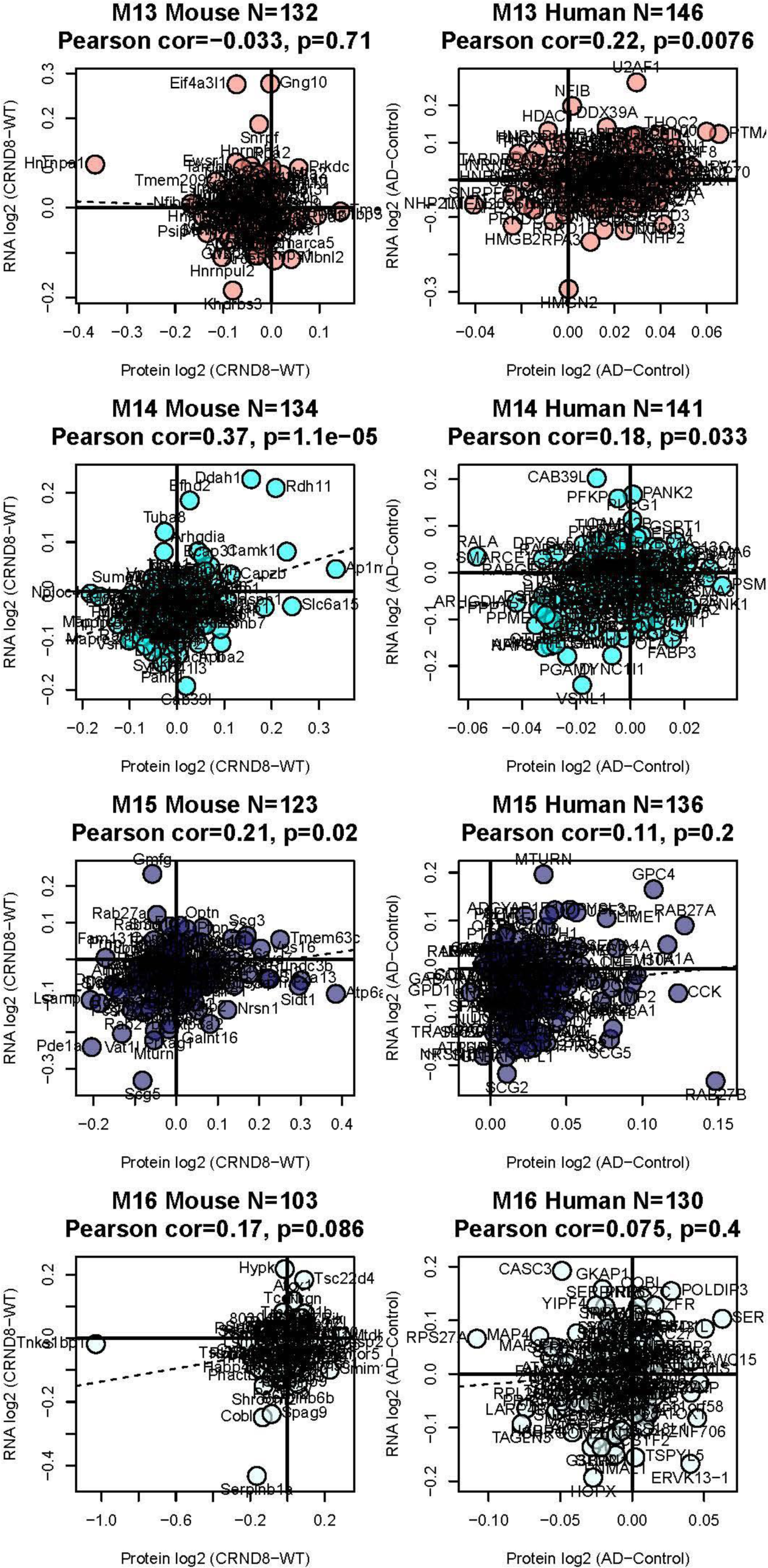
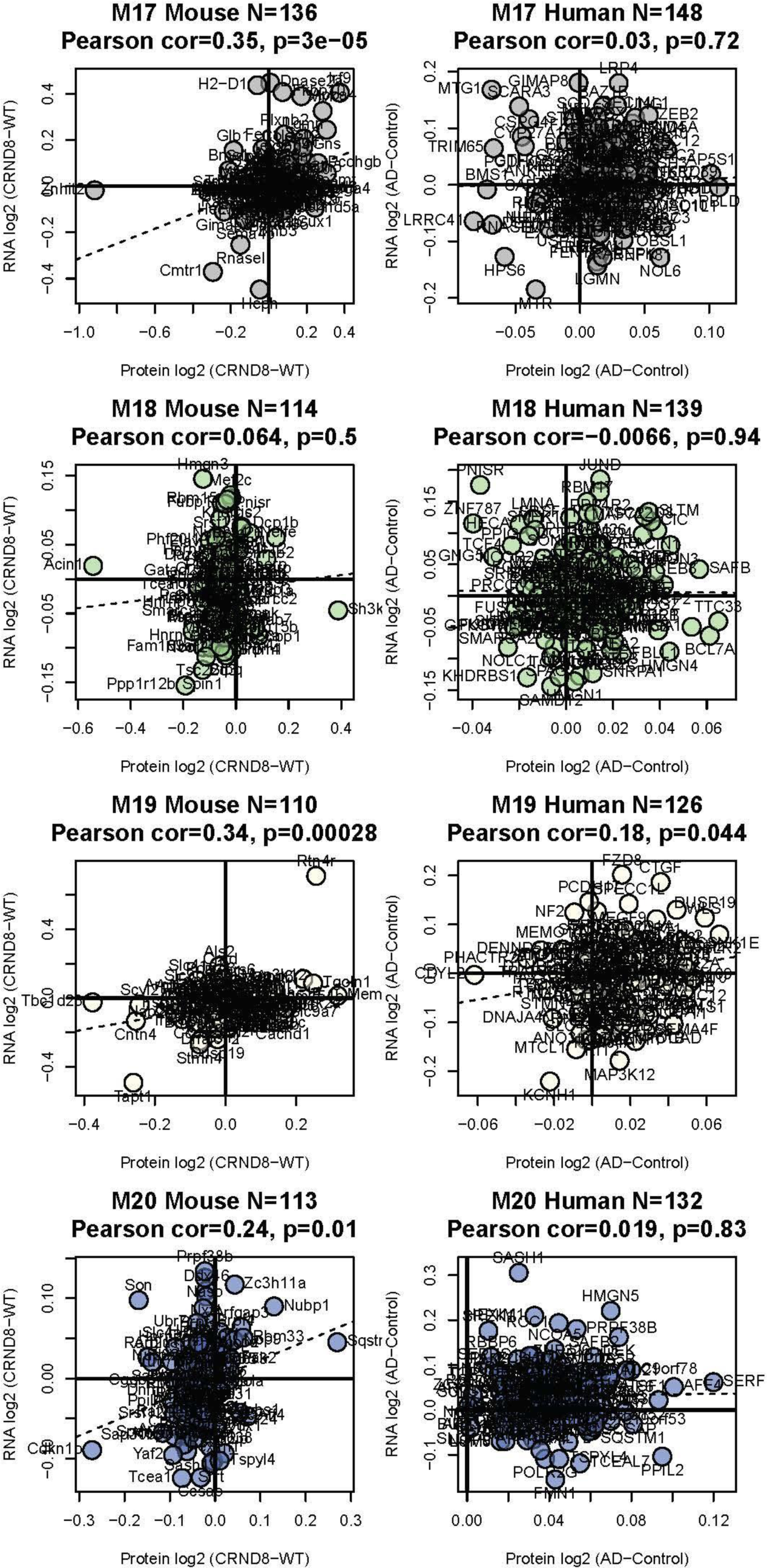
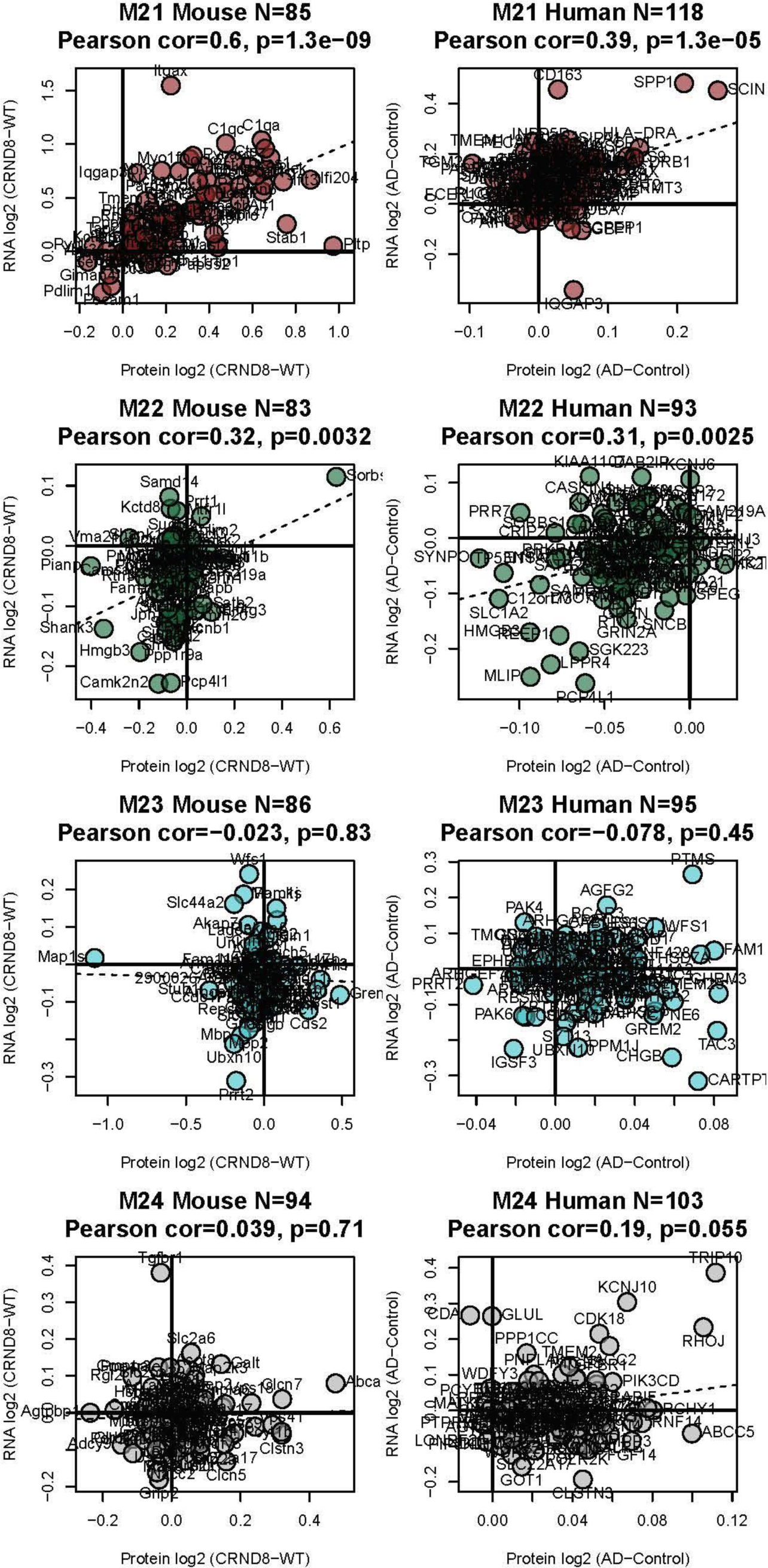
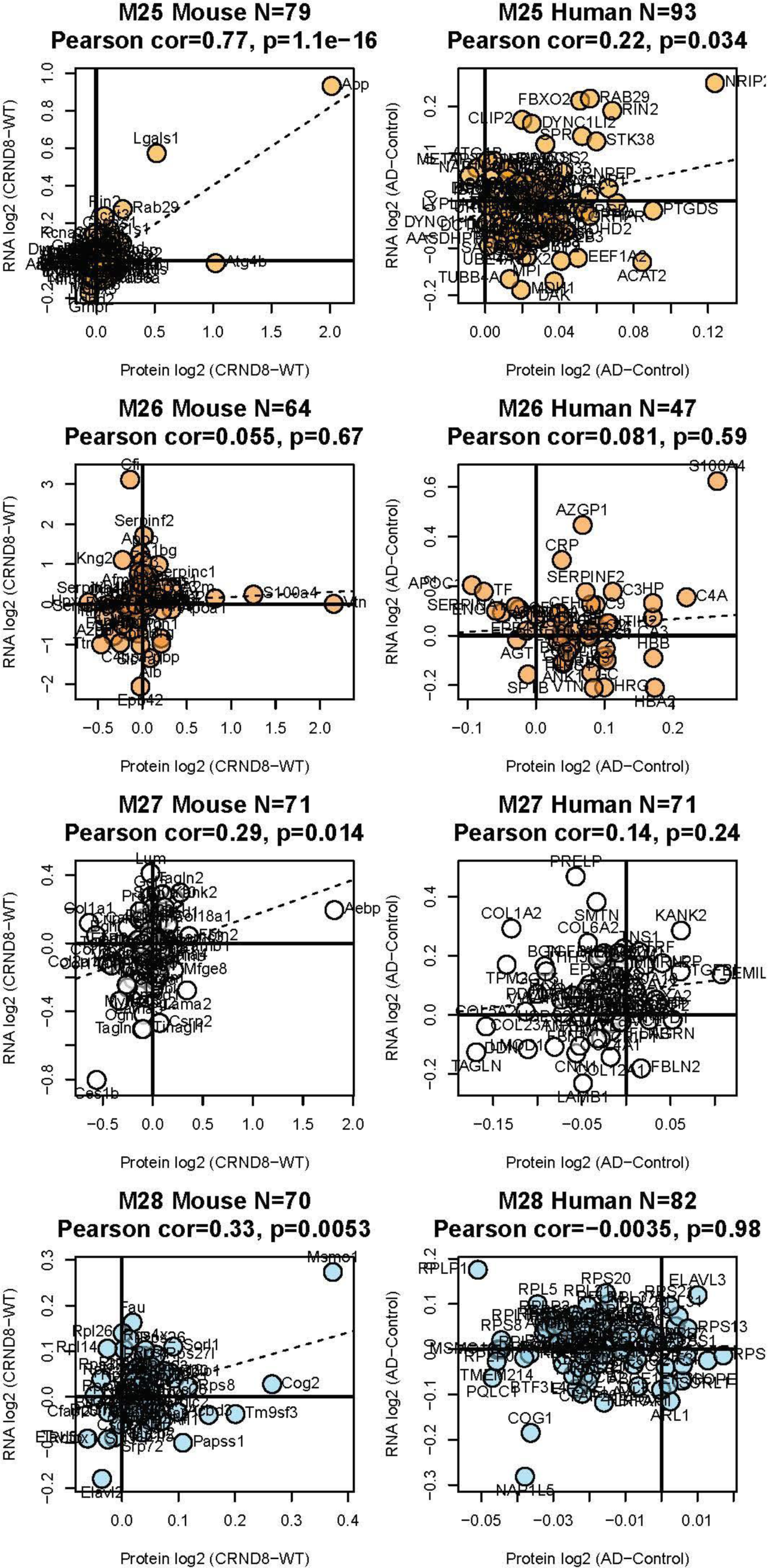
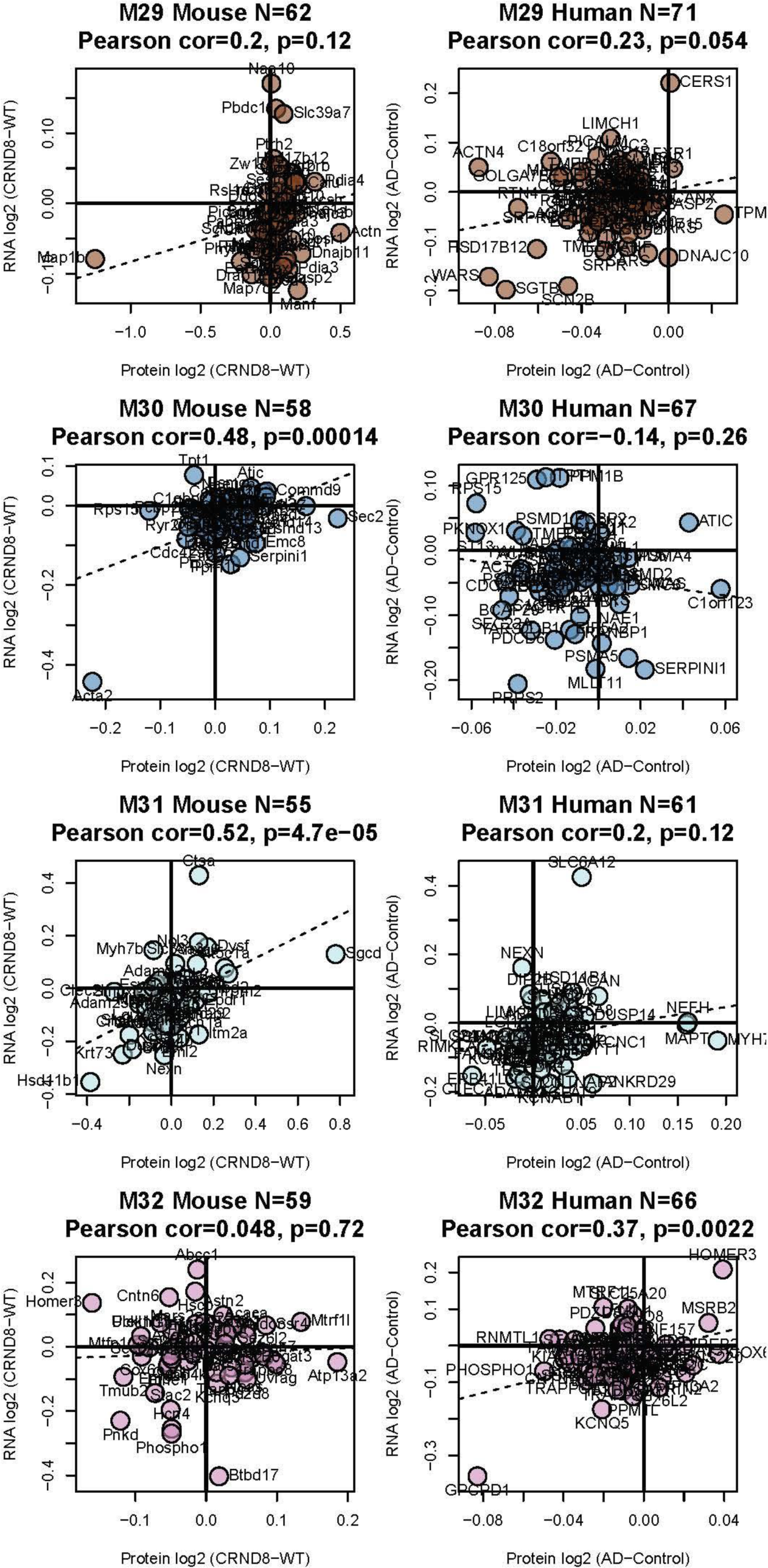
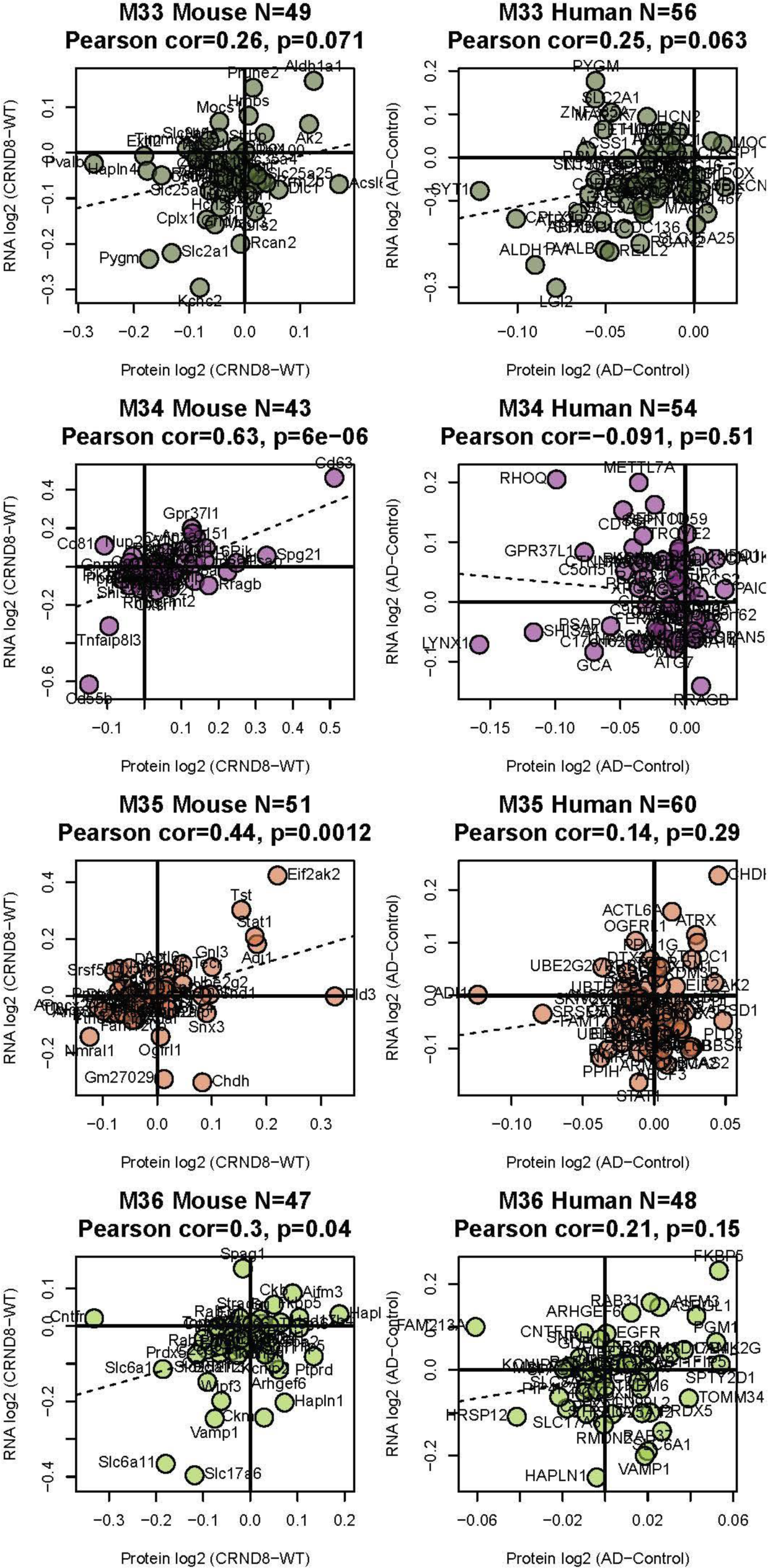
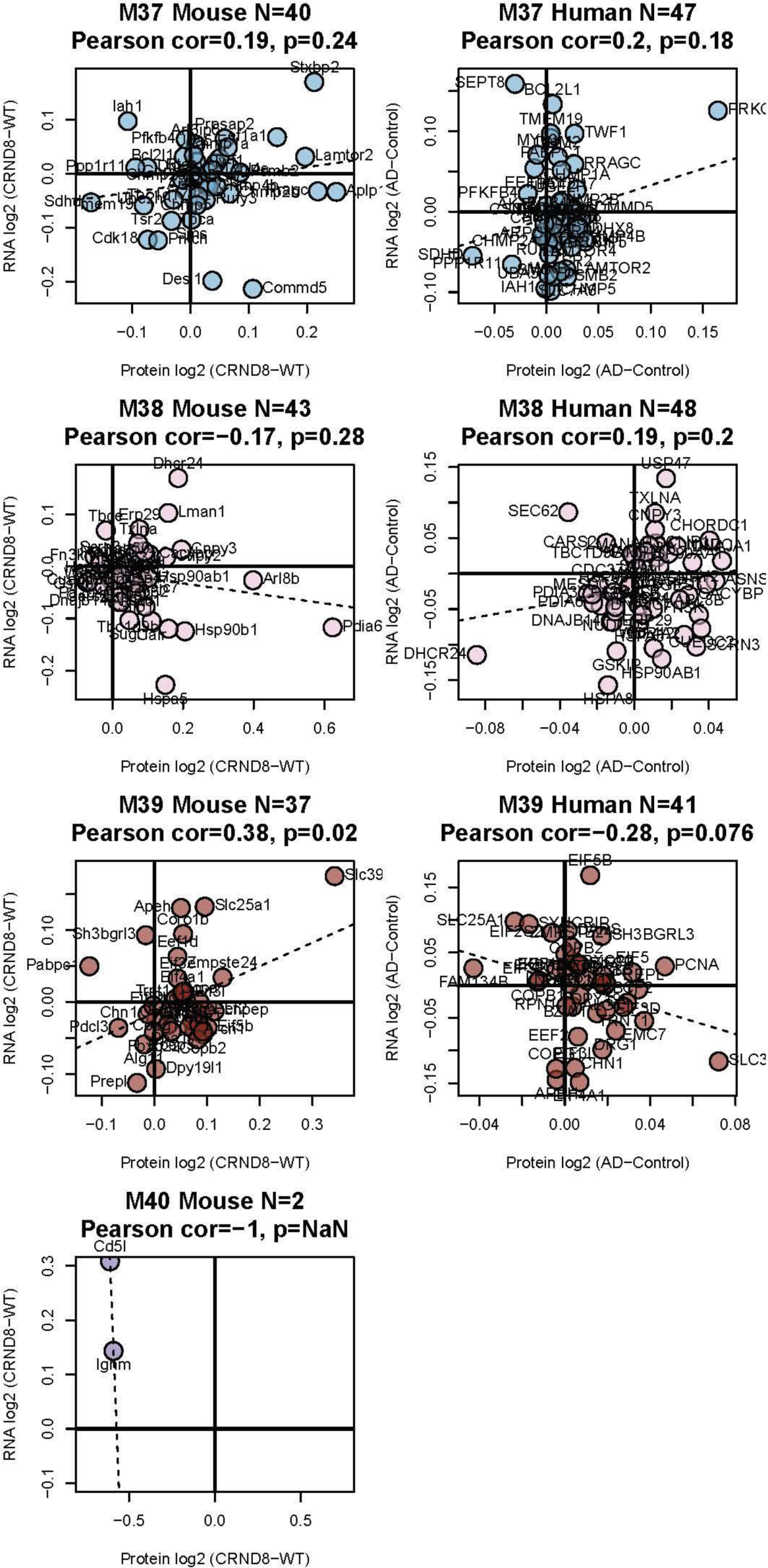
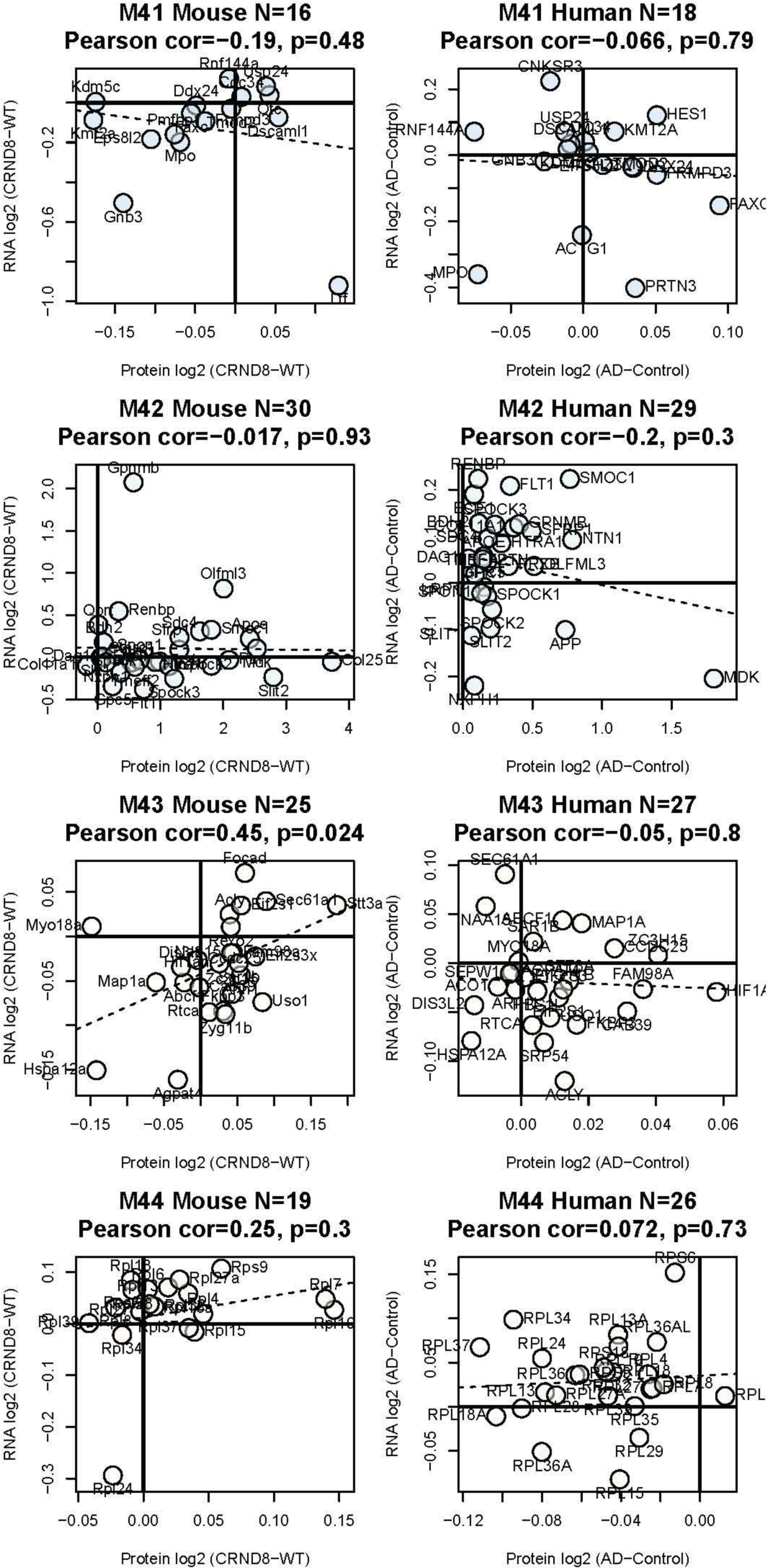
Protein-RNA effect size correlation varies in mouse and human for all 44 consensus brain modules. Gene product names label each point, with the module number (size rank), count of proteins mapped to mouse (*left*) from human (*right*), Pearson correlation rho and Student’s significance of correlation given in each plot title.

**Figure S6.**
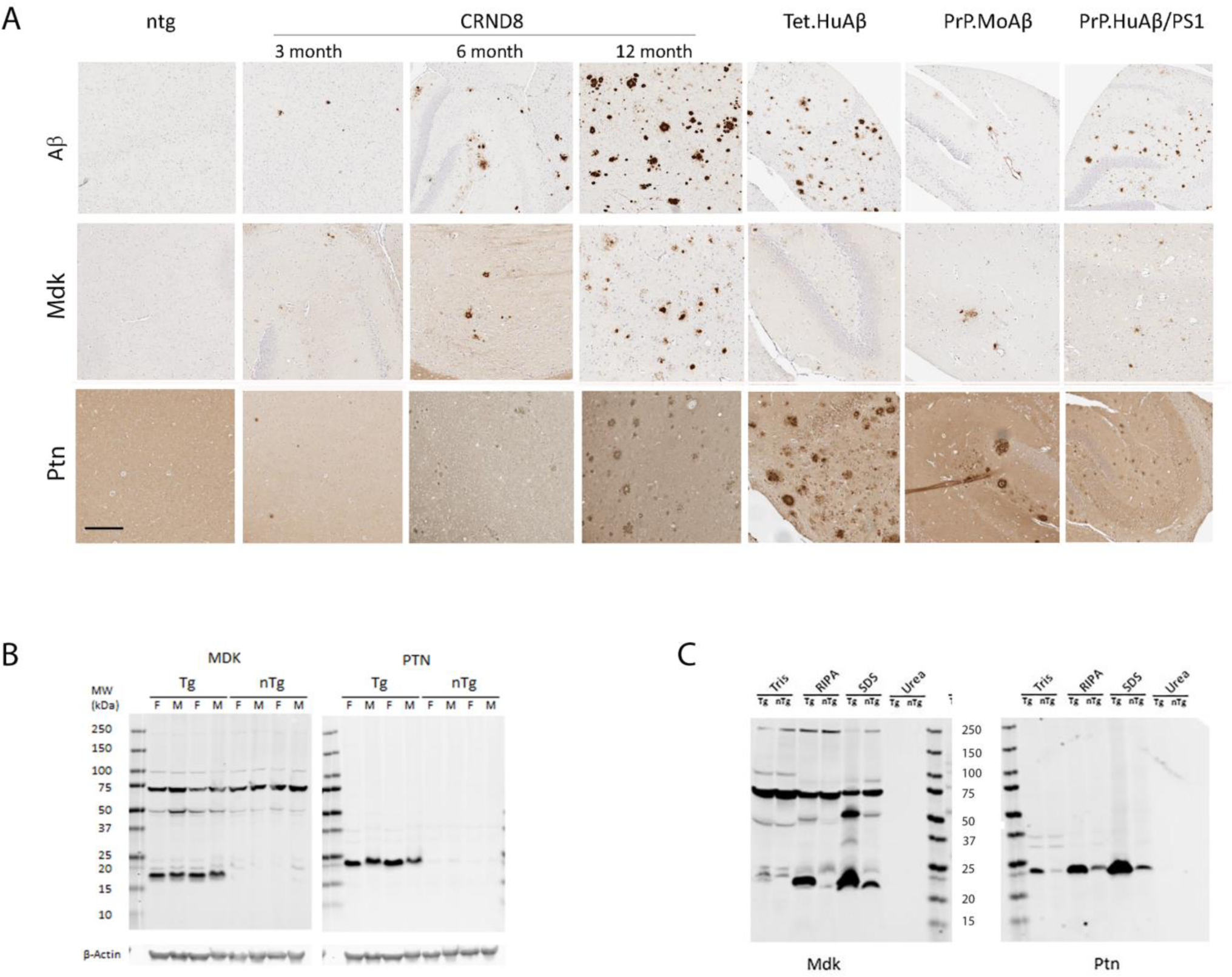
Mdk and Ptn colocalization with amyloid is age dependent, occurs in various AD transgenic mouse models, and accumulation does not alter overall solubility. (A) Paraffin slides containing brain tissue from 12M non-Tg mice; 3, 6 and 12M CRND8 mice; 12M Tet.HuAPP, PrP.MoAβ and PrPHuAβ/PS1 mice were stained with mouse anti-pan-Aβ, sheep anti-Mdk or rabbit anti-Ptn antibodies. scale 50 mm. (B). Western blots detecting Mdk and Ptn in the urea extracts of 15M CRND8 mice and non-Tg control brains as shown by SDS-PAGE stained with anti-Mdk or anti-Ptn antibodies in total Urea extracts (first two lanes for each are male and second two lanes are females. Blots are representatives of three independent experiments. (C) Sequential extractions with TBS, RIPA, 2% SDS and 8M Urea, show that neither Mdk nor Ptn become detergent insoluble. Blots are representatives of three independent experiments.

**Figure S7.**
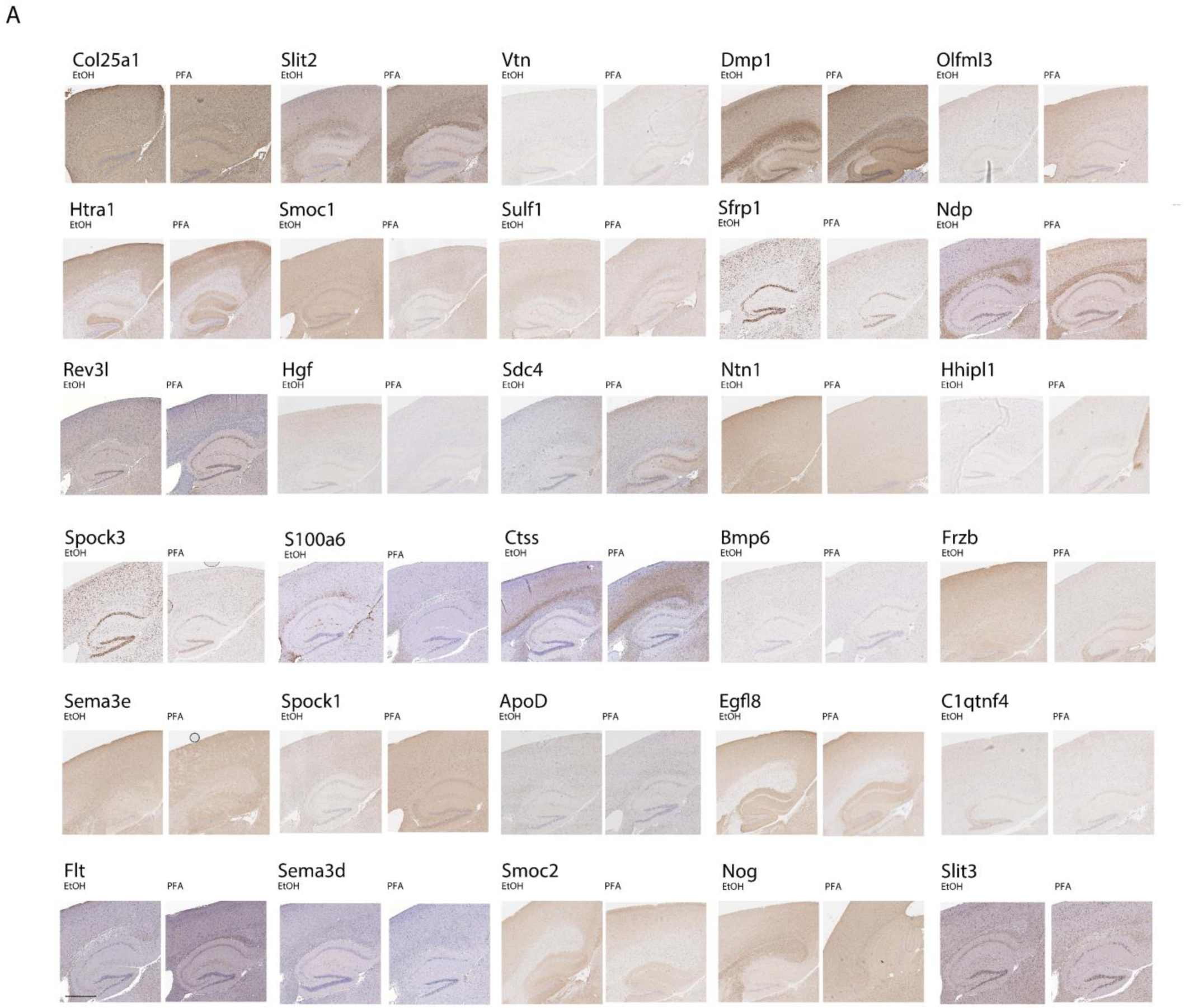
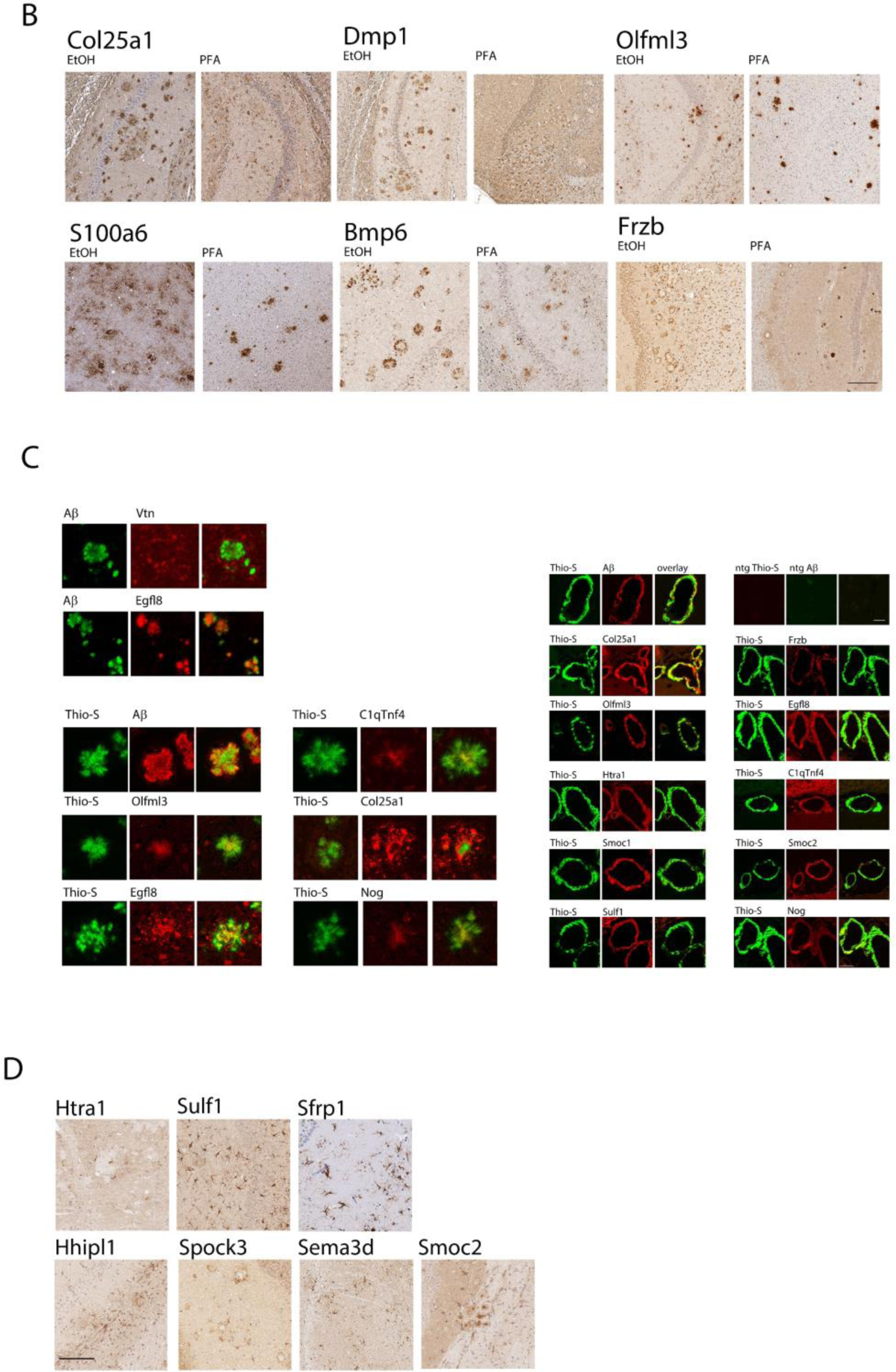
Non-Tg staining, fixation effects, co-localization, and astrocyte staining. (A) DEP staining of 15-18M old non-Tg controls. Paraffin slides containing brain tissue from 15-18M non-Tg fixed in 4% paraformaldehyde (one hemibrain) or 10% ethanol (second hemibrain), were stained with anti-DEP antibodies. Low magnification (scale 500 μm) images were taken from cortex and hippocampus. (B) Fixation impacts localization, but impacts are dependent on the antiserum used. Paraffin slides containing brain tissue from 15M CRND8 mice, fixed in 4% paraformaldehyde (one hemibrain) or 10% ethanol (second hemibrain), were stained with the indicated anti-DEP antibodies. Scale 200 μm. (C, D) Colocalization of DEP in plaques. Paraffin slides containing brain tissue from 15M TgCRND8 mice, were stained with either mouse anti-pan-Aβ antibody or Thio-S and rabbit anti-DEP. Secondary fluorescent anti-mouse (green) and rabbit (red) antibodies were used. Colocalization was visualized by overlap of green and red colors to create yellow in plaques (C) or CAA (D) Scale 50 μm. (E) Select DEPs accumulate in astrocytes surrounding amyloid plaques. Paraffin slides containing brain tissue from 15-18M CRND8 mice were stained with anti-DEP antibodies. Scale 200 μm.

**Figure S8.**
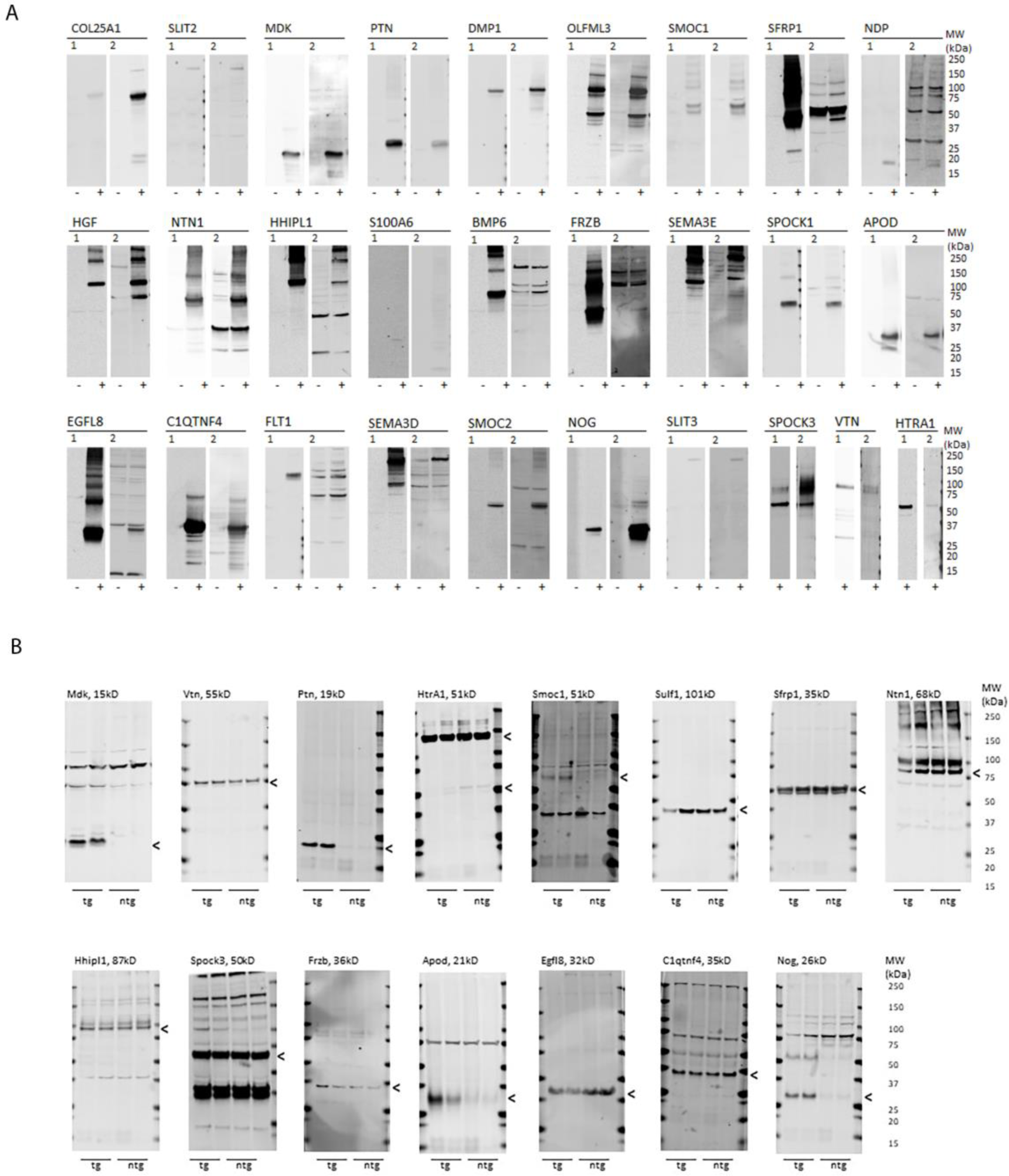
Antibody validation studies and additional DEPS validated by Western blotting. (A) Antibody validation. 293T HEK cells were transfected with plasmids containing FLAG-tagged cDNAs encoding M42+ DEP and gene expression in the media (SPOCK3, VTN, HTRA1) and lysates (all other genes) was detected on SDS-PAGE with 1 (anti-Flag) and 2 (anti-DEP sera). Non-transfected cells served as control. Representative blots of at least three independent experiments are shown. Sources of the antibodies are listed in Table S10, S11. (B) M42+ DEPs are detected in total UREA extracts from 15M CRND8 mice. Brain homogenates from Tg and age matched non-Tg mice were subjected to SDS-PAGE and stained with anti-DEP antibodies. (n=2; one male and one female).

**Figure S9.**
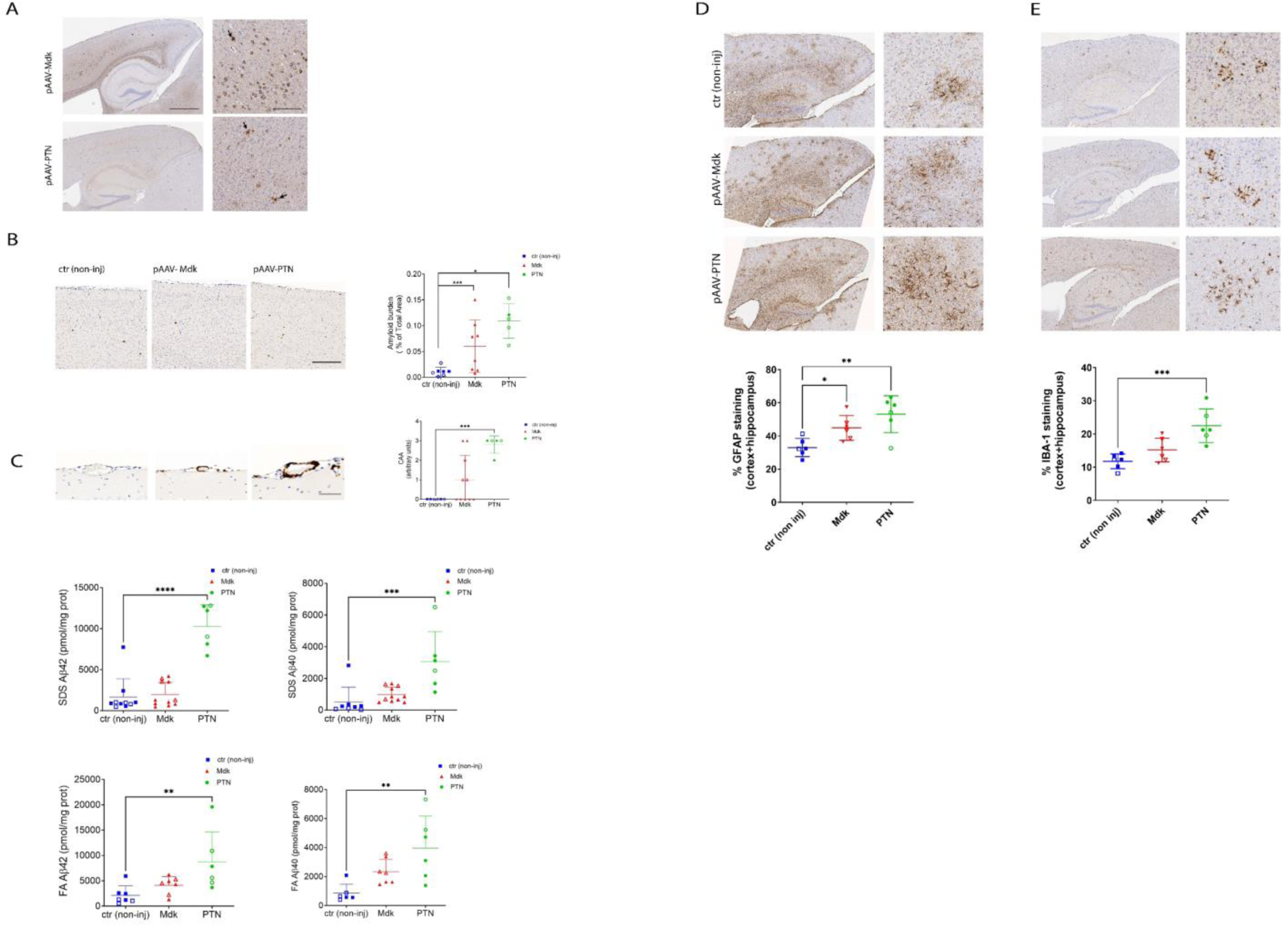
Midkine and Pleiotrophin overexpression increase formation of Aβ amyloid deposition and CAA in 3M CRND8 mice and 6M mouse data on gliosis. Mice were intracerebrally injected with rAAV2/8-Mdk, rAAV2/8-PTN or not-injected (control) at P0 and aged 3 months. (A, B) Representative images of cortex and hippocampus (A) or CAA (B) stained with biotinylated anti-Aβ mAb 33.1.1 (anti-Aβ 1-16). Scale bar: 500 µm, 50 µm (inset). Quantification data of the entire brain plaque count in three non-consecutive sections represented by a scatter dot plot of male (closed circle/square) and female (open circle/square) ± standard error of the mean. n=5-15. Plaques were quantified with one-way ANOVA test (*, p<0.05; ***, p<0.001). (C) RIPA, 2% SDS and 70% formic acid (FA) extracted Aβ40 and Aβ42 levels were detected by ELISA and plotted as scatter dot plot of male (closed circle/square) and female (open circle/square) ± standard error of the mean. n=5-12. Aβ42 and Aβ40 levels were quantified with corresponding one-way ANOVA and paired comparison test (*, p<0.05; **, p<0.01, ***, p<0.001). (D, E) Mdk and PTN overexpression induces astrocytosis and microgliosis in 6M CRND8 mice. Representative images of GFAP-reactive astrocyte (D) and Iba-1 reactive microglia (E) in 6-month-old TgCRND8 mice overexpressing Mdk or PTN genes. Quantitation of the Iba-1 or GFAP staining was performed in cortex and hippocampus combined area. N = 6 mice. Scale bar, 500 µm, 50 µm. Clear symbols denote female mice and filled symbols denote male mice. Data represents mean ± sem. One-way ANOVA; ***p < 0.001; **p < 0.01; *p < 0.05.

**Figure S10.**
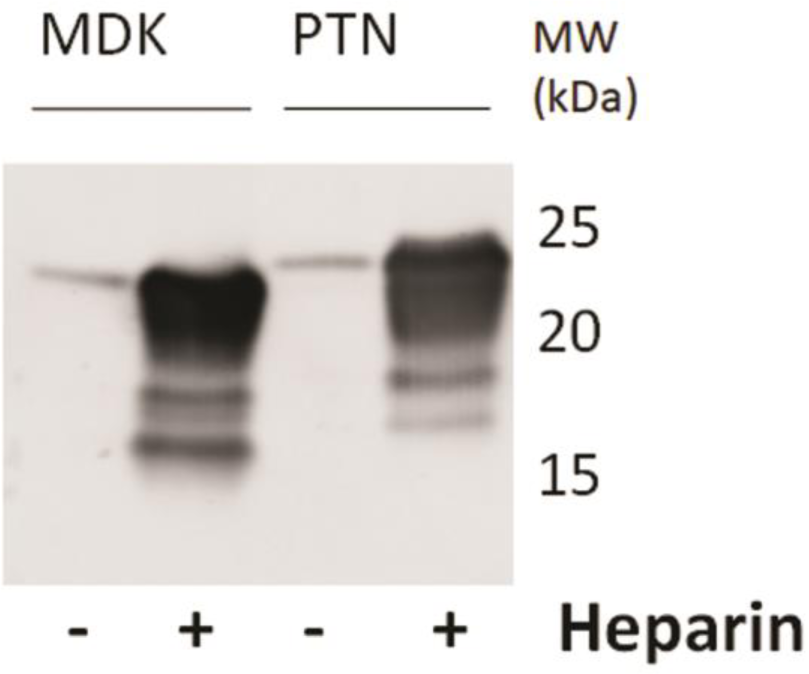
Mdk and PTN interact with heparin. Flag-tagged Mdk and PTN secreted into the media of transiently transfected HEK cells binds to heparin-agarose beads.

**Figure S11.**
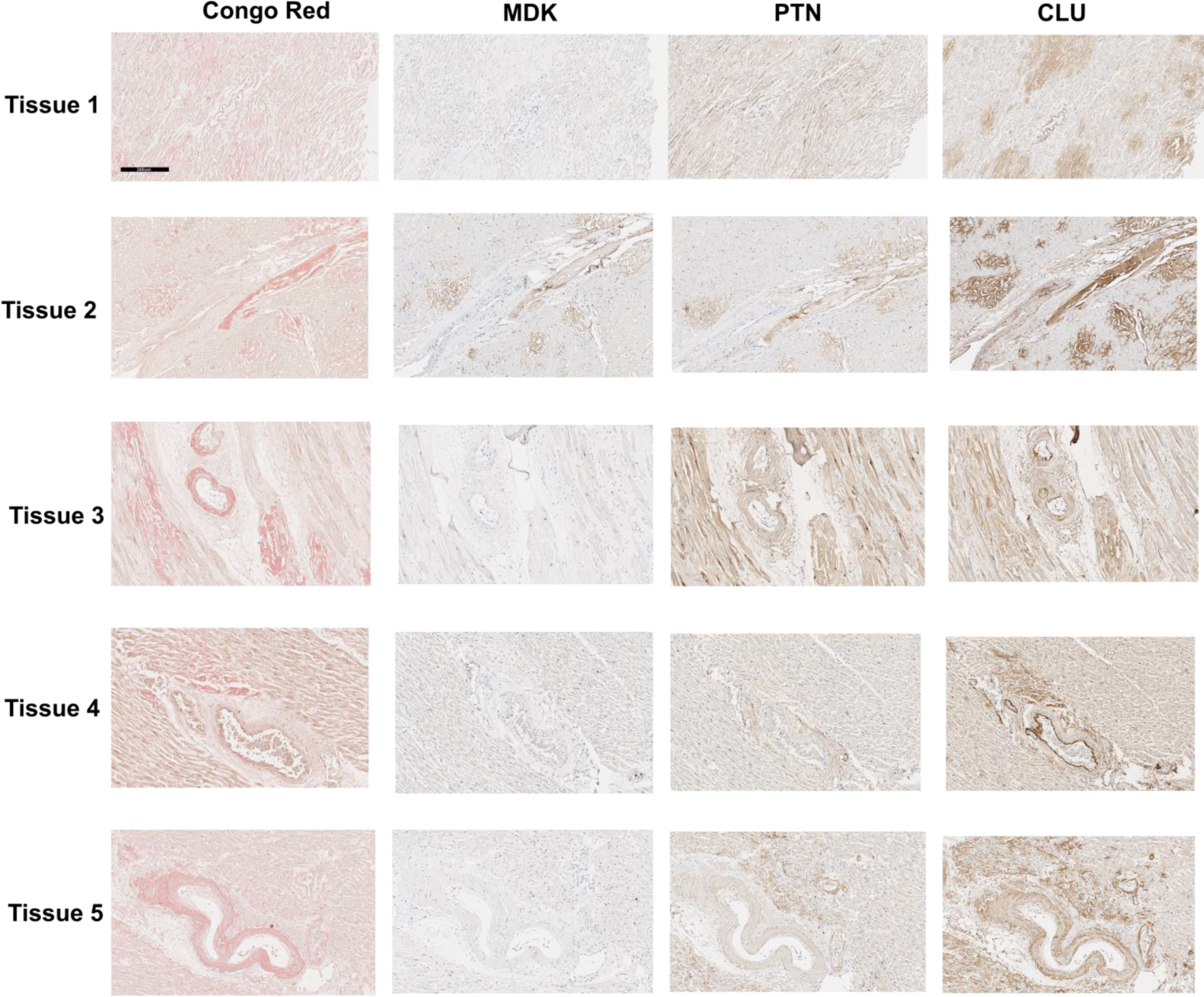
Human heart tissues stained with Congo red, MDK, PTN and CLU. Representative images of immunohistochemical stains for Congo Red, MDK, PTN or CLU on human cardiac amyloidosis sections. Scale bar, 300 µm.

**Figure S12.**
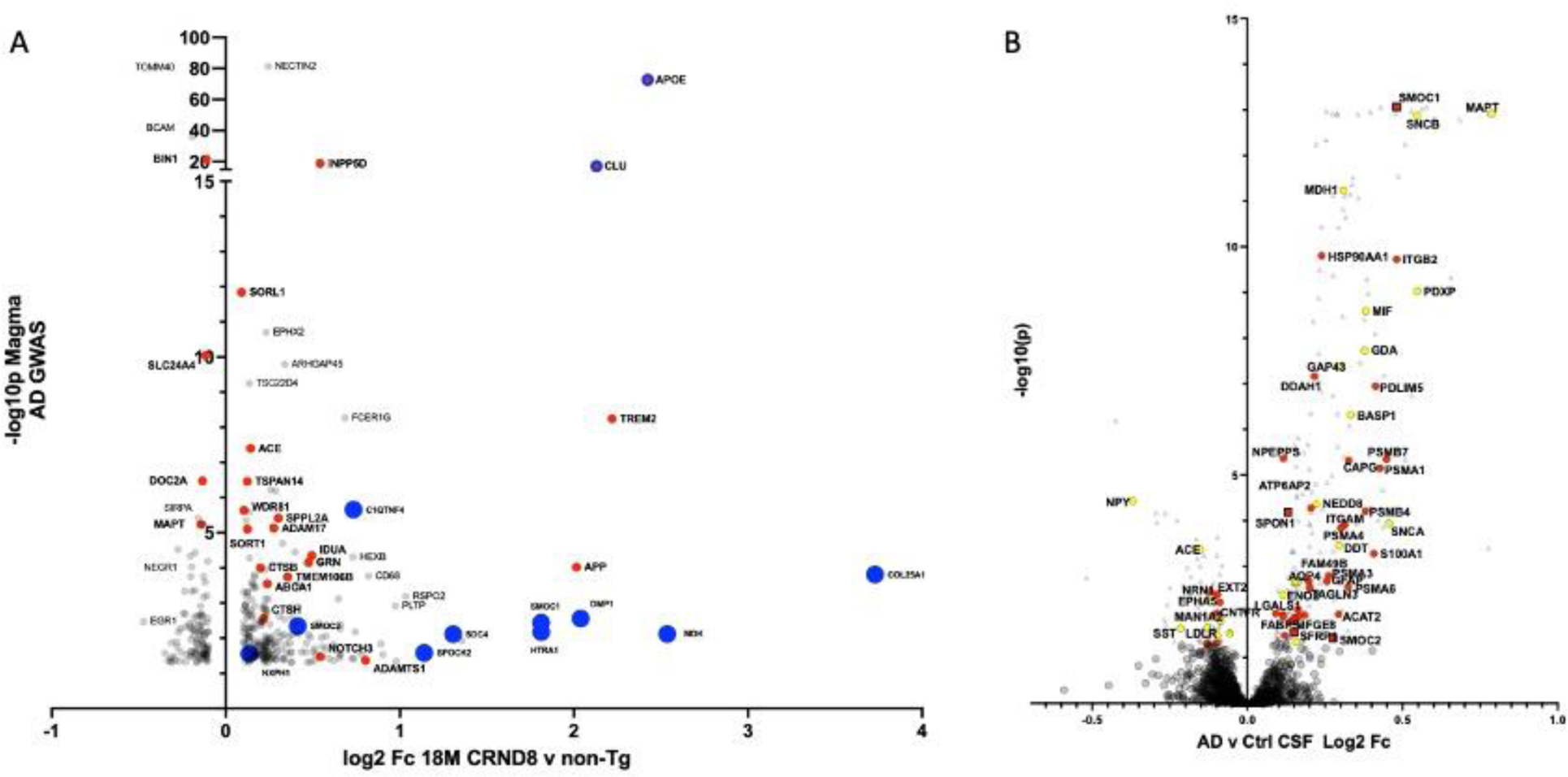
Contextualization of CRND8 18M proteome by integration with AD GWAS MAGMA p values and AD CSF proteomic changes. (A) Volcano plot of log2 Fc of DEPs in the 18M CRND8 proteome versus the −log10p MAGMA AD GWAS values (Table S1). Red circles indite GWAS loci that have been prioritized for further study. Blue circles indicate M42+ DEPs that accumulate in the CRND8 proteome. APOE and CLU are denoted as known GWAS loci that accumulate with amyloid by circles with red interior with blue border. (B) Volcano plot of the AD vs control CSF log 2 Fc derived from the Caucasian cohort in Modeste et. al 51, annotated by showing as colored symbols the overlapping DEPS in the CRND8 18M brain proteome. Red indicates a change that is concordant in direction between CSF and mouse brain. Yellow indicates a discordant change. M42 members are denoted by red squares. Grey triangles indicate significantly altered proteins in CSF that are either not detected in the 18M CRND8 proteome or not significantly changed (see Table S13).

**Figure S13.**
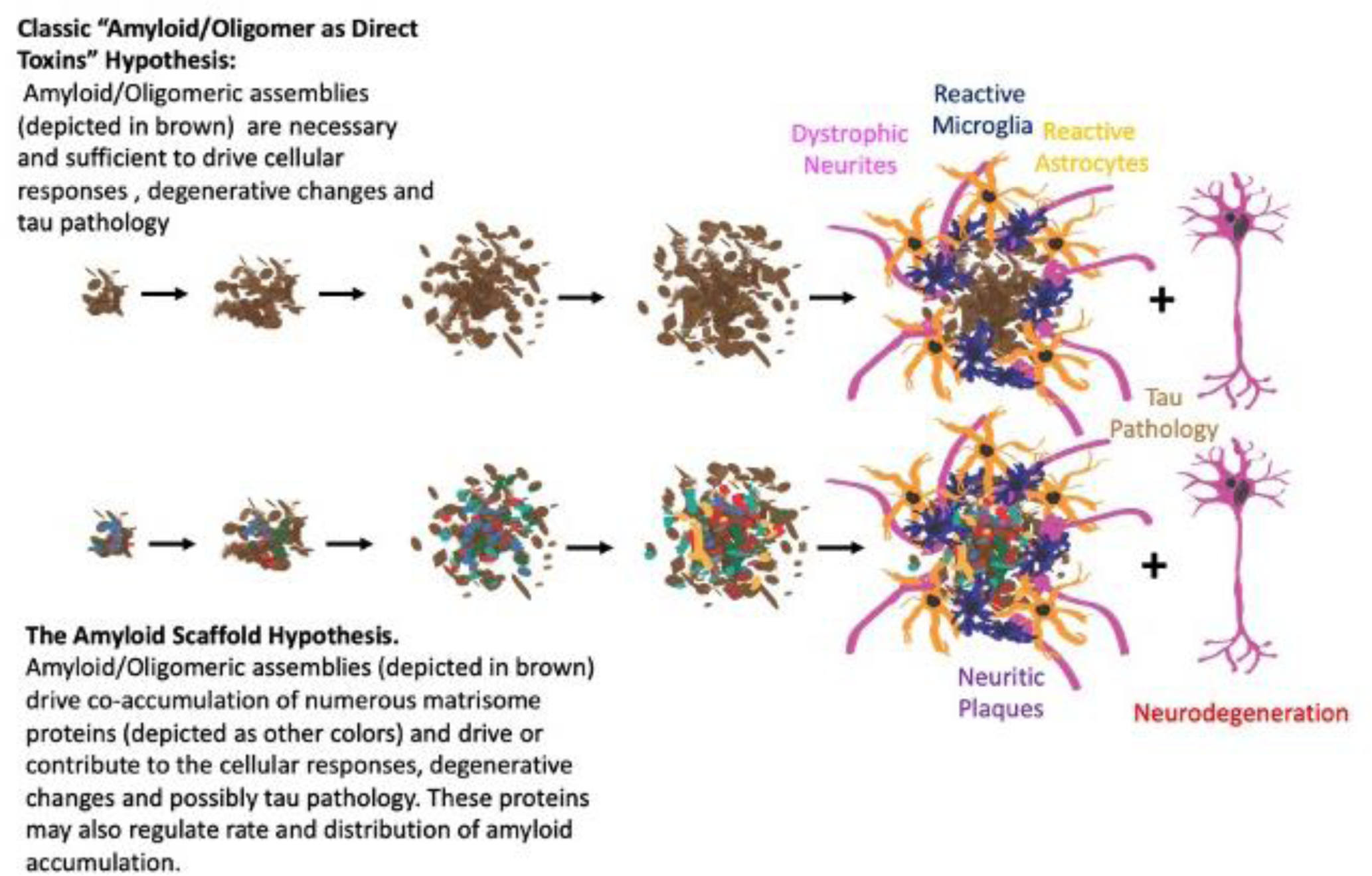
A schematic illustrating the amyloid scaffold hypothesis.

### Supplemental Tables

Table S1. Master stats table for all 3 age group batches, including MAGMA p values from AD GWAS and human brain data from Johnson et al 9.

Table S2. Unified 3 TMT batch traits

Table S3. TMT Normalized abundance

Table S4. TMT Normalized abundance 5% < FDR < 5%, Granular quantifications

Table S5. CRND8 3 age group batch statistics for proteins from table.

Table S6. Genes and mRNA expression (log2 Fc) from reprocessed AD TCx and 20M CRND8 mice used for Figure S1A.

Table S7. List of DEPS used in GO analysis in Figure 1.

Table S8. RNA-Seq Counts, Dispersion-normalized, with Statistics for 18M CRND8 vs 18M Non-Tg

Table S9. Histopathology results summary CRND8 studies.

Table S10. Antibodies used in mice and staining conditions.

Table S11. Antibodies used in humans and staining conditions.

Table S12. List of peptides used to create new antibodies and list of antibodies/antisera that were tested from commercial sources that did not work on the pathological specimens.

Table S13. List of CSF proteins and DEPs in 18M CRND8 brains used for Figure S12.

